# Polymodal sensory perception of mechanical and chemical cues drives robust settlement and metamorphosis of a marine pre-vertebrate zooplanktonic larva

**DOI:** 10.1101/2023.07.03.547492

**Authors:** Jorgen Hoyer, Kushal Kolar, Athira Athira, Meike van den Burgh, Daniel Dondorp, Zonglai Liang, Marios Chatzigeorgiou

## Abstract

The Earth’s oceans brim with an incredible diversity of microscopic planktonic animals, many of which correspond to the transient larval stage in the life cycles of benthic marine organisms. The mechanisms by which marine larvae use their nervous system to sense and process diverse environmental cues (physical and chemical) in the water column and the benthos to settle and metamorphose is a major problem across the fields of neuroscience, development, evolution and ecology, yet they remain largely unclear.

Here, we employ Ca^2+^ imaging and behavioral assays using the larval form of the protochordate *Ciona intestinalis* to characterise the mechanical and chemical stimuli these larvae respond to during settlement and metamorphosis. We also identify the polymodal sensory cells that detect these stimuli. Whole brain Ca^2+^ imaging further revealed that the presentation or removal of ethological chemosensory stimuli engages the activities of different neuronal sub-populations resulting in brain state changes, which may underlie behavioral action selections and metamorphosis. Finally, chemogenetic interrogation coupled to behavioral analysis reveals that peptidergic sensory neurons including polymodal cells capable of chemotactile perception and chemosensory/neurosecretory cells of proto-placodal ectoderm origin play a pivotal role in regulating stimulus induced settlement and metamorphosis. This work suggests that marine zooplanktonic larvae utilise their streamlined nervous systems to perform multimodal integration of ethologically physical and chemical cues to explore the oceanic water column and benthos.

## Introduction

The oceanic appearance of Eumetazoa approximately 600 Mya defines a major event in the history of life^1–4^. Today, the oceans harbour an incredible diversity of microscopic animals, many of which correspond to the transient larval stage of animals that metamorphose into larger-bodied bottom-dwelling adults. While eumetazoans exhibit a diversity of life cycles, this biphasic life cycle is the most abundant^5–7^. It is characterized by a morphologically distinct larva, which exhibits a dispersal phase in the plankton followed by a descend to the bottom of the water column where it uses its sensory apparatus to identify a suitable substrate on which to attach, settle and metamorphose into a benthic adult^5,6,8^.

To date, our understanding of how sensory modalities are employed by marine larvae to build a representation of their surroundings in pelagic environments and to identify suitable sites for settlement and metamorphosis in the benthos comes primarily from behavioral studies (reviewed in ^9–11^). The prevailing hypothesis is that nervous systems of eumetazoan larvae perceive environmental cues and integrate these with internal state changes indicative of metamorphic competence to determine whether they will settle and metamorphose^9,10^. Performing neurophysiological studies in these small marine larvae poses several challenges including for example the presence of densely packed ganglia which complicate the discrimination of individual neurons and the limited availability of established imaging methods and transgenesis tools since the vast majority are not standard laboratory models.

*C. intestinalis* a marine invertebrate chordate that belongs to the sister group of vertebrates^12,13^ and has a biphasic cycle which entails a tadpole-shaped, freely swimming larva which undergoes metamorphosis to generate the filter feeding adult ^14–18^. *C. intestinalis* larvae possess a small optically accessible nervous system suitable for functional imaging^19^, with a fully mapped synaptic connectome^20,21^, which generates a rich behavioral repertoire^22^ making them an ideal system for studying the sensory mechanisms underlying attachment, settlement, and metamorphosis of marine larvae. Previous studies have provided a detailed description of the morphogenetic steps that take place during settlement and metamorphosis over the course of a few hours^14,17,18^. It has also been shown that substrate attachment is mediated by the ascidian larval attachment organs called papillae (often referred to as palps)^14,23–31^ originating from the cranial placode^32–36^, in response to mechanical stimuli, polysaccharides and cations^24,29,37–40^. The 3D organization of *C. intestinalis* palps and their connectivity to the rest of the larval nervous system has been established using electron microscopy ^20,21,28^. Briefly, they are a triplet of protrusions at the anterior-most part of the larva, each housing four lateral primary sensory neurons (PSNs), four axial columnar cells (ACCs) and twelve central collocytes^28^. The axons of the PSNs connect to rostral trunk epidermal neurons (RTENs), which send projections to the brain vesicle^21^. Despite substantial progress over the last few years, we have a limited knowledge of the range of physical and chemical cues that can be transduced by *C. intestinalis* sensory cells, especially those involved in settlement and metamorphosis. In addition, while we benefit from the connectome of the larval brain, we have no detailed knowledge of how different settlement and metamorphosis cues are encoded in the brain and how they influence ethologically relevant behavioral actions.

Here, we identify naturally occurring and anthropogenic cues that can promote or inhibit settlement behavior and metamorphosis in *C. intestinalis* larvae. We show that at least a subset of these chemical cues can be sensed by the palp residing PSNs and ACCs, which are polymodal sensory cells, since they can also respond to qualitatively distinct types of mechanical stimuli. In addition, we report on another cell type, the non-palp epidermal neurons called ATENs, which are also able to sense chemical stimuli. Using whole brain calcium imaging we show that the presentation and removal of these chemical cues drastically alters global brain dynamics. The coordinated, nervous system wide activity in response to chemical stimulation, likely plays an important role in the settlement and metamorphosis behaviors. This view is corroborated by a series of chemogenetic silencing experiments.

## Results

### Natural and anthropogenic cues can promote or impede settlement and the first phase of metamorphosis, the process of tail regression

The identification of a suitable site for attachment, settlement, and metamorphosis of marine larvae depends highly on mechanical^40–44^ as well as long and short range chemical cues^9,45,46^ (Fig. 1A).

**Fig. 1.**
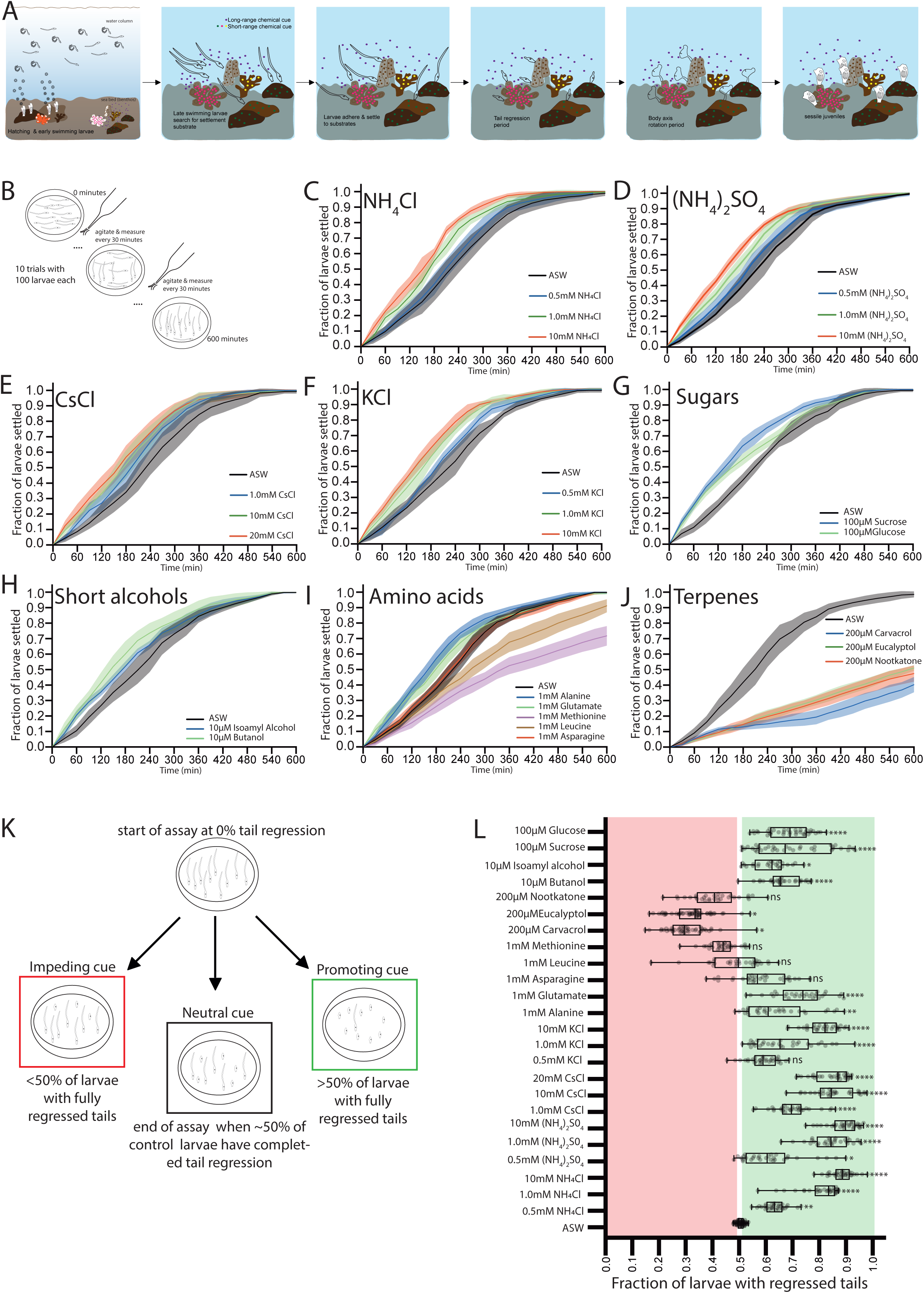
Chemosensory cues promote or impede settlement and tail regression of *C. intestinalis* larvae. (A) Life cycle of *C. intestinalis*: early and mid-stage larvae swim in the water column; late-stage larvae employ mechanical and chemical cues to identify a suitable site in the benthos for settlement and metamorphosis. Successful completion of metamorphosis results in a new generation of juveniles. (B) Schematic of the settlement assay. (C-J) Settlement curves for Artificial Seawater (ASW) versus different chemicals at indicated concentrations specified in each plot. Solid lines plot mean; shaded areas show standard deviation (SD). Statistics: two-way RM ANOVA followed by Dunnett’s Multiple comparison test (Table S1-S8). See table S10 for timepoints at which 50% of larvae have settled for a given stimulus. (K) Schematic of the tail regression assay. The end point of the assay was set when 50% of control (ASW) larvae have fully regressed tails. Number replicates (plates) per cue: ≥30. (L) Box plots show median, quantiles of the fraction of larvae that have completed tail regression. Individual grey circles correspond to each replicate. Statistics: Kruskal-Wallis test followed by Dunn’s multiple comparisons test (table S9).

To determine which chemical cues, promote or impede settlement and metamorphosis, we performed two types of behavioral assays to test a wide panel of stimuli, which have been previously implicated in settlement and metamorphosis across metazoans^9,37–39,45–48^ as well as cues that are known to be contaminants of human-made chemicals or activities that may act as anthropogenic effectors of settlement and metamorphosis^49,50^.

In the first assay we quantified the fraction of larvae that had adhered to the plastic dish bottom at 30-minute intervals (Fig. 1B). We found that cations NH_4_^+^, Cs^+^, K^+^ at concentrations ≥ 1mM significantly accelerated larval settlement in a concentration dependent manner (Fig. 1C-F, Tables S1-S4, S10). Short alcohols which are known to trigger behavioral responses, including settlement in marine organisms^51,52^ and sugars which are degradation products of the polysaccharides contained in biofilms, eelgrass, seagrass or algae also resulted in significantly faster settlement rates (Fig. 1G, H, Tables S5, S6, S10). Amino acids are also relevant aquatic chemical cues for numerous marine organisms^53–56^. We found that Alanine and Glutamate significantly accelerated settlement (Fig. 1I, table S7, S10). Two other amino acids Leucine and Methionine decreased settlement rates with the difference relative to controls becoming significant at later stages of the assay (Fig. 1I, table S7, S10). Terpenes are amongst the most abundant class of natural products in terrestrial and marine environments^57–59^. We found that three terpenoids significantly inhibited settlement (Fig. 1J, table S8, S10), suggesting that they have a negative settlement valence.

We went on to measure the effect of these chemicals on metamorphosis. We focused our analysis on the fraction of settled larvae that had completed the first step of metamorphosis, which is tail regression. To be able to identify in the same assay chemicals that either promote or impede larval tail regression we used as our endpoint the timepoint when 50% of larvae in Artificial Sea Water ASW (the control group) had regressed their tails (Fig. 1K). We found that NH_4_^+^, Cs^+^, K^+^ promote metamorphosis in a concentration dependent manner (Fig. 1L, table S9). Short alcohols, sugars, Alanine and Glutamate resulted in a significantly higher fraction of metamorphosed larvae (Fig. 1L, table S9). In contrast the terpenes eucalyptol and carvacrol significantly impeded metamorphosis (Fig. 1L, Tables S9). The terpene nootkatone and the amino acid methionine also showed a modest but not significant reduction in metamorphosis (Fig. 1L, table S9). Our data suggest that in *C. intestinalis,* the settlement behavior and the first step of metamorphosis process, namely tail regression, mostly share common promoting and impeding sensory cues.

We also tested noxious concentrations of NH_4_Cl (100mM), copper (1mM), chloroquine (10mM) and the detergent SDS (0.1%), which prevented settlement and caused larval mortality after 120 minutes of exposure (fig. S1A, table S11). We were not able to quantify metamorphosis assays since prolonged exposure to these cues resulted in death, which is in line with existing evidence, showing that high concentrations of ammonia, copper and detergents are toxic for marine organisms^49,60^.

### The PSNs are polymodal sensory neurons responding to mechanical, soluble and poorly soluble chemical cues

To monitor the activity of the PSNs upon exposure to strongly or weakly soluble attractants, repellents, and other stimuli we used the calcium indicator GCaMP6s. We generated transient transgenics under the control of the *proprotein convertase 2* (*pc2*) promoter, which expresses in the PSNs but not the ACCs (labelled by the *βγ-crystallin* promoter^61^) (Fig. 2A, B) to monitor Ca^2+^ activity. Transgenic animals were immobilized using a holding pipette and placed in a small perfusion chamber under constant flow of Artificial Sea Water. To deliver precise mechanical stimuli we used a piezo-electric driven glass pipette (Fig. 2C). We used a perfusion pencil with millisecond-latency solenoid valve control for olfactory and gustatory chemical cues (Fig. 2D). Robust responses for both mechanical and chemical stimuli were observed in the PSN cell body, as well as in the PSN apical protrusions (Fig. 2E, F). Perfusion with just ASW did not yield Ca^2+^ transients in the PSNs (Fig. 2G, P; fig. S2H). However, the PSNs responded with robust Ca^2+^ transients in a concentration dependent manner to 1mM, 10mM and 100mM NH_4_Cl summarized in Fig. 2H as average normalized fluorescence change (ΔF/F_0_) versus time (Fig. 2H, P; fig. S2H). The PSNs also responded to CsCl (Fig. 2I, P; fig. S2H), the weakly soluble terpenes Eucalyptol and Carvacrol (Fig. 2J, K, P; fig. S2H), the short alcohol Butanol (Fig. 2L, P; fig. S2H). In addition, we recorded smaller responses to the alkaloid repellent chloroquine and the amino acid Glutamate but not in response to Asparagine (Fig. 2M, N, P; fig. S2A, S2H). We observed Ca^2+^ transients upon stimulation with the detergent SDS (Fig. 2O, P; fig. S2H). Upon presenting the larvae with a hyperosmotic stimulus (1M Glycerol) we recorded a small decrease in GCaMP signal of the PSNs relative to their resting activity levels, followed by robust stimulus OFF Ca^2+^ transients (Fig. 2P; fig. S2B, S2H). No motion artifacts were observed in controls where *pc2>GFP* transgenic animals were stimulated with either ASW or 1mM NH_4_Cl (fig. S2C, D, G, H).

**Fig. 2.**
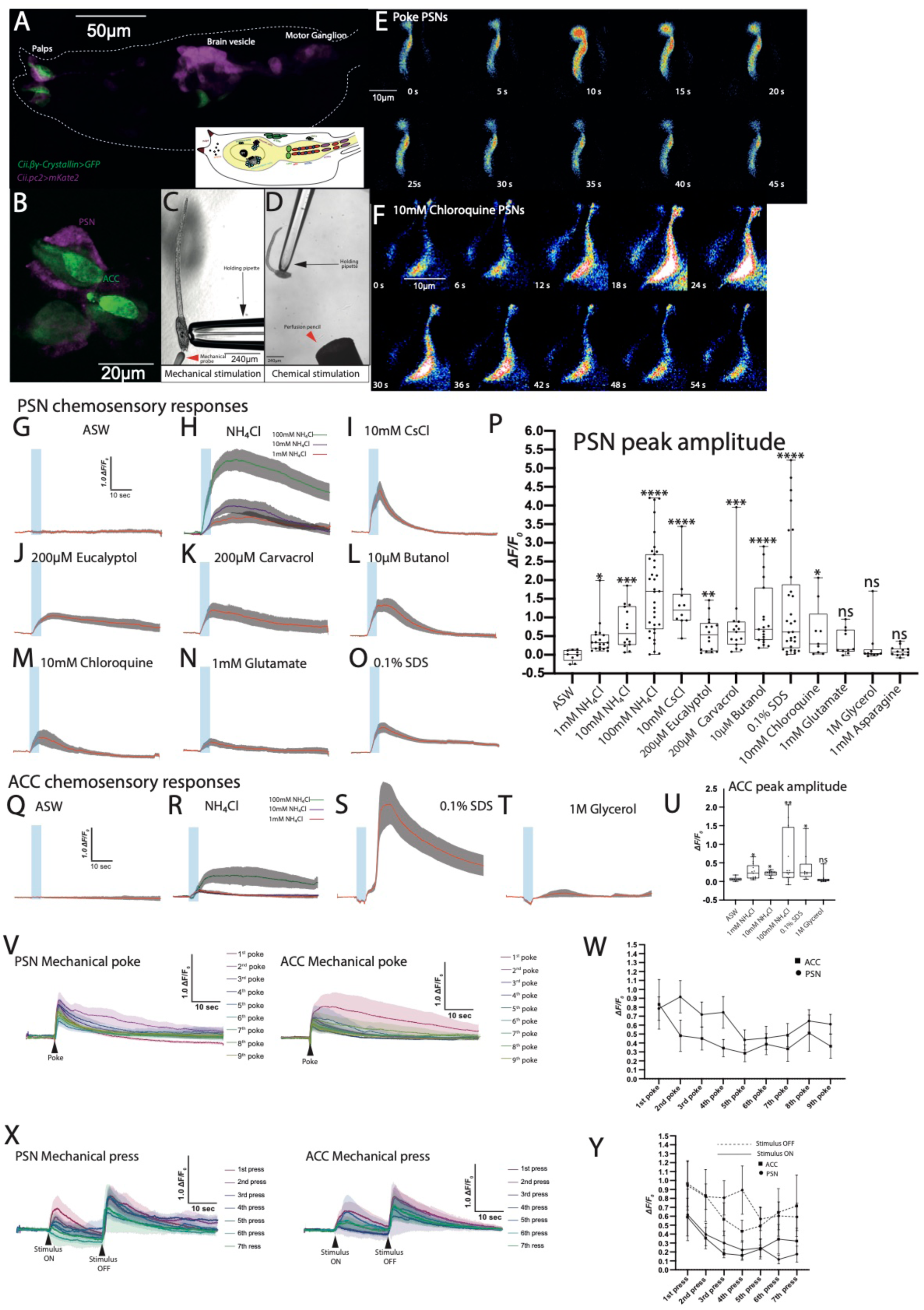
The PSNs and ACCs respond to diverse mechanical and chemical cues. (A) Larva expressing *Cii-βγ-Crystallin>GFP* (green) and *Cii-pc2>mKate2* (magenta). Inset: schematic of larval brain highlighting several neuronal cell types. (B) *Cii-βγ-Crystallin>GFP* (ACCs) and *Cii-pc2>mKate2* (PSNs) label distinct cell populations in the palps. (C, D) Experimental set-ups for mechanical and chemical stimulations. (E, F) PSN Ca^2+^ responses to mechanical (Movie S1) and chemical (Movie S2) stimulations. (G-O) Average PSN Ca^2+^ responses to chemical stimuli. Solid lines: means; Shaded areas: standard error of the mean (SEM); Blue bar: stimulus presentation period. (P) Quantification of PSN Ca^2+^ peak amplitudes across different chemical stimuli. (Q-T) Average ACC Ca^2+^ responses to chemical stimuli. Solid lines: means; Shaded areas: standard error of the mean (SEM); Blue bar: stimulus presentation period. (U) Quantification of ACC Ca^2+^ peak amplitudes. (V) PSN and ACC Ca^2+^ responses to 9 consecutive mechanical pokes. Black arrowhead: poke application; error bands: S.E.M. Statistics: Kruskal-Wallis tests, followed by Dunn’ test compared to ASW Controls (tables S13, S14). (W) Ca^2+^ peak amplitudes of PSNs (black circles) and ACCs (black squares) to 9 consecutive mechanical pokes. (X) PSN and ACC Ca^2+^ responses to 7 consecutive mechanical presses. Black arrowheads: press On and OFF; error bands: S.E.M. (Y) Ca^2+^ peak amplitudes of PSNs (black circles) and ACCs (black squares) in response to 7 consecutive mechanical presses. (G-Y) See Table S12 for number of cells contributing to each average trace and boxplot.

We then asked the question of whether the ACCs are also able to respond to some of the same chemical cues as the PSNs. We expressed GCaMP6s under the control of the ACC specific promoter *βγ-crystallin* and we imaged using the same approach as for the PSNs. The ACCs did not respond to control ASW flow (Fig. 2 Q, R; fig. S2I) while *βγ-crystallin*>GFP also showed no response to ASW or 1mM NH_4_Cl (fig. S2E-G, I). We found that the ACCs responded to multiple concentrations of NH_4_Cl. The responses to 1mM and 10mM NH_4_Cl were of similar magnitude, while presentation of 100mM NH_4_Cl yielded larger and more sustained responses (Fig. 2R, U; fig. S2I). We also obtained stimulus ON responses to the detergent SDS, however a stronger Ca^2+^ signal was observed when we removed the SDS stimulus (stimulus OFF response) (Fig. 2S, U fig. S2I). We found that the ACCs responded weakly to an osmotic stimulus (1M Glycerol) and in sharp contrast to the PSN osmotic stimulus responses they did not exhibit a stimulus OFF response (Fig. 2T, U; fig. S2I).

Sensory cells in the palps are important for sensing the mechanical properties of settlement substrates^29,31,40^. We observed Ca^2+^ transients in the PSNs in response to both poke (short stimulus) (Fig.2V, W; fig. S3A) and press stimulation (longer stimulus) (Fig. 2X, Y; fig. S3C). Interestingly, PSN Ca^2+^ responses to mechanical press are characterized by the presence of clearly discernible stimulus-on and stimulus-off responses (Fig. 2X; fig. S3C). In contrast, with mechanical pokes stimulus-on and stimulus-off responses cannot be characterized due to the instantaneous nature of the stimulus (Fig. 2V; fig. S3A). We tested the ability of the PSNs to adapt to repeated poke or press stimulations. We found that PSNs showed little adaptation to as many as nine poke repetitions (Fig. 2V, W black circle datapoints, fig. S3A). In contrast, the PSNs showed evidence of adaptation after only three rounds of press stimulation (Fig. 2X, Y black circle datapoints, fig. S3C). The different Ca^2+^ transient and adaptation profiles suggest that the PSNs respond in a distinct manner to different forms of mechanical stimuli, possibly by leveraging multiple types of mechanosensitive ion channels.

We then tested the ability of the ACCs to respond to mechanical stimuli. We found that they responded to both mechanical poke and mechanical press stimuli (Fig. 2V-Y; fig. S3B, D). The mechanical press responses in ACCs were characterized by a stimulus-on response, followed by a larger in magnitude stimulus-off response (Fig. 2X, Y black square data points; fig. S3D). We tested the ability of the ACCs to adapt to repeated poke or press stimulations. We found that the ACCs showed strong adaptation already from the second poke trial, which persisted for the additional seven rounds of poking (Fig. 2V, W black square data points; fig. S3B). In contrast, the ACCs did not show a robust adaptation to press stimulation (Fig. 2X, Y, 2AA black square data points; fig. S3D). While we observed a decrease in Ca^2+^ peak amplitude between the 2^nd^ and 4^th^ press stimulation, the ACCs Ca^2+^ responses were able to partially recover in the subsequent 3 trials (Fig. 2X, Y black square data points; fig. S3D).

### Presentation and removal of settlement cues alter brain-wide activity

We then wondered how chemosensory information is processed upstream of the PSNs and the ACCs. We performed whole-brain single-cell-resolution Ca^2+^ imaging with a pan-neuronally expressed nuclear localized Ca^2+^ sensor (GCaMP6s), which permitted the unambiguous discrimination of individual neurons within the densely packed brain of *C. intestinalis* larvae. To discern neuronal outlines, we used a plasma membrane marker (Fig. 3A, B). Animals were immobilized under a spinning disk microscope. We presented six chemical stimuli (1mM NH_4_Cl, 100mM NH_4_Cl, 10mM CsCl, 200μM Carvacrol, 0.1% SDS and 10μM Butanol) and recorded neuronal activity prior, during and after stimulus presentation. The imaging volume spanned the entire trunk, which includes most of the larval sensory neurons, interneurons as well as motor neurons. In each recording we detected approximately 100 neurons, however only a subset of these neurons showed strong activity during the recordings. Examples of neurons that showed strong activity during imaging include the AMG secondary interneurons the peripheral relay neurons (PNRNs) and the motorneurons (MN) (Fig. 3C-E). These observations are in line with the predicted mechano-chemosensory pathways predicted by the connectome^20^. To better scrutinize the relationships between the neuronal signatures and population dynamics before, during and after chemosensory stimulation we constructed a low-dimensional representation of the neural activity of *C. intestinalis* by performing principal components analysis (PCA) on the z-scored Ca^2+^ activity. The neural trajectories showed the brain state evolution over time, across different animals stimulated with the chemical cues (Fig. 3F-K). For all stimuli tested the larvae explored more of the neuronal dynamics space during or after stimulation compared to the pre-stimulus phase. For chemical cues which strongly impede settlement (200μM Carvacrol) (Fig. 3F; fig. S4A,B) or are noxious (100mM NH_4_Cl and 0.1% SDS) (Fig. 3G,H; fig. S4C-F) we observed that the end points of the neuronal dynamics trajectories ended far away from the starting point. For cues which are associated with promoting settlement we observed that the neuronal dynamics evolved in circular trajectories in the PCA space, with the starting and ending points of the trajectories being close to each other in PCA space (Fig. 3I-K; fig. S4G-L).

**Fig. 3.**
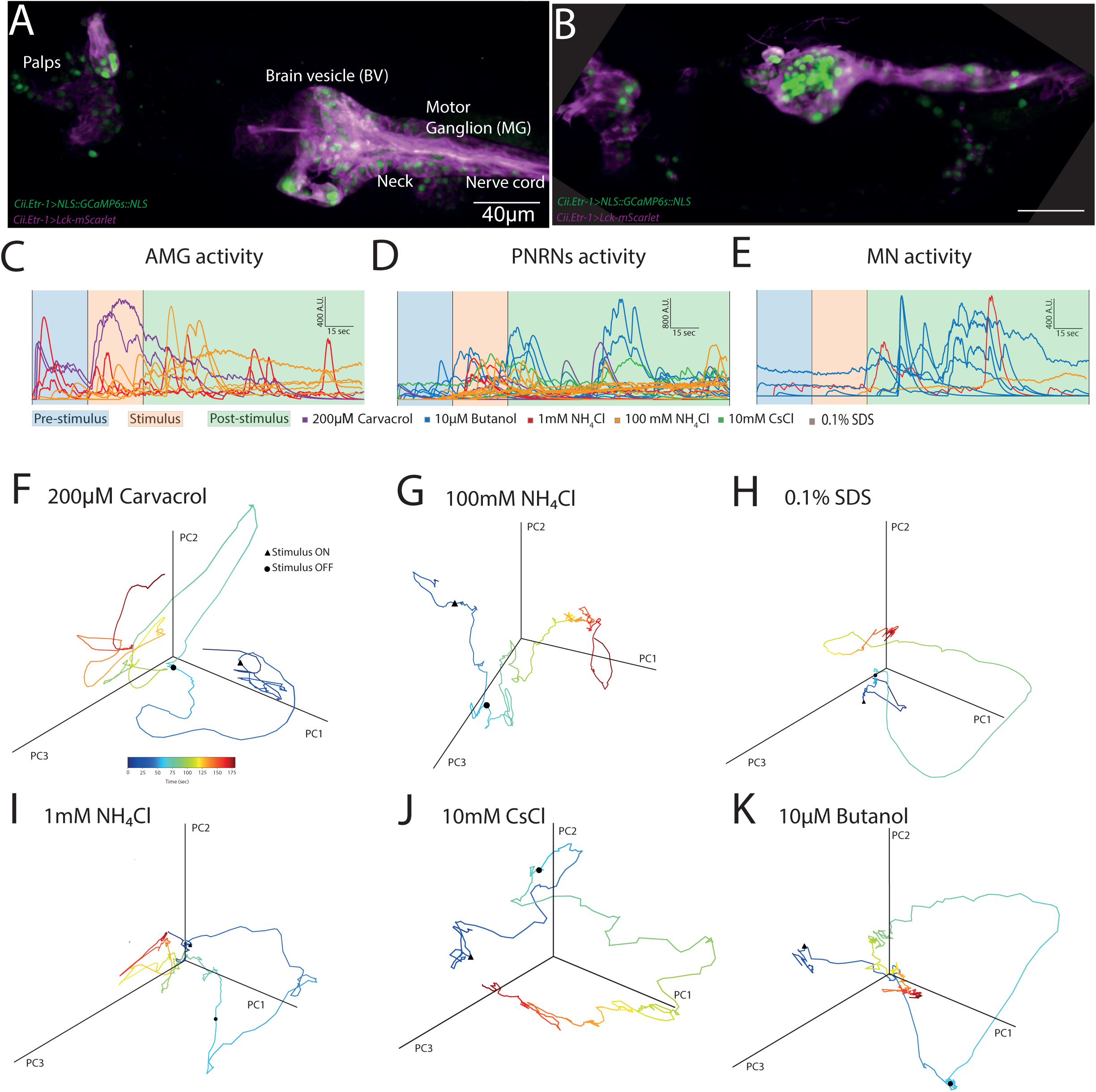
Whole-brain imaging reveals brain wide state changes in response to chemical stimulation. (A) Maximal projection confocal micrograph of a larval trunk (dorsal view). The nervous system of the larva is labelled with *Cii.Etr-1>nls::GCaMP6s::nls* (green nuclei) and *Cii.Etr-1>Lck-mScarlet* (magenta plasma membrane/outline of neurons). Different regions of the nervous system are labelled. (B) Lateral view of the nervous system of a *C. intestinalis* larva. The transgenes expressed are the same as in the panel A larva. (C-E) Examples of Ca^2+^ activity from different groups of neurons in the larval brain in response to chemical stimuli. These include the (C) ascending motor ganglion interneurons (AMGs), (D) PNS relay neurons (PNRN), (E) motorneurons (MG), (F-K) Phase plots of PC 1-3 showing the brain state’s evolution across different animals stimulated with (F) 200μM Carvacrol, (G) 100mM NH_4_Cl, (H) 0.1% SDS, (I) 1mM NH_4_Cl, (J) 10mM CsCl and (K) 10μM Butanol. For each phase plot the stimulus onset (black triangle) and stimulus end (black circle) are indicated. Temporal progression is indicated by a change in the trajectory line colour from dark blue to dark red (see also colorbar in panel F).

While analysing our whole-brain imaging data we noticed that two neuronal subpopulations termed apical trunk epidermal neurons (ATENs) were showing robust responses in the presence and/or upon the removal of the chemical stimuli (Fig. 4A-J). The ATENs are hypothesized to resemble an ancestral cell type with a dual neurosecretory and chemosensory function^62^. To characterize the responses of the ATENs to the sensory stimuli we generated tuning curves (Fig. 4K i-vi). We found that for most of the 10μM Butanol and 100mM NH_4_Cl trials the tuning curve peak was during the stimulus period (Fig. 4 K ii, v). Tuning of the ATENs in trails with 200μM Carvacrol, 10mM CsCl and 1mM NH_4_Cl was more variable (Fig. 4K i, iii, iv), suggesting that the ATENs are indeed sensory cells capable of responding to both soluble and poorly soluble chemical cues that may be used by the larvae to evaluate the suitability of a substrate for settlement and metamorphosis. In contrast the tuning curve peak of the ATENs in response to 0.1%SDS was during the stimulus-OFF period (Fig. 4K vi). To better evaluate the influence of the ATENs’ responses to whole brain population dynamics, we performed PCA analysis and generated phase plots for additional whole brain imaging movies (Fig. 4L i-vi). We found that similarly to other recordings, sensory stimulation of the larval nervous system with chemical cues induced a brain state change as evident from the evolution of the neural dynamics trajectory in the PCA space (Fig. 4L i-vi). In some cases, the ATENs mark inflection points along the brain state trajectories suggesting that they can influence the evolution of the brain state dynamics.

**Fig. 4.**
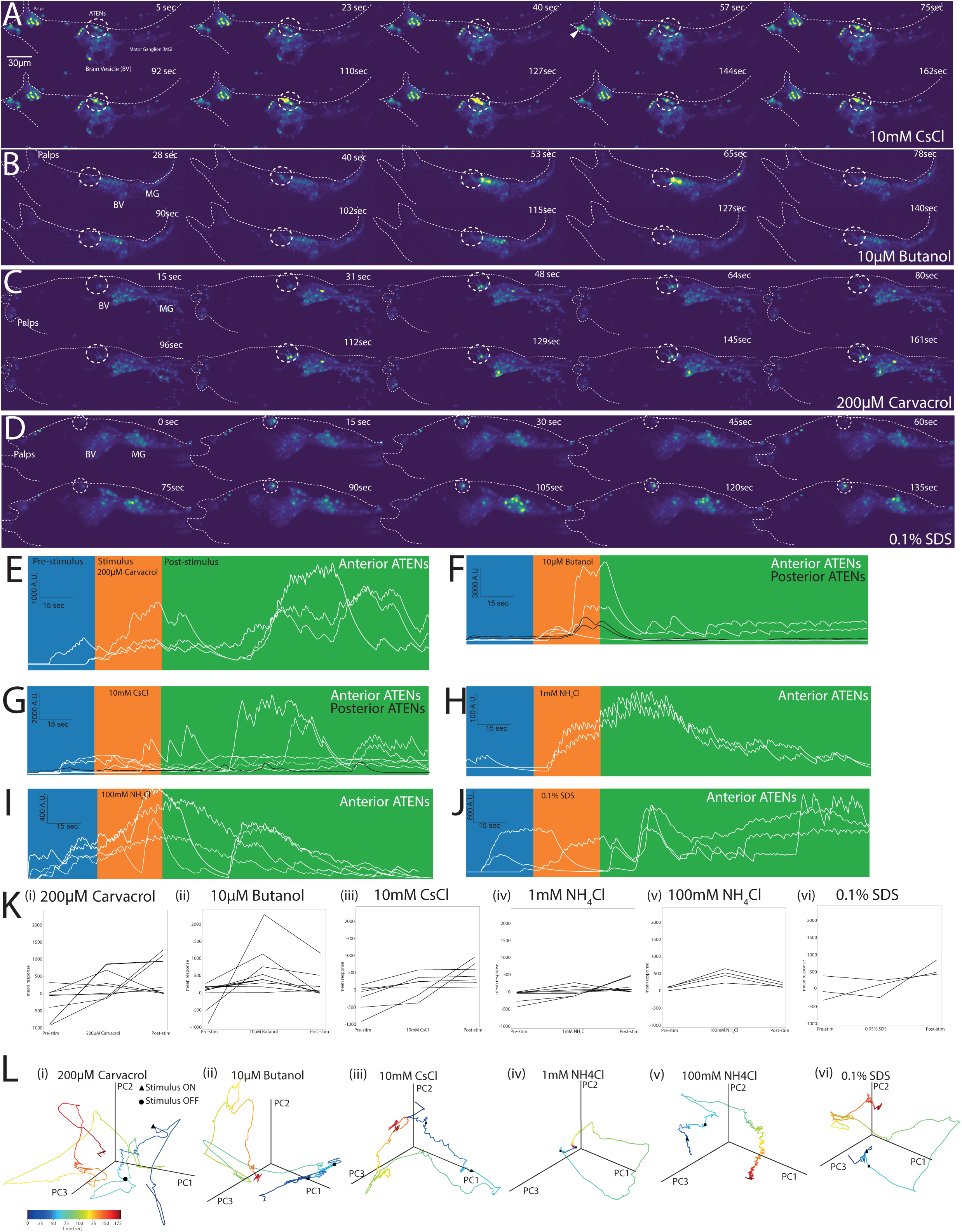
ATENs are chemosensory neurons responding to a variety of settlement cues. (A-D) Montage of different whole-brain Ca^2+^ imaging recordings using *Cii.Etr-1>nls::GCaMP6s::nls* in response to (A) 10mM CsCl (S3 Movie), (B) 10μM Butanol (Movie S4), (C) 200μM Carvacrol (Movie S5), (D) 0.1% SDS (Movie S6). ATENs in all panels are indicated using a thick dashed line circle. (E-J) Example traces of Ca^2+^ activity in the Anterior ATENs (white traces) and Posterior ATENs (black traces) in response to (E) 200μM Carvacrol, (F) 10μM Butanol, (G) 10mM CsCl, (H) 1mM NH_4_Cl, (I) 100mM NH_4_Cl and (J) 0.1% SDS. (K) Panels (i-vi) show the tunning curves for (i) 200μM Carvacrol, (ii) 10μM Butanol, (iii) 10mM CsCl, (iv) 1mM NH_4_Cl, (v) 100mM NH_4_Cl and (vi) 0.1% SDS. (L) Phase plots of PC 1-3 showing the brain state’s evolution across different animals stimulated with (i) 200μM Carvacrol, (ii) 10μM Butanol, (iii) 10mM CsCl, (iv) 1mM NH_4_Cl, (v) 100mM NH_4_Cl and (vi) 0.1% SDS. For each phase plot the stimulus onset (black triangle) and stimulus end (black circle) are indicated. Temporal progression is indicated by a change in the trajectory line colour from dark blue to dark red (see also colorbar in panel L).

We also examined the PCA loadings in each case to examine which cells have the largest effect on each of the principal components. We found that, in cases where most of the cells strongly respond to the presentation of the stimulus, for example 0.1% SDS and 1mM NH4Cl (fig. S4 E, G; fig. S5 E, G) and the primary principal component PC1 has positive loadings values corresponding to these cells (fig. S4 F, H, fig. S5 F, H). This suggests that in these cases, PC1 encodes the activity during the stimulus-on period. The sparse activity during pre-stimulus periods in 0.1% SDS is encoded by PC2 with strong negative PCA loadings in the corresponding cells whereas PC3 encodes the activity profile in the post-stimulus period (fig. S4 F, fig. S5F).

### Chemogenetic perturbations demonstrate the importance of neuropeptidergic and dopaminergic cells in mediating stimulus-dependent settlement and tail regression

To further investigate the neuronal underpinnings of settlement and metamorphosis we used chemogenetic perturbations^63,64^ to silence specific neuronal populations in a temporally specified manner at the onset of settlement or metamorphosis. To determine the effects of silencing the ACCs and a range of sensory neurons including the PSNs and the ATENs we expressed the chemogenetic tool hM4D and under the *βγ-crystallin* and *pc2* promoters respectively. We first tested whether clozapine-N-oxide (CNO) acts as a settlement and metamorphosis cue in the context of our assays by exposing pc2>GFP transgenic larvae to either ASW or 10µM CNO. We quantified the settlement rates and fraction of larvae with regressed tails using pc2>GFP (+) larvae and we found no significant difference between the two conditions (fig. S6A, B, Tables S15, S16, S20). This suggests that CNO does not act as an inductive or impeding cue. We then asked the question of what happens to settlement rates in the absence of a chemical stimulus (Fig. 5A). In transgenic larval populations without the exogenous actuator clozapine-N-oxide (CNO) we found that settlement rates were comparable to chorionated controls presented in Fig. 1 (Fig. 5A, table S17, S20). In *βγ-crystallin>hM4D* or *pc2>hM4D* single transgenic larvae and *βγ-crystallin>hM4D; pc2>hM4D* double transgenic larvae where CNO was added we observed a delay in settlement rates suggesting that functional ACCs and peptidergic sensory neurons including the PSNs are required for sensing the mechanical settlement substrate (Fig. 5A, table S17, S20). We then tested the effects of silencing these cell populations on the settlement rates in the presence of cues that either promote (10mM NH_4_Cl) (Fig. 5B, table S18, S20) or impede (200μM Carvacrol) settlement (Fig. 5C, table S19). In the presence of 10mM NH_4_Cl and CNO, *βγ-crystallin>hM4D* or *pc2>hM4D* single transgenic and double transgenic *βγ-crystallin>hM4D; pc2>hM4D* larvae where neuronal activity was silenced, both single and double transgenics exhibited reduced settlement rates relative to CNO(-) controls, suggesting that NH_4_Cl mediated settlement is mediated by the ACCs and peptidergic neurons including the PSNs (Fig. 5B, table S18, S20). In the presence of 200μM Carvacrol, *βγ-crystallin>hM4D* and *pc2>hM4D* single transgenic and double transgenic *βγ-crystallin>hM4D; pc2>hM4D* larvae in the absence of CNO showed a strong impedance of settlement rates (Fig. 5C, table S19). In the presence of CNO single and double transgenic larvae exhibited higher settlement rates suggesting that silencing of the ACCs and peptidergic neurons including the PSNs alleviated the settlement this impedance was induced by the presence of Carvacrol (Fig. 5C, table S19).

**Figure 5.**
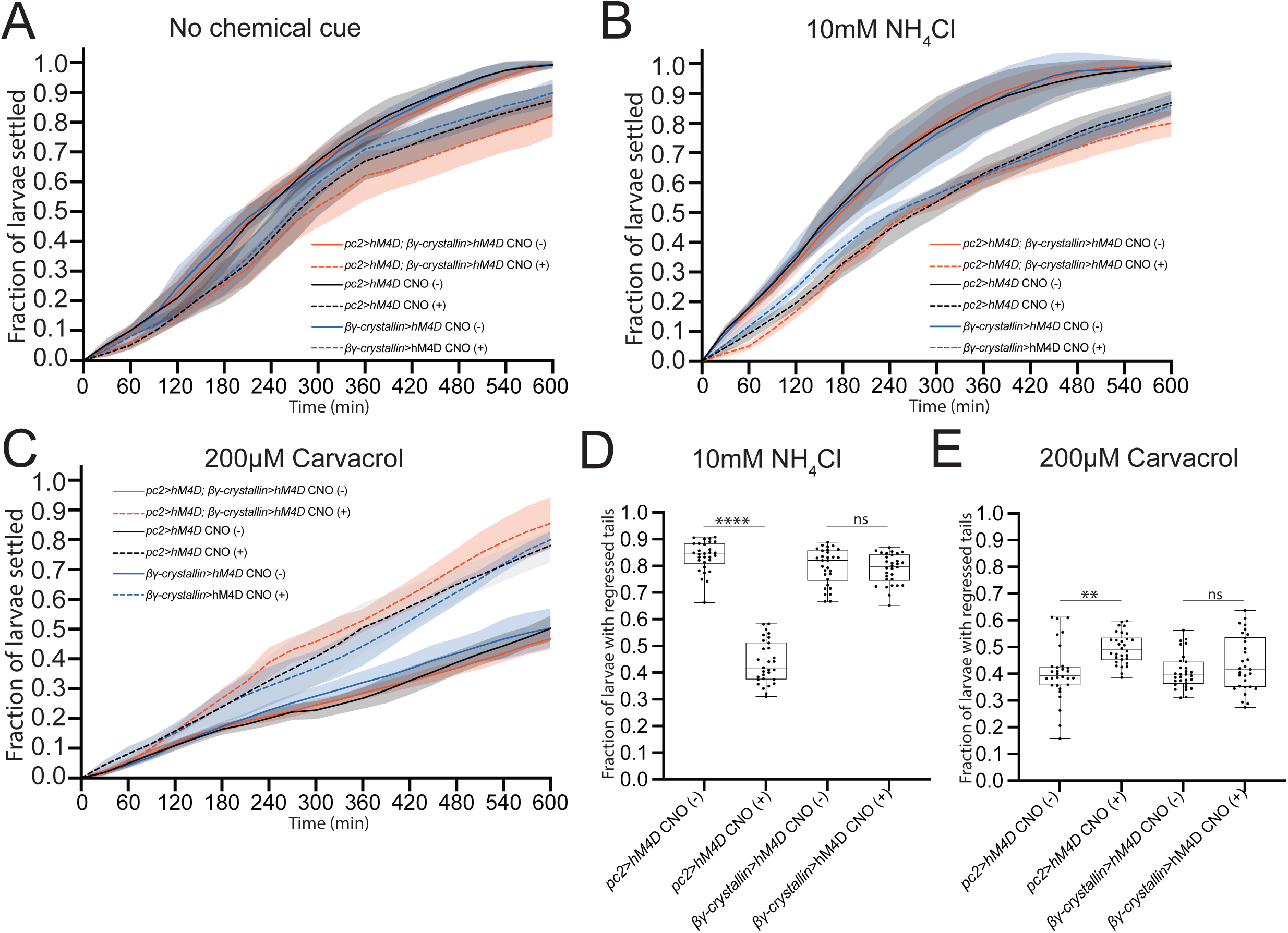
ACCs are essential only for settlement, while peptidergic neurons including the PSNs and ATENs are required for both settlement and tail regression. (A) Larval settlement curves in the absence of a chemical cue for transgenic animals expressing the chemogenetic tool hM4D in the ACCs (blue), peptidergic neurons including PSNs and ATENs (black) and in both ACCs and peptidergic neurons (red). Solid lines: CNO(-), dashed lines CNO(+). Solid lines plot mean; shaded areas show SD. (B) Larval settlement curves in the presence of 10mM NH_4_Cl. (C) Larval settlement curves in the presence of 200μM Carvacrol. For each genotype we assayed a total of 1000 larvae in 10 independent replicates. Statistics: for panels A-C we used a Two-way RM ANOVA test, followed by Tukey’s multiple comparisons test (tables S17-S19) (D,E) Box plots showing the fraction of larvae with fully regressed tails, carrying either the *pc2>hM4D* or *βγ-crystallin>hM4D* in the presence (+) or absence (-) of CNO and either 10mM NH_4_Cl (D) or 200μM Carvacrol (E). Statistics: for panels D and E we used a Kruskal-Wallis test, followed by Dunn’s multiple comparisons test (tables S21, S22).

We went on to investigate the effect of silencing or activating these sensory neuron populations on the first step of metamorphosis, tail regression. We allowed *βγ-crystallin>hM4D* and *pc2>hM4D* single transgenic larvae to settle and we then measure the fraction of larvae with fully regressed tails in the absence or presence of CNO in combination with either 10mM NH_4_Cl or 200μM Carvacrol. We found that silencing or activating the ACCs had no significant effect in the rates of tail regression (Fig. 5D, E, Tables S21, S22) indicating that the ACCs are dispensable following settlement. We found that silencing the pc2(+) cells in the presence of 10mM NH_4_Cl or 200μM Carvacrol, altered the metamorphosis fraction to a value comparable to that of no chemical stimulus present (Fig. 5D, E, Tables S21, S22). This suggests that pc2(+) cells which have access to the environment following settlement (possibly the ATENs) are sensing the chemical cues present in the sea water and promote or impede metamorphosis.

## Discussion

Settlement and metamorphosis are critical processes in the life cycle of a very wide range of marine organisms. The robust initiation of these processes in response to sensory cues has a profound impact on the population dynamics and evolutionary fitness of individual species as well as in the ecology and biodiversity of the oceans. Our study provides insights into the neuronal mechanisms of polymodal sensory perception initiating the initiation of settlement and metamorphosis in a marine planktonic larva equipped with one of the smallest nervous systems known to date^65^.

A key finding of our study is that the PSNs and ACCs show the ability to respond to a surprisingly diverse mechanical and chemical cues, demonstrating that these sensory cells are broadly tuned and polymodal. Utility of polymodal sensory cells that are capable of perceiving innocuous or noxious stimuli of various modalities is widespread across protostomes and deuterostomes^66,67^, where they have been extensively characterised in invertebrate models with simple nervous system such as the nematode *C. elegans* and the fruit fly *D. melanogaster* ^68–72^. Most vertebrate sensory systems are primarily specialized to detect one sensory modality, even though polymodal chemoreceptors and olfactory sensory receptor cells have been reported in vertebrates. These include the Grueneberg ganglion neurons and the carotid body chemoreceptors in mammals, as well as the chemoreceptor cells in the gills of fish^73,74^. Moreover, the septal organ olfactory sensory neurons (OSNs) in the mammalian olfactory neuroepithelium that can detect two discrete sensory modalities namely, chemical, and mechanical stimuli^75^ bearing strong resemblance to the functional properties of the PSNs and ACCs in this work. The presence of polymodal sensory cells in marine invertebrate chordates had previously not been documented. Furthermore, while numerous studies have hypothesized that sensory cells in the apical organ and other structures of zooplanktonic larvae can sense chemical and mechanical cues to induce larval settlement, the functional evidence has been lacking. This work suggests that polymodality is an ancient nervous system feature preserved in invertebrate chordates planktonic larvae, enabling the expansion of the sensory abilities of a miniaturized planktonic nervous system.

Furthermore, we provide functional demonstration that the ATENs are broadly tuned chemosensory cells. Previous work has postulated that the ATEN sensory neurons which originate from the proto-placodal ectoderm may have dual neurosecretory and chemosensory properties^62^. Our work presents functional evidence that the ATENs are ancestral chemosensory neurons capable of responding to different chemical stimuli. What may be the role of the ATENs in sensory transduction and information processing of chemosensory cues? A distinguishing feature of the ATENs relative to the PSNs and ACCs is their localization away from the adhesive organ. From an ethological point of view this distinct location may provide the larvae with the ability to track plumes of soluble chemical cues in three dimensions. Alternatively, they may serve as the main chemosensory-neurosecretory cells after settlement since exposure to the extracellular milieu of the PSNs and ACCs is found to be blocked following attachment. Intriguingly, *Ci-gnrh1* is expressed in the adhesive organ (likely the PSNs)^76^ while *Ci-gnrh2* is expressed in the aATENs^77^. GnRH peptides encoded by these genes have been shown to promote initiation of metamorphosis^78^. Thus, it is reasonable to hypothesize that a GnRH based ‘wireless’ network may also play an important role in the integration of sensory information related to settlement and metamorphosis. Furthermore, it is conceivable that the availability of the ATENs to respond to sensory cues following settlement may be important for the stimulus driven release of GnRH peptides to regulate the rate of metamorphosis. Indeed, the chemogenetic behavioral evidence provided here supports our model that chemical cues are able to control the rate of metamorphosis via pc2(+) cells that include, but are not limited to the PSNs and the ATENs. Given that the PSNs lose their access to the extracellular milieu following settlement the ATENs are the likely chemosensors employed by larvae in the post-settlement period. More generally, our functional imaging and chemogenetic analyses suggest that chemosensation is a major sensory modality employed by *C. intestinalis* larvae to modulate settlement and metamorphosis.

That the finding that the PSNs and their neighbouring cells the ACCs present in the papillae respond to the same mechanical and chemical cues (at least for those tested in both cell types) suggests that the adhesive organ of *C. intestinalis* larvae may operate under a coincidence detection mechanism, whereby activation of both the PSNs and ACCs across the papillae is required to elicit responses from downstream interneurons and a robust behavioral output. Interestingly, coincidence detection-like mechanisms have been sparsely identified in invertebrates^79,80^, but have been extensively documented in various auditory, visual and somatosensory circuits in vertebrates ^81–83^. The observation that the ATENs are responding to some of the same chemosensory cues as the PSNs raises the possibility that they can function in parallel to sense external chemical stimuli and transmit sensory input to the central nervous system. This phenomenon has been documented in multiple contexts in C. elegans^84–86^. The importance of the PSNs and ATENs for action selection in the brain in response to positive or negative valence settlement sensory inputs is corroborated by our whole brain imaging data. In the absence of sensory input, the animals explore a small part of the neuronal dynamics space. This situation changes drastically when the brain receives input from the PSNs and ATENs in response to stimulus presentation and/or removal, which leads to an extensive exploration of neuronal dynamics space and short or long-lasting alterations to neuronal state depending on the stimulus valence.

Why would it be beneficial for *C. intestinalis* to have a coincidence detection, or a combinatorial action mechanism associated with the settlement mediating neural circuit? We have shown that the papillae are the major transduction site for cues that promote or deter larval settlement and metamorphosis. The presence of multiple independent inputs which converge to a shared information processing centre in the brain would reduce noise and increase the robustness of the sensory responses in this circuit. Aberrant signalling in the circuit could lead to the rejection of a suitable site, or even worse, result in settlement in an unsuitable site. Given the critical importance of the settlement and metamorphosis processes, ensuring that the larvae identify a suitable substrate and reject others based on the sensory input they receive is fundamental to the evolutionary fitness of animals. It is worth noting that we did not detect single-neuron representations of sensory stimuli downstream of the sensory neurons but rather a sparse representation of the sensory cues across the brain which is in line with what has been observed in terrestrial invertebrate and vertebrate model systems^87–89^. To what extent this strategy is conserved across other zooplanktonic organisms is not clear. Functional imaging studies on chemosensory systems of marine zooplanktonic organisms are very sparse, with the notable exception of a study using the marine annelid Platynereis dumerilii^90^, where it was shown that in juveniles four types of sensory organs respond with different efficacy to chemical cues. From an evolutionary perspective, the mechanosensory and chemosensory-neurosecretory functions of the PSNs, ACCs and ATENs may have been segregated into dedicated cell types that function together in coherent multi-layered circuits. Cellular subfunctionalization likely has been a generally important mechanism of neuronal circuit evolution in vertebrates.

Our work demonstrated that both the PSNs and the ACCs can respond to two different types of mechanical stimuli (press and poke) with distinct response dynamics. The press stimulus elicited a transient response to stimulation-onset and stimulation-offset in both cell types. This response profile is observed in vertebrate Pacinian corpsuscles^91,92^ and more recently in trigeminal mechanoreceptors innervating the mouse tongue^93^. In contrast, poke stimuli yielded intermediately adapting responses reminiscent of those observed in some vertebrate nociceptors^94–96^. Our findings suggest that the pre-vertebrate chordate sensory cells possess multiple mechanosensory modalities, which were segregated in the vertebrate lineage. In addition, multiple mechanically gated ion channels must function in the palps. A recent study suggested that mechanical activation of the PSNs is partially mediated by the TRP channel PKD2^31^, however it is very likely that more mechanosensitive ion channels function in these cells and the ACCs because only a subtle phenotype was reported.

Our behavioral data revealed that chemical sensory cues which elicit Ca^2+^ responses in the PSNs and the ACCs influence the ethologically relevant behaviors of settlement and metamorphosis. Here, a concentration dependent increase in settlement and metamorphosis rates in response to NH^4+^, Cs+ and K^+^ ions, with NH^4+^ represents the strongest settlement cue confirms and extends previously findings in ascidians and other marine larvae^38,39,47^. NH^4+^ may be an important settlement cue due to interactions between ammonia producing bacteria and algae in the ocean and the importance of algae as a food source for juvenile and adult *C. intestinalis*. While the identity of the NH^4+^ sensor is unknown, plausible candidates include the ammonium transporters Ci-AMT1a and b which appear to be expressed in the palps and in the ATENs^97^. Notably, Amt in *Drosophila* has been shown to be a non-canonical olfactory receptor for ammonia^98^. Other candidate receptors may include members of the recently identified MS4A chemoreceptor family^99^ which have been recently shown to be expressed in the palps^100^. Future studies will leverage the freshly identified PSN and established ACC specific promoters^61,101,102^ to determine the role of these receptors in chemosensation using cell type specific genome editing.

Our results show that some sugars, short alcohols, and amino acids have a positive settlement and metamorphosis valence. From an ethological perspective, the positive valance of these compounds could be attributed to the suitability of the source from which they originate. For example, sugars are the degradation products of the polysaccharides present on biofilms, seagrass or algae coated solid structures which can act as favourable settlement and metamorphosis sites for a large range of marine larvae^48^.

Negative cues to settlement have been harder to identify compared to positive ones (reviewed in). A novel finding of our study is the identification of terpenoids as biogenic volatile organic compounds with a negative settlement and metamorphosis valance for *C. intestinalis* larvae. Terpenoids are amongst the most abundant class of natural metabolite products in terrestrial and marine environments with a broad range of signalling functions including defence in marine and terrestrial organisms^57,103,104^. The function of terpenoids as negative valence settlement and metamorphosis cues is unlikely to be restricted to *C. intestinalis*, as previous studies have shown that terpenes prevent settlement of barnacle larvae ^105,106^. Octopuses are also capable of sensing terpenes by employ specialized chemotactile sensory receptors on their arms^107–109^. Whether these or similar chemotactile receptors are present in *C. intestinalis* is unknown, however, this study raises the possibility that similarly to octopus, *C. intestinalis* larvae employ a ‘taste by touch’ system to sense poorly soluble chemical cues. Therefore, it is conceivable that like other organisms that make use of the benthos^103,104,110^, *C. intestinalis* larvae is capable of following chemical traces adherent to the substrate using their palps, allowing them to evaluate potential settlement and metamorphosis sites.

We also found that contaminants of human-made chemicals or activities such as the metal ion Cu2+ and the surfactant SDS^49,50^ strongly inhibited settlement and metamorphosis, leading to eventual larval death, indicating that anthropogenic pollutants are perceived by the sensory machinery of the larva and interfere with settlement and metamorphosis.

A recurring question in the field is whether the same sensory cues can induce both settlement and metamorphosis, or different cues are involved in the two processes^45^. The answer to this question remains unexplored for most marine invertebrate species^45^. Our study shows that in *C. intestinalis* larvae, most inducing or impeding cues affect the entire recruitment process, which includes both settlement and metamorphosis.

In summary, we show that *C. intestinalis* larvae have the capability to differentially respond to multiple mechanical and chemosensory cues with positive or negative valance for settlement and metamorphosis which opens the possibility of complex multimodal sensory integration, a surprising feat for an animal with such a streamlined nervous system. In the future, understanding how the sensory processes underlying settlement and metamorphosis are affected by the numerous anthropogenic influences that affect the ocean such as acidification and underwater noise will emerge as a key challenge at the interface of ecology and neurobiology. *C. intestinalis* may hold the key to address this challenge.

## Acknowledgments

We would like to thank Dr. Mie Wong for her feedback on the manuscript.

## Data and Code Availability

For calcium imaging analysis done outside Mesmerize the code is available from here: https://github.com/ChatzigeorgiouGroup/Hoyer_etal_2023

## Author contributions

K.K. and M.C. conceived the project. K.K., M.C. and J.H. designed the experiments, J.H., K.K., Z.L., M.C. and M.v.B carried out the experiments. J.H., analyzed the data with help from D.D., K.K, A.A.. M.C. and J.H. wrote the paper with input from all authors.

## Methods

### Animal collection

We collected adult *C. intestinalis* (Type B) from: Døsjevika, Bildøy Marina AS, postcode 5353, in Bergen, Norway. The collection site can be identified using the following GPS coordinates: 60.344330, 5.110812.

### Rearing conditions for adult *C. intestinalis*

Adult C. intestinalis (Type B) were housed in our purpose-made facility at the Michael Sars Centre. Around 50 to 100 adults were kept in large 50L tanks with running sea water. The temperature of the sea water was maintained at 10°C with constant light and food supply (this included several species of diatoms and brown algae) to increase egg production and reduce the likelihood of spontaneous spawning^111^.

### Electroporations of zygotes and staging of embryos

We performed egg collection, fertilization and rearing following standard protocols^112^. We dissected gravid adult *C. intestinalis* to obtain mature eggs and sperm for *in vitro* fertilization. We dechorionated eggs using chemical dechorionation in a 1% sodium thioglycolate and 0.1% pronase mix dissolved in filtered seawater. We placed the eggs solution on a rocker for approximately 10 min until we observed that zygotes were completely dechorionated. We washed dechorionated eggs several times using artificial sea water and then we fertilized with sperm for ∼10 min. We performed electroporations according to published protocols with minor modifications^113^. After we thoroughly washed the fertilized eggs, we electroporated them in a mannitol solution with 70 to 160 μg of DNA depending on the anticipated levels of expression for each construct. We electroporated fertilized eggs using 4 mm gap electroporation cuvettes (MBP Catalog #5540). To deliver the pulses we made use of a BIORAD GenePulserXcell equipped with a CE-module. We used the following settings: Exponential Protocol: 50 V, Capacitance: 800–1300μF, Resistance : οο. Typically we obtained electroporation time constants between 12–30 milliseconds. We cultured embryos in ASW (artificial sea water, Red Sea Salt) at 14°C until we used them for experiments. We set the pH of the ASW to 8.4 at 14°C. The salinity of the ASW was set to 3.3–3.4%.

### Molecular cloning

To obtain *βγ-crystallin* and *pc2* promoter we used gDNA at a concentration of 100-150ng/μl, the primers shown in table S23, dNTPs (Thermofisher, R0182 mix) and Q5 High-Fidelity DNA Polymerase (M0491L, NEB) to perform PCR reactions. We gel purified the PCR products using Zymogclean Gel DNA Recovery Kit (Zymo research, D4002). These were inserted into the P4-P1R vector using BP Clonase II (Invitrogen, P/N56480). We identified positive clones using restriction digest and we sequenced multiple clones with Sanger sequencing. For generating the hM4D::mCherry construct we used the same approach with the difference that the purified PCR product was inserted in a pDONR221 vector. The PCR template for the hM4D::mCherry was the following plasmid: pAAV-hSyn-hM4D(Gi)-mCherry which was a gift from Bryan Roth (Addgene plasmid # 50475; http://n2t.net/addgene:50475; RRID:Addgene_50475). The relevant PCR primers are shown in table S23. Generation of middle position GCaMP6s has been previously described^19^.

We performed four-way Gateway reactions choosing one of the promoters each time in the 1^st^ position, with an appropriate 2^nd^ position entry clone and the unc-54 3’ UTR in the 3^rd^ position. We recombined these entry clones into a pDEST II backbone using LR Clonase II (Invitrogen, P/N56485).

We generated high-quality DNA of the expression constructs for electroporations using the NucleoBond Xtra Midi kit (Macherey-Nagel 740410.50)

### Larval settlement assay

Chorionated embryos grown at 14°C hatched as larvae on average at 36 hours post fertilization (h.p.f.)^111^ and they were collected from multiple 9cm plates coated with agarose to be were transferred to fresh 9cm plates also coated with agarose in ASW. When the larvae reached 43h.p.f they were transferred to 3cm plastic petri-dishes (not coated). We transferred 100 swimming larvae per 3cm plate. Most of the ASW was removed and 3ml of the15hemicall cue diluted in ASW was used to fill the 3cm plate. To determine whether larvae had settled we used a 14cm glass Pasteur pipette with a rubber bulb, which was used to swirl gently the water. The movement of larvae during the swirling was monitored under a dissecting microscope to determine whether larvae settled or not to the bottom of the petri dish (and rarely on the sides of the dish). The larvae that had settled could not move from the site at which they had permanently adhered. The number of larvae that had stably attached to the petri-dish were counted every 30 minutes for a period of 10 hours (0 to 600 minutes). During the experiments the plates were kept in tabletop incubators at 14°C without any light. Data was plotted using Graphpad Prism 9. For statistical analysis we performed Two-way RM ANOVA, followed by either Dunnett’s or Tukey’s multiple comparisons test for multiple comparisons. In addition, we applied standard curves (sigmoidal, 4PL) to determine the time point at which 50% of larvae had settled for a given stimulus.

### Tail regression (metamorphosis) assay

Approximately 500 chorionated larvae ca 43h.p.f. at 14°C were transferred in 3cm plastic petri dishes with ASW. We monitored settlement every 30 minutes until ca 100-150 larvae had settled. We then removed most of the ASW with the remaining larvae that had not settled. We replaced the ASW with either 3ml of ASW or the chemical cues diluted in ASW. The fraction of larvae metamorphosed (fully resorbed tails) was monitored under a dissecting microscope. When approximately 50% of the control (just ASW) larvae had metamorphosed we quantified the fraction of metamorphosed larvae across the different chemical treatments. We measured 30 plates for each chemical cue treatment, (5 plates per treatment, over 6 different days). During the experiments the plates were kept in tabletop incubators at 14°C without any light. Data was plotted using Graphpad Prism 9. For statistical analysis we first checked for normality of our data using the Shapiro-Wilk test. We then performed a Kruskal-Wallis test with a correction for multiple comparisons using Dunn’s test.

### Mechanical stimulation assays

Electroporated embryos were raised in artificial seawater (ASW) at 14°C until hatching. The hatched animals were incubated in 99.97% ASW, with pH 8.4 at 14°C and 32% salinity, and 0.03% MS-222 for immobilization, in a 9 cm petri dish under the microscope objective. By applying gentle suction from a customized glass pipette, with an outer diameter of 120 um and inner diameter of 25 um, fitting roughly half the size of the head, the animals were held in position for stimulation. We illuminated animals using a mercury lamp with a BP470/20, FT493, BP505-530 filter set. The animals were imaged under a 40x water immersion objective mounted on an Axioskop A1 equipped with a Hamamatsu Orca Flash 4.0V2 CMOS camera. The field of view was 320μm in each direction (2048×2048 pixels, with the FOV cropped to a box around the palp cells).

### Poke assay

The mechanical poke assay started by recording a pre-stimulus period of 10 seconds. Upon the completion of these 10 seconds a blunt glass needle connected to a micromanipulator (Sutter Instrument MPC-365 System) was moved towards the animal so that it could touch the pad of the palps briefly before it was withdrawn to its starting position. We then kept recording the calcium activity in the palp cells for a minimum post stimulation period of 1 minute until the calcium signal had decreased to the intensity level it had before excitation. Data was acquired using the Hamamatsu Corporation imaging software HCImageLive at 40 Hz.

The intensity levels of the imaged cells were then extracted using Mesmerize. First adjusting for any movement using NoRMCorre^114^ motion correctionthrough Mesmerize^19^ followed by signal extraction by manually drawing regions of interest (ROI) around each cell producing a calcium trace for each ROI. Since the promoters for the PSN and ACC cells only expressed GCaMP in a handful of spatially distinct cells within the palps, it was not necessary to demix or subtract fluctuating neuropil background during signal extraction. The traces obtained were imported to a Mesmerize project for annotation and organization in a pandas DataFrame^115^.

### Press assay

After 10 seconds a blunt glass needle connected to a micromanipulator (Sutter Instrument MPC-365 System) were introduced to press the palp 5 to 10 um inwards. The glass needle was kept in this position for 20 seconds before it was quickly withdrawn to its starting position. We then kept recording the calcium activity in the palp cells for a minimum post stimulation period of 1 minute until the calcium signal had decreased to the intensity level it had before excitation. Data was acquired using the Hamamatsu Corporation imaging software HCImageLive at 40Hz.

The intensity levels of the imaged cells were then extracted using Mesmerize. The signal was extracted by drawing regions of interest (ROI) around each cell producing a calcium trace for each ROI. The traces obtained were imported to a Mesmerize project for annotation and organization in a data frame.

### Chemical stimulation assays

#### Palps chemical stimulation assay

Electroporated embryos were raised in ASW at 14°C until hatching. The hatched animals were incubated in 99.97% ASW, with pH 8.4 at 14°C and 32% salinity, and 0.03% MS-222 for immobilization, in a 9 cm petri dish under the microscope objective. The perfusion pencil which was used for ASW and stimulus perfusion to the animal was placed at a distance of about 250 μm, pointing directly towards the palps of the animals. We illuminated animals using a mercury lamp with a BP470/20, FT493, BP505-530 filter set. The animals were imaged under a 40x water immersion objective mounted on an Axioskop A1 equipped with a Hamamatsu Orca Flash 4.0V2 CMOS camera. The field of view was 320μm in each direction (2048×2048pixels). The first 8 seconds of the experiments the animals were perfused with a continuous flow of a control solution consisting of 99.97% ASW and 0.03% MS222 before switching to the stimulation solution for 4 seconds. We then switched back to perfusing with the control solution and kept recording the calcium activity in the palp cells for a minimum post stimulation period of 1 minute until the calcium signal had decreased to the intensity level it had before excitation. Data was acquired using the Hamamatsu Corporation imaging software HCImageLive at 40Hz.

The intensity levels of the Imaged cells were then extracted using Mesmerize. First adjusting for any movement using the NORMCorre^114^ motion correction followed by signal extraction by drawing regions of interest (ROI) around each cell producing a calcium trace for each ROI. The traces obtained were imported to a Mesmerize project for annotation and analysis.

### Whole brain chemical stimulation assay

Electroporated embryos were raised in ASW at 14°C until hatching. The hatched animals were incubated in a 6 cm petri dish with glass bottom in 98.5% ASW, with pH 8.4 at 14°C and 32% salinity, and 1.5 % agarose (in ASW) for immobilization. The solidified agarose around the anterior part of the trunk were carefully removed using a tweezer to expose the palps and anterior part of the trunk before 2ml of ASW was added to cover the agarose. A perfusion pencil was positioned at the edge of the lens of the objective pointing directly towards the palps of the animals. We originally attempted to image the larvae using a two-photon excitation microscope, but we found that illumination of the brain regions close to the pigment cells, led to significant light-induced-damage. Thus, the animals were imaged under a 40x objective mounted on an Olympus iXplore SpinSR10 confocal microscope equipped with two Hamamatsu Orca Flash 4.0V2 CMOS cameras. The FOV (1024×1024 with 2×2 pixel binning) was cropped to fit only the trunk of the animal. For the live imaging of the calcium intensity levels in the nervous system, we illuminated the animals using a Coherent 488nm laser mounted on the confocal microscope for approximately 288 volumetric cycles, adjusting the number respectively based on the size of the cropped FOV to set the time of the experiment to 3 minutes. The laser illuminated the animals with a step of 0.32um on the z-axis within a range of 28.8um. The data was acquired at approximately 1.6 brain volumes per second. The animals were firstly perfused with a continuous flow of control solution (ASW) for 30 seconds before switching to a stimulation solution for 30 seconds. We then switched back to perfusing with the control solution and kept recording for 2 minutes. For the accompanied two-channel stacks where we imaged both the nucleus and the cytoplasm, we illuminated the animals using a Coherent 488nm laser and a Coherent 561nm laser mounted on the confocal microscope. The lasers illuminated the animals and we collected 180s at 7 slices on the z-axis with a step of 0.38um within a range of 28.8um. All data was acquired using the Olympus software cellSens Dimensions.

The GCaMP6s intensity levels of the imaged cells were then extracted using Mesmerize for signal extraction of 3-dimensional imaging, CNMF-3D^116,117^.. The traces obtained were imported to a Mesmerize project for annotation and analysis. The traces were then z-scored using tslearn^118^. Principal component analysis (PCA) was then performed on z-scored traces to obtain a low dimensional (3 component) trajectory for each of the recorded samples using the Python library scikit-learn^119^.

