## Supplementary Data for "Polymodal sensory perception of mechanical and chemical cues drives robust settlement and metamorphosis of a marine pre-vertebrate zooplanktonic larva"

A

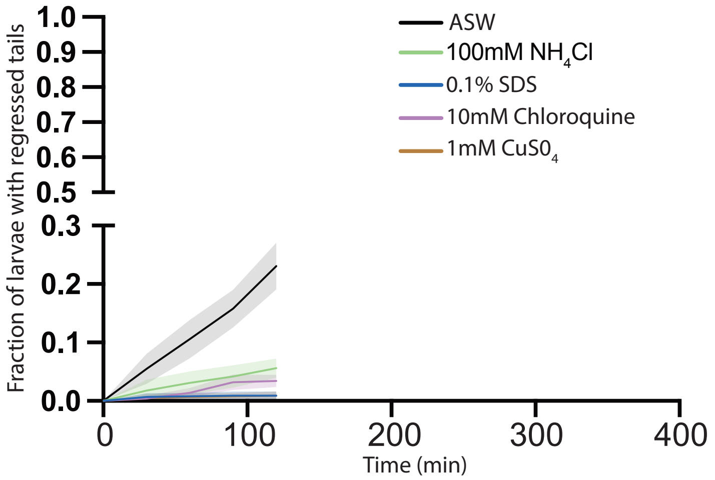

ASW
100mM $NH_4Cl$
0.1% SDS
10mM Chloroquine
1mM $CuS0_4$
Fraction of larvae with regressed tails
1.0
0.9
0.8
0.7
0.6
0.5
0.3
0.2
0.1
0.0
0
100
200
300
400
Time (min)

**fig. S1 Noxious cues prevent settlement.**

(A) Settlement curves for larvae exposed to ASW, 100mM $NH_4Cl$, 0.1% SDS, 10mM Chloroquine and 1mM $CuSO_4$. For data shown in panels A and B we performed a two-way RM ANOVA, followed by Dunnett's multiple comparisons test (table S11).

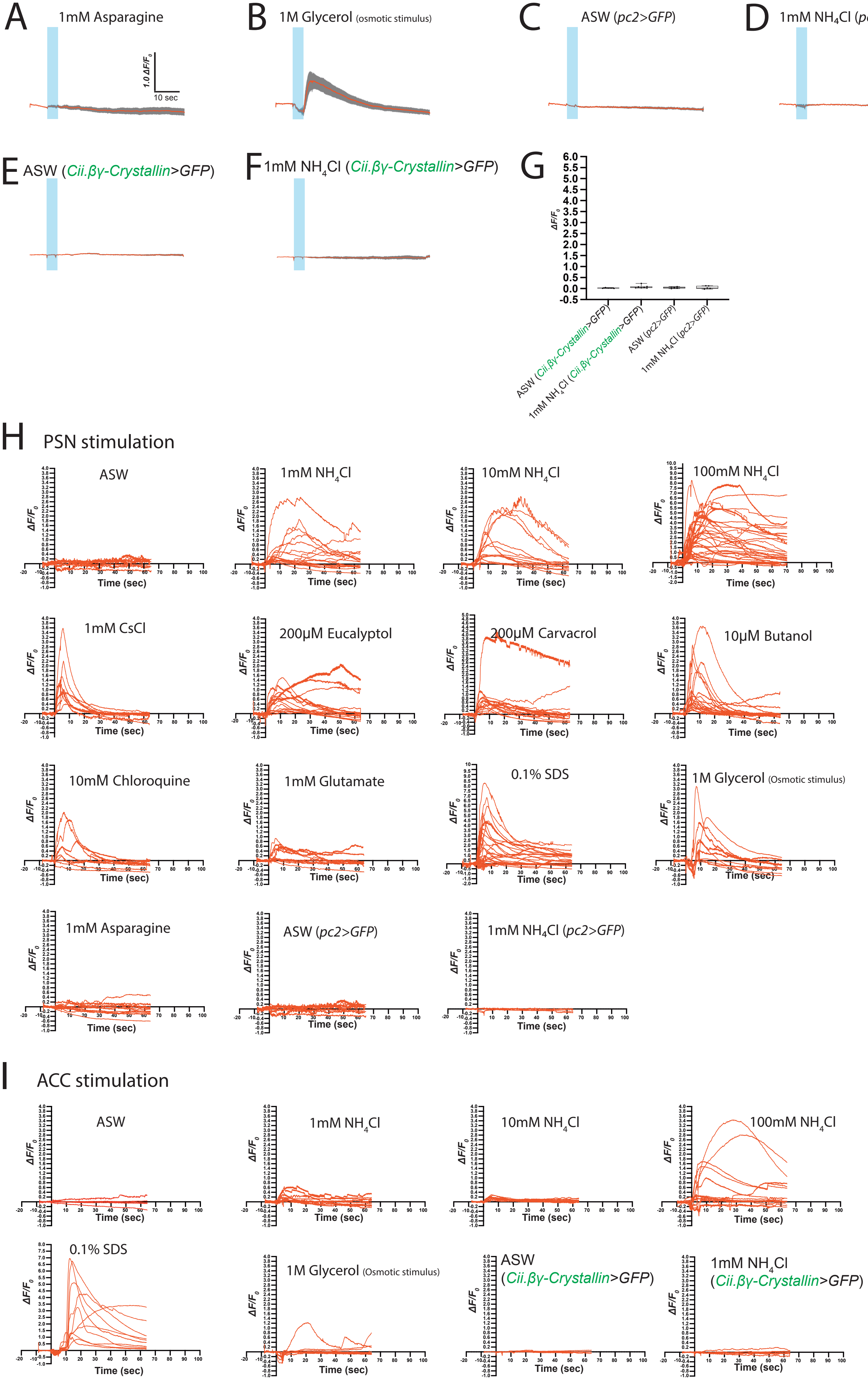
A
1mM Asparagine
1.0 ΔF/F0
10 sec
B
1M Glycerol (osmotic stimulus)
C
ASW (pc2>GFP)
D
1mM NH4Cl (pc2>GFP)
E
ASW (Cii.βγ-Crystallin>GFP)
F
1mM NH4Cl (Cii.βγ-Crystallin>GFP)
G
ΔF/F0
ASW (Cii.βγ-Crystallin>GFP)
1mM NH4Cl (Cii.βγ-Crystallin>GFP)
ASW (pc2>GFP)
1mM NH4Cl (pc2>GFP)
H
PSN stimulation
ASW
1mM NH4Cl
10mM NH4Cl
100mM NH4Cl
1mM CsCl
200μM Eucalyptol
200μM Carvacrol
10μM Butanol
10mM Chloroquine
1mM Glutamate
0.1% SDS
1M Glycerol (Osmotic stimulus)
1mM Asparagine
ASW (pc2>GFP)
1mM NH4Cl (pc2>GFP)
ΔF/F0
Time (sec)
I
ACC stimulation
ASW
1mM NH4Cl
10mM NH4Cl
100mM NH4Cl
0.1% SDS
1M Glycerol (Osmotic stimulus)
ASW (Cii.βγ-Crystallin>GFP)
1mM NH4Cl (Cii.βγ-Crystallin>GFP)
ΔF/F0
Time (sec)

**fig. S2 Average traces and individual responses to chemical stimuli and controls in PSNs and ACCs.**

(A) PSN average trace of $Ca^{2+}$ responses to 1mM Asparagine. (B) PSN average trace of $Ca^{2+}$ responses to 1M Glycerol. (C-F) Control experiments showing that average traces of *pc2>GFP* or *$\beta\gamma$-crystallin>GFP* intensity calculated as $\Delta F/F_0$ in response to a switch between two channels (B, D) both containing ASW, (C, E) ASW and 1mM $NH_4Cl$. (G) Summary of data in (B-E) showing peak amplitude $\Delta F/F_0$. (H, I) Individual $Ca^{2+}$ transients recorded in the PSNs (H) and ACCs (I) grouped according to the chemical stimulus delivered. The number of animals used in panels A and B are indicated in table S11, and for panels B-F in table S14.

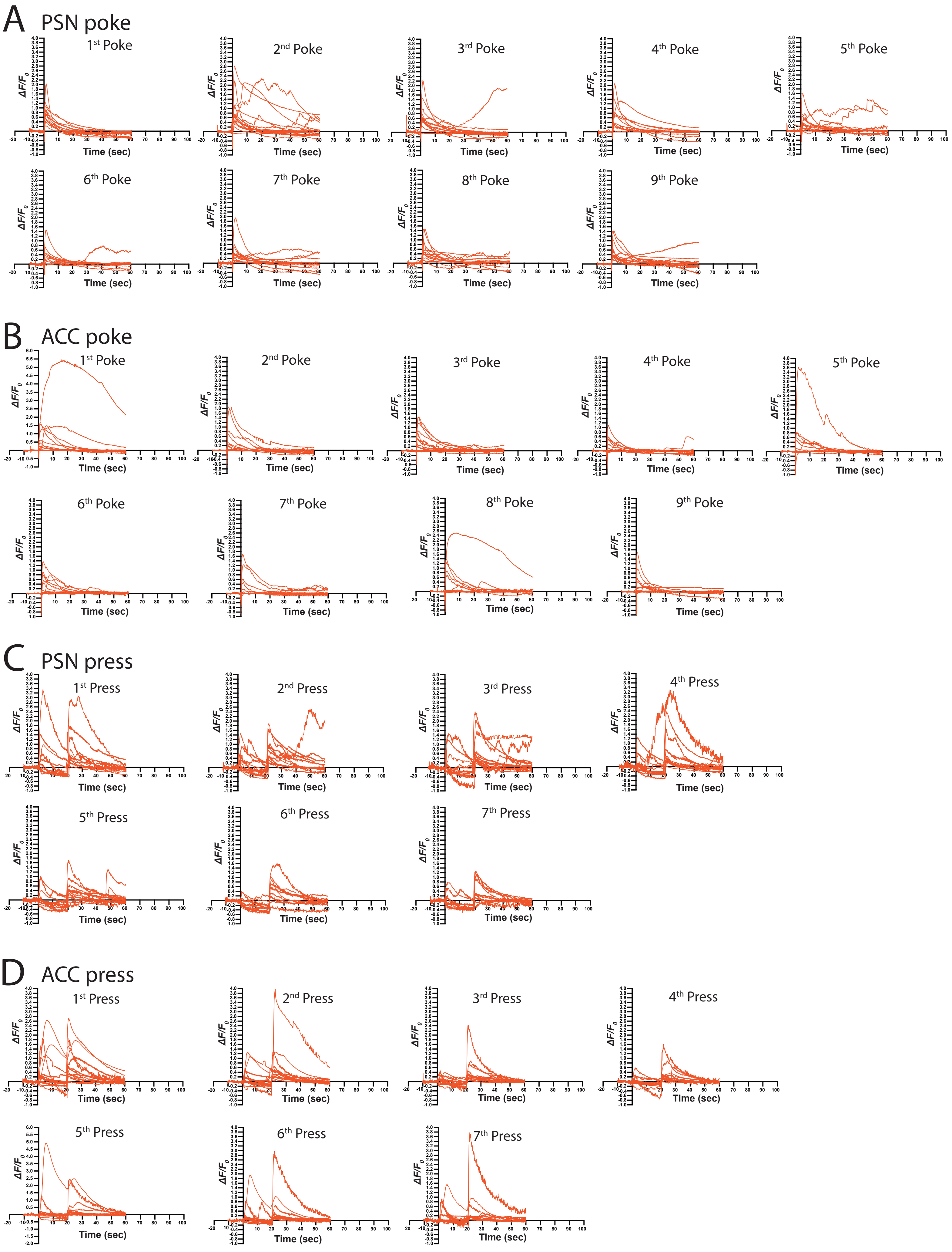
A
PSN poke
1st Poke
2nd Poke
3rd Poke
4th Poke
5th Poke
6th Poke
7th Poke
8th Poke
9th Poke
ΔF/F0
Time (sec)
B
ACC poke
1st Poke
2nd Poke
3rd Poke
4th Poke
5th Poke
6th Poke
7th Poke
8th Poke
9th Poke
C
PSN press
1st Press
2nd Press
3rd Press
4th Press
5th Press
6th Press
7th Press
D
ACC press
1st Press
2nd Press
3rd Press
4th Press
5th Press
6th Press
7th Press

**fig. S3 Individual $Ca^{2+}$ responses to repeated mechanical press and poke stimuli in PSNs and ACCs.**

Individual PSN (A) and ACC (B) $Ca^{2+}$ responses (red traces) to mechanical poke stimulation grouped according to trial number ($1^{st}$ to $9^{th}$ stimulus repetition). Individual PSN (C) and ACC (D) $Ca^{2+}$ responses (red traces) to mechanical press stimulation grouped according to trial number ($1^{st}$ to $7^{th}$ stimulus repetition).

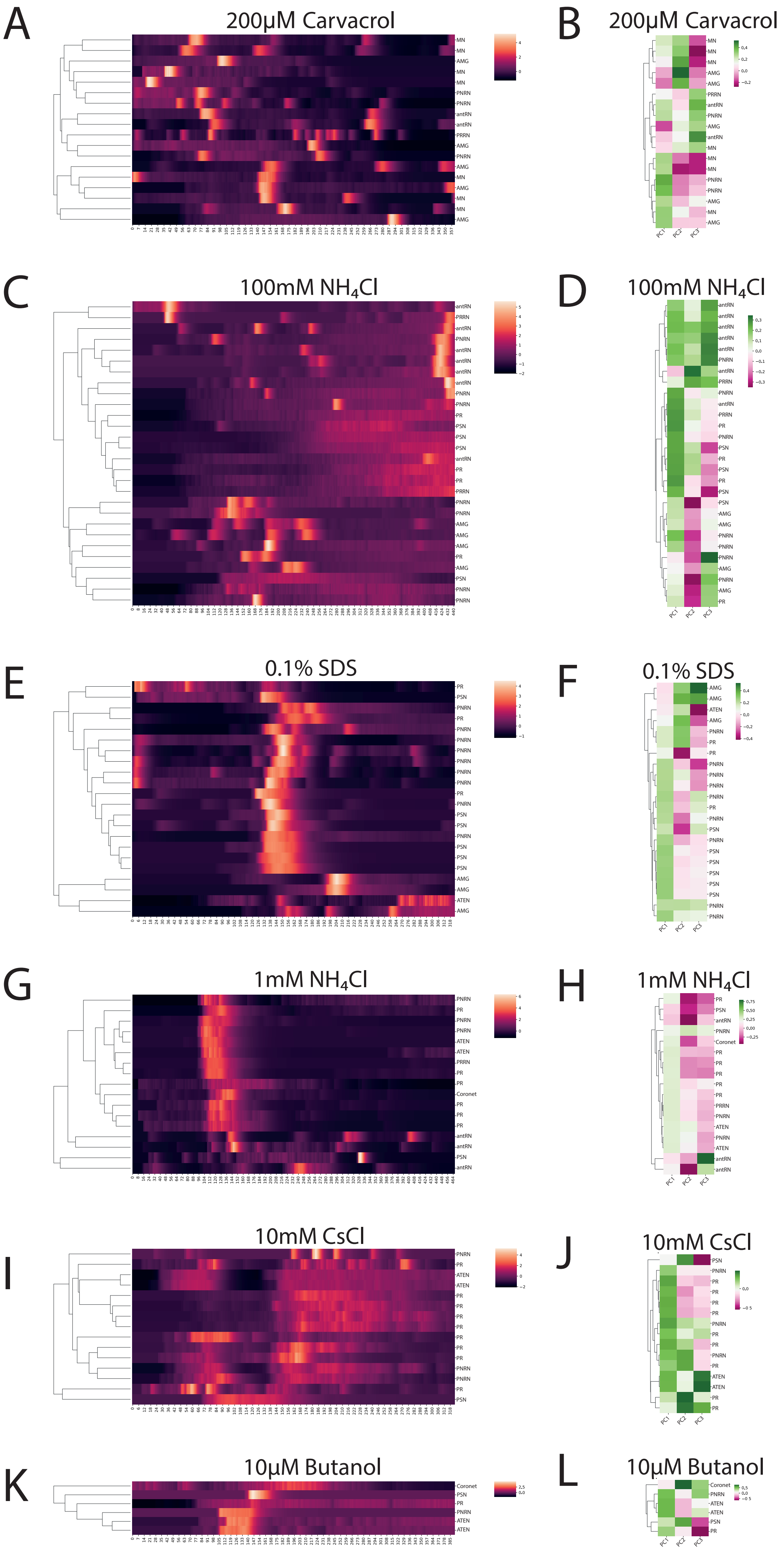
A
200μM Carvacrol
B
200μM Carvacrol
C
100mM NH4Cl
D
100mM NH4Cl
E
0.1% SDS
F
0.1% SDS
G
1mM NH4Cl
H
1mM NH4Cl
I
10mM CsCl
J
10mM CsCl
K
10μM Butanol
L
10μM Butanol

**fig. S4 Hierarchically clustered heatmaps of z-scored whole brain imaging data and PCA loading corresponding to Fig. 3.**

(A,C,E,G,I,K) Hierarchically clustered heatmaps of z-scored $Ca^{2+}$ activity and (B,D,F,H,J,L) PCA loading maps for (A, B)200μM Carvacrol, (C, D) 100mM $NH_4Cl$, (E, F) 0.1% SDS, (G, H) 1mM $NH_4Cl$, (I, J) 10mM CsCl and (K, L) 10μM Butanol.

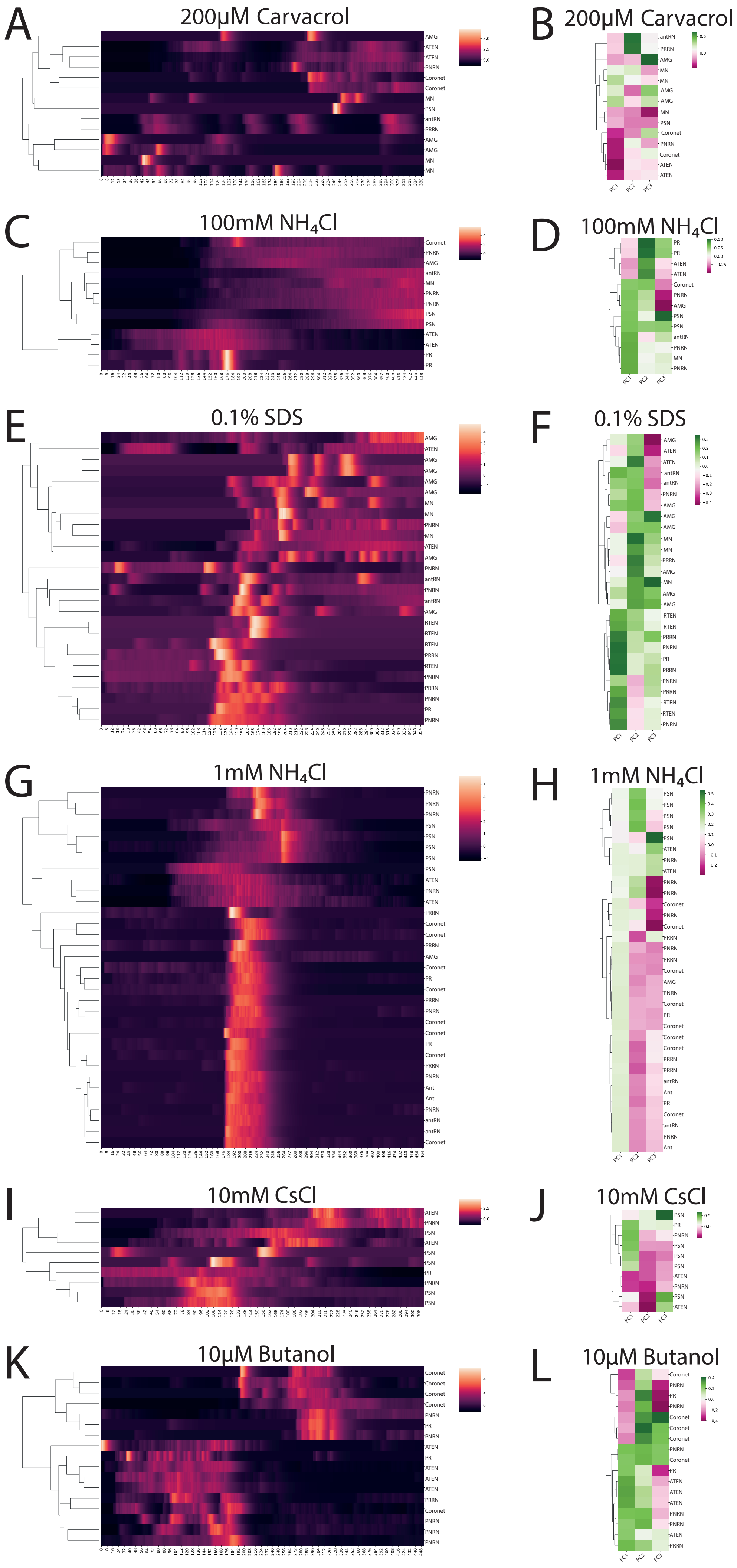
A
200µM Carvacrol
B
200µM Carvacrol
C
100mM NH4Cl
D
100mM NH4Cl
E
0.1% SDS
F
0.1% SDS
G
1mM NH4Cl
H
1mM NH4Cl
I
10mM CsCl
J
10mM CsCl
K
10µM Butanol
L
10µM Butanol

**fig. S5 Hierarchically clustered heatmaps of z-scored whole brain imaging data and PCA loading corresponding to Fig. 4.**

(A,C,E,G,I,K) Hierarchically clustered heatmaps of z-scored $Ca^{2+}$ activity and (B,D,F,H,J,L) PCA loadings for (A, B)200μM Carvacrol, (C, D) 100mM $NH_4Cl$, (E, F) 0.1% SDS, (G, H) 1mM $NH_4Cl$, (I, J) 10mM CsCl and (K, L) 10μM Butanol.

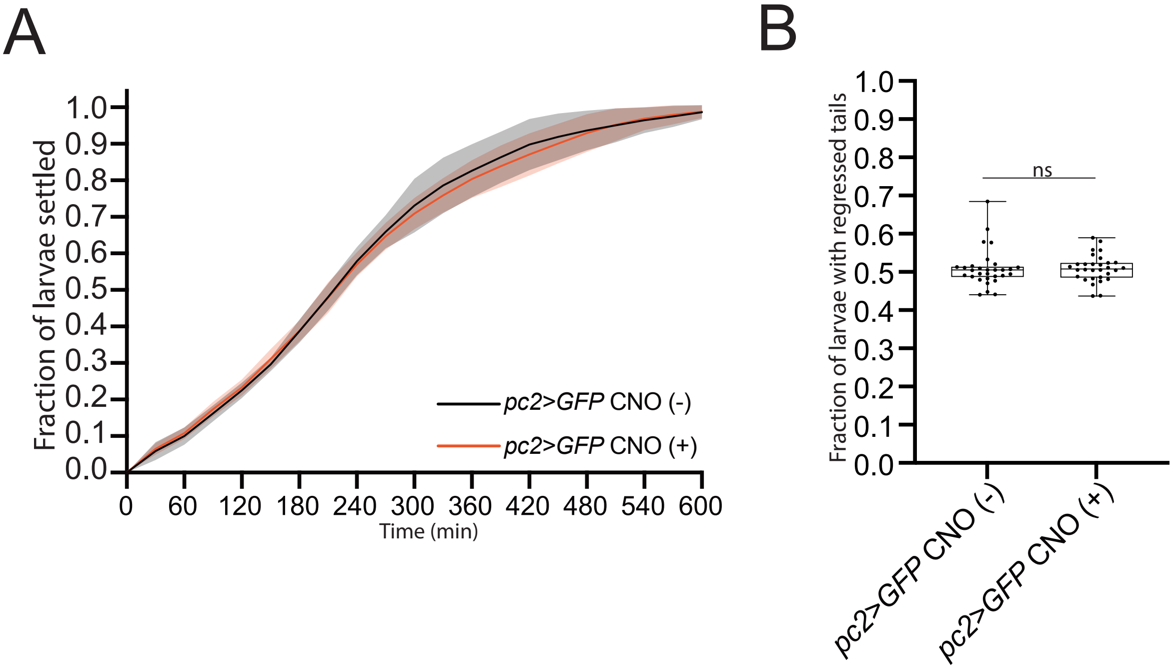

A
B
Fraction of larvae settled
1.0
0.9
0.8
0.7
0.6
0.5
0.4
0.3
0.2
0.1
0.0
0
60
120
180
240
300
360
420
480
540
600
Time (min)
pc2>GFP CNO (-)
pc2>GFP CNO (+)
Fraction of larvae with regressed tails
ns
pc2>GFP CNO (-)
pc2>GFP CNO (+)

**fig. S6 Clozapine-N-Oxide is not a settlement and metamorphosis cue for *C. intestinalis*.**

(A) Larval settlement curves for *pc2>GFP* transgenic larvae in the presence or absence of CNO. For statistical analysis we used a Two-way RM ANOVA, followed by Dunnett's multiple comparisons test (table S15) (B) Fraction of larvae metamorphosed for *pc2>GFP* transgenic larvae in the presence or absence of CNO. For statistical analysis we performed a Welch's test (table S16).

**S1 Movie**
A PSN neuron expressing GCaMP6s responds to mechanical stimulation (poke).

**S2 Movie**
A PSN neuron expressing GCaMP6s responds to 10mM Chloroquine.

**S3 Movie**
Volumetric whole-brain $Ca^{2+}$ imaging using *Cii.Etr- 1>nls::GCaMP6s::nls*. The larva was stimulated with 10mM CsCl.

**S4 Movie**
Volumetric whole-brain $Ca^{2+}$ imaging using *Cii.Etr- 1>nls::GCaMP6s::nls*. The larva was stimulated with 10µM Butanol.

**S5 Movie**
Volumetric whole-brain $Ca^{2+}$ imaging using *Cii.Etr- 1>nls::GCaMP6s::nls*. The larva was stimulated with 200µM Carvacrol.

**S6 Movie**
Volumetric whole-brain $Ca^{2+}$ imaging using *Cii.Etr- 1>nls::GCaMP6s::nls*. The larva was stimulated with 0.1% SDS.

table S1 Statistics for settlement rates analysis in the presence of $NH_4Cl$

| Figure & Panel | Test | Data point | Comparison | P Value summary | P Value |
|---|---|---|---|---|---|
| Fig. 1C | Two-way RM ANOVA | | | | |
| | | | Time | **** | <0,0001 |
| | | | Fraction Settled | **** | <0,0001 |
| | | | Time x Fraction Settled | **** | <0,0001 |
| | Dunnett's multiple comparisons test | | | | |
| | | 30 min | ASW vs. 1mM $NH_4Cl$ | ns | 0,2657 |
| | | | ASW vs. 10mM $NH_4Cl$ | *** | 0,0003 |
| | | | ASW vs. 0.5mM $NH_4Cl$ | ns | 0,9466 |
| | | 60 min | ASW vs. 1mM $NH_4Cl$ | **** | <0,0001 |
| | | | ASW vs. 10mM $NH_4Cl$ | **** | <0,0001 |
| | | | ASW vs. 0.5mM $NH_4Cl$ | ns | 0,7286 |
| | | 90 min | ASW vs. 1mM $NH_4Cl$ | **** | <0,0001 |
| | | | ASW vs. 10mM $NH_4Cl$ | **** | <0,0001 |
| | | | ASW vs. 0.5mM $NH_4Cl$ | ns | 0,9543 |
| | | 120 min | ASW vs. 1mM $NH_4Cl$ | **** | <0,0001 |
| | | | ASW vs. 10mM $NH_4Cl$ | **** | <0,0001 |
| | | | ASW vs. 0.5mM $NH_4Cl$ | ns | 0,6667 |
| | | 150 min | ASW vs. 1mM $NH_4Cl$ | **** | <0,0001 |
| | | | ASW vs. 10mM $NH_4Cl$ | **** | <0,0001 |
| | | | ASW vs. 0.5mM $NH_4Cl$ | ns | 0,5017 |
| | | 180 min | ASW vs. 1mM $NH_4Cl$ | **** | <0,0001 |
| | | | ASW vs. 10mM $NH_4Cl$ | **** | <0,0001 |
| | | | ASW vs. 0.5mM $NH_4Cl$ | ns | 0,6723 |
| | | 210 min | ASW vs. 1mM $NH_4Cl$ | **** | <0,0001 |
| | | | ASW vs. 10mM $NH_4Cl$ | **** | <0,0001 |
| | | | ASW vs. 0.5mM $NH_4Cl$ | ns | 0,3920 |
| | | 240 min | ASW vs. 1mM $NH_4Cl$ | **** | <0,0001 |
| | | | ASW vs. 10mM $NH_4Cl$ | **** | <0,0001 |
| | | | ASW vs. 0.5mM $NH_4Cl$ | ns | 0,2493 |
| | | 270 min | ASW vs. 1mM $NH_4Cl$ | **** | <0,0001 |

| | | | | | |
|---|---|---|---|---|---|
| | | | ASW vs. 10mM $NH_4Cl$ | **** | <0,0001 |
| | | | ASW vs. 0.5mM $NH_4Cl$ | ns | 0,2217 |
| | | 300 min | ASW vs. 1mM $NH_4Cl$ | **** | <0,0001 |
| | | | ASW vs. 10mM $NH_4Cl$ | **** | <0,0001 |
| | | | ASW vs. 0.5mM $NH_4Cl$ | ns | 0,0640 |
| | | 330 min | ASW vs. 1mM $NH_4Cl$ | **** | <0,0001 |
| | | | ASW vs. 10mM $NH_4Cl$ | **** | <0,0001 |
| | | | ASW vs. 0.5mM $NH_4Cl$ | * | 0,0185 |
| | | 360 min | ASW vs. 1mM $NH_4Cl$ | *** | 0,0001 |
| | | | ASW vs. 10mM $NH_4Cl$ | **** | <0,0001 |
| | | | ASW vs. 0.5mM $NH_4Cl$ | * | 0,0405 |
| | | 390 min | ASW vs. 1mM $NH_4Cl$ | ** | 0,0042 |
| | | | ASW vs. 10mM $NH_4Cl$ | **** | <0,0001 |
| | | | ASW vs. 0.5mM $NH_4Cl$ | ns | 0,2236 |
| | | 420 min | ASW vs. 1mM $NH_4Cl$ | ns | 0,0816 |
| | | | ASW vs. 10mM $NH_4Cl$ | **** | <0,0001 |
| | | | ASW vs. 0.5mM $NH_4Cl$ | ns | 0,6709 |
| | | 450 min | ASW vs. 1mM $NH_4Cl$ | ns | 0,2156 |
| | | | ASW vs. 10mM $NH_4Cl$ | *** | 0,0005 |
| | | | ASW vs. 0.5mM $NH_4Cl$ | ns | 0,8645 |
| | | 480 min | ASW vs. 1mM $NH_4Cl$ | ns | 0,1425 |
| | | | ASW vs. 10mM $NH_4Cl$ | *** | 0,0003 |
| | | | ASW vs. 0.5mM $NH_4Cl$ | ns | 0,5901 |
| | | 510 min | ASW vs. 1mM $NH_4Cl$ | ns | 0,1912 |
| | | | ASW vs. 10mM $NH_4Cl$ | *** | 0,0008 |
| | | | ASW vs. 0.5mM $NH_4Cl$ | ns | 0,6155 |
| | | 540 min | ASW vs. 1mM $NH_4Cl$ | ns | 0,1034 |
| | | | ASW vs. 10mM $NH_4Cl$ | ** | 0,0023 |
| | | | ASW vs. 0.5mM $NH_4Cl$ | ns | 0,6075 |
| | | 570 min | ASW vs. 1mM $NH_4Cl$ | ns | 0,0957 |
| | | | ASW vs. 10mM $NH_4Cl$ | ** | 0,0090 |
| | | | ASW vs. 0.5mM $NH_4Cl$ | ns | 0,7725 |

| | | | | | |
|---|---|---|---|---|---|
| | | 600 min | ASW vs. 1mM $NH_4Cl$ | ns | 0,1924 |
| | | | ASW vs. 10mM $NH_4Cl$ | * | 0,0243 |
| | | | ASW vs. 0.5mM $NH_4Cl$ | ns | 0,9989 |

table S2 Statistics for settlement rates analysis in the presence of $(NH_4)_2SO_4$

| Figure & Panel | Test | Data point | Comparison | P Value summary | P Value |
|---|---|---|---|---|---|
| Fig. 1D | Two-way RM ANOVA | | | | |
| | | | Time | **** | <0,0001 |
| | | | Fraction Settled | **** | <0,0001 |
| | | | Time x Fraction Settled | **** | <0,0001 |
| | Dunnett's multiple comparisons test | | | | |
| | | 30 min | ASW vs. 0.5mM $(NH_4)_2SO_4$ | ns | 0,2038 |
| | | | ASW vs. 1mM $(NH_4)_2SO_4$ | *** | 0,0001 |
| | | | ASW vs. 10mM $(NH_4)_2SO_4$ | **** | <0,0001 |
| | | 60 min | ASW vs. 0.5mM $(NH_4)_2SO_4$ | ns | 0,2418 |
| | | | ASW vs. 1mM $(NH_4)_2SO_4$ | **** | <0,0001 |
| | | | ASW vs. 10mM $(NH_4)_2SO_4$ | **** | <0,0001 |
| | | 90 min | ASW vs. 0.5mM $(NH_4)_2SO_4$ | ns | 0,0980 |
| | | | ASW vs. 1mM $(NH_4)_2SO_4$ | **** | <0,0001 |
| | | | ASW vs. 10mM $(NH_4)_2SO_4$ | **** | <0,0001 |
| | | 120 min | ASW vs. 0.5mM $(NH_4)_2SO_4$ | * | 0,0483 |
| | | | ASW vs. 1mM $(NH_4)_2SO_4$ | **** | <0,0001 |
| | | | ASW vs. 10mM $(NH_4)_2SO_4$ | **** | <0,0001 |
| | | 150 min | ASW vs. 0.5mM $(NH_4)_2SO_4$ | ns | 0,2009 |
| | | | ASW vs. 1mM $(NH_4)_2SO_4$ | **** | <0,0001 |
| | | | ASW vs. 10mM $(NH_4)_2SO_4$ | **** | <0,0001 |
| | | 180 min | ASW vs. 0.5mM $(NH_4)_2SO_4$ | ns | 0,2826 |
| | | | ASW vs. 1mM $(NH_4)_2SO_4$ | *** | 0,0002 |
| | | | ASW vs. 10mM $(NH_4)_2SO_4$ | **** | <0,0001 |
| | | 210 min | ASW vs. 0.5mM $(NH_4)_2SO_4$ | ns | 0,1188 |
| | | | ASW vs. 1mM $(NH_4)_2SO_4$ | *** | 0,0007 |
| | | | ASW vs. 10mM $(NH_4)_2SO_4$ | **** | <0,0001 |
| | | 240 min | ASW vs. 0.5mM $(NH_4)_2SO_4$ | ns | 0,1157 |

| | | | | | |
|---|---|---|---|---|---|
| | | | ASW vs. 1mM $(NH_4)_2SO_4$ | *** | 0,0005 |
| | | | ASW vs. 10mM $(NH_4)_2SO_4$ | **** | <0,0001 |
| | | 270 min | ASW vs. 0.5mM $(NH_4)_2SO_4$ | ns | 0,0727 |
| | | | ASW vs. 1mM $(NH_4)_2SO_4$ | **** | <0,0001 |
| | | | ASW vs. 10mM $(NH_4)_2SO_4$ | **** | <0,0001 |
| | | 300 min | ASW vs. 0.5mM $(NH_4)_2SO_4$ | * | 0,0394 |
| | | | ASW vs. 1mM $(NH_4)_2SO_4$ | *** | 0,0001 |
| | | | ASW vs. 10mM $(NH_4)_2SO_4$ | **** | <0,0001 |
| | | 330 min | ASW vs. 0.5mM $(NH_4)_2SO_4$ | * | 0,0337 |
| | | | ASW vs. 1mM $(NH_4)_2SO_4$ | *** | 0,0007 |
| | | | ASW vs. 10mM $(NH_4)_2SO_4$ | *** | 0,0004 |
| | | 360 min | ASW vs. 0.5mM $(NH_4)_2SO_4$ | ns | 0,3465 |
| | | | ASW vs. 1mM $(NH_4)_2SO_4$ | * | 0,0215 |
| | | | ASW vs. 10mM $(NH_4)_2SO_4$ | * | 0,0164 |
| | | 390 min | ASW vs. 0.5mM $(NH_4)_2SO_4$ | ns | 0,9483 |
| | | | ASW vs. 1mM $(NH_4)_2SO_4$ | * | 0,0238 |
| | | | ASW vs. 10mM $(NH_4)_2SO_4$ | * | 0,0498 |
| | | 420 min | ASW vs. 0.5mM $(NH_4)_2SO_4$ | ns | 0,9974 |
| | | | ASW vs. 1mM $(NH_4)_2SO_4$ | ns | 0,0512 |
| | | | ASW vs. 10mM $(NH_4)_2SO_4$ | ns | 0,0976 |
| | | 450 min | ASW vs. 0.5mM $(NH_4)_2SO_4$ | ns | 0,9883 |
| | | | ASW vs. 1mM $(NH_4)_2SO_4$ | * | 0,0413 |
| | | | ASW vs. 10mM $(NH_4)_2SO_4$ | * | 0,0369 |
| | | 480 min | ASW vs. 0.5mM $(NH_4)_2SO_4$ | ns | 0,9382 |
| | | | ASW vs. 1mM $(NH_4)_2SO_4$ | * | 0,0200 |
| | | | ASW vs. 10mM $(NH_4)_2SO_4$ | * | 0,0220 |
| | | 510 min | ASW vs. 0.5mM $(NH_4)_2SO_4$ | ns | 0,9362 |
| | | | ASW vs. 1mM $(NH_4)_2SO_4$ | * | 0,0176 |
| | | | ASW vs. 10mM $(NH_4)_2SO_4$ | * | 0,0173 |
| | | 540 min | ASW vs. 0.5mM $(NH_4)_2SO_4$ | ns | 0,8610 |
| | | | ASW vs. 1mM $(NH_4)_2SO_4$ | * | 0,0123 |
| | | | ASW vs. 10mM $(NH_4)_2SO_4$ | * | 0,0135 |

| | | | | | |
|---|---|---|---|---|---|
| | | 570 min | ASW vs. 0.5mM $(NH_4)_2SO_4$ | ns | 0,9961 |
| | | | ASW vs. 1mM $(NH_4)_2SO_4$ | ** | 0,0060 |
| | | | ASW vs. 10mM $(NH_4)_2SO_4$ | * | 0,0103 |
| | | 600 min | ASW vs. 0.5mM $(NH_4)_2SO_4$ | ns | 0,9464 |
| | | | ASW vs. 1mM $(NH_4)_2SO_4$ | ns | 0,4877 |
| | | | ASW vs. 10mM $(NH_4)_2SO_4$ | ns | 0,2327 |

table S3 Statistics for settlement rates analysis in the presence of CsCl

| Figure & Panel | Test | Data point | Comparison | P Value summary | P Value |
|---|---|---|---|---|---|
| Fig. 1E | Two-way RM ANOVA | | | | |
| | | | Time | **** | <0,0001 |
| | | | Fraction Settled | **** | <0,0001 |
| | | | Time x Fraction Settled | **** | <0,0001 |
| | Dunnett's multiple comparisons test | | | | |
| | | 30 min | ASW vs. 1mM CsCl | ns | 0,9042 |
| | | | ASW vs. 10mM CsCl | *** | 0,0005 |
| | | | ASW vs. 20mM CsCl | **** | <0,0001 |
| | | 60 min | ASW vs. 1mM CsCl | ** | 0,0020 |
| | | | ASW vs. 10mM CsCl | **** | <0,0001 |
| | | | ASW vs. 20mM CsCl | **** | <0,0001 |
| | | 90 min | ASW vs. 1mM CsCl | ** | 0,0019 |
| | | | ASW vs. 10mM CsCl | **** | <0,0001 |
| | | | ASW vs. 20mM CsCl | **** | <0,0001 |
| | | 120 min | ASW vs. 1mM CsCl | ns | 0,0670 |
| | | | ASW vs. 10mM CsCl | *** | 0,0001 |
| | | | ASW vs. 20mM CsCl | **** | <0,0001 |
| | | 150 min | ASW vs. 1mM CsCl | * | 0,0166 |
| | | | ASW vs. 10mM CsCl | *** | 0,0002 |
| | | | ASW vs. 20mM CsCl | **** | <0,0001 |
| | | 180 min | ASW vs. 1mM CsCl | * | 0,0156 |
| | | | ASW vs. 10mM CsCl | *** | 0,0003 |
| | | | ASW vs. 20mM CsCl | **** | <0,0001 |
| | | 210 min | ASW vs. 1mM CsCl | * | 0,0105 |

| | | | | | |
|---|---|---|---|---|---|
| | | | ASW vs. 10mM CsCl | *** | 0,0006 |
| | | | ASW vs. 20mM CsCl | **** | <0,0001 |
| | | 240 min | ASW vs. 1mM CsCl | ** | 0,0029 |
| | | | ASW vs. 10mM CsCl | *** | 0,0008 |
| | | | ASW vs. 20mM CsCl | *** | 0,0001 |
| | | 270 min | ASW vs. 1mM CsCl | ** | 0,0095 |
| | | | ASW vs. 10mM CsCl | ** | 0,0017 |
| | | | ASW vs. 20mM CsCl | *** | 0,0006 |
| | | 300 min | ASW vs. 1mM CsCl | * | 0,0241 |
| | | | ASW vs. 10mM CsCl | ** | 0,0019 |
| | | | ASW vs. 20mM CsCl | *** | 0,0009 |
| | | 330 min | ASW vs. 1mM CsCl | * | 0,0144 |
| | | | ASW vs. 10mM CsCl | ** | 0,0033 |
| | | | ASW vs. 20mM CsCl | ** | 0,0020 |
| | | 360 min | ASW vs. 1mM CsCl | * | 0,0104 |
| | | | ASW vs. 10mM CsCl | ** | 0,0017 |
| | | | ASW vs. 20mM CsCl | ** | 0,0021 |
| | | 390 min | ASW vs. 1mM CsCl | * | 0,0215 |
| | | | ASW vs. 10mM CsCl | * | 0,0219 |
| | | | ASW vs. 20mM CsCl | * | 0,0221 |
| | | 420 min | ASW vs. 1mM CsCl | ns | 0,1010 |
| | | | ASW vs. 10mM CsCl | ns | 0,1104 |
| | | | ASW vs. 20mM CsCl | ns | 0,1278 |
| | | 450 min | ASW vs. 1mM CsCl | ns | 0,1009 |
| | | | ASW vs. 10mM CsCl | ns | 0,1430 |
| | | | ASW vs. 20mM CsCl | ns | 0,2066 |
| | | 480 min | ASW vs. 1mM CsCl | ns | 0,0832 |
| | | | ASW vs. 10mM CsCl | ns | 0,1966 |
| | | | ASW vs. 20mM CsCl | ns | 0,2746 |
| | | 510 min | ASW vs. 1mM CsCl | ns | 0,5000 |
| | | | ASW vs. 10mM CsCl | ns | 0,9517 |
| | | | ASW vs. 20mM CsCl | ns | 0,9576 |

| | | | | | |
|---|---|---|---|---|---|
| | | 540 min | ASW vs. 1mM CsCl | ns | 0,2914 |
| | | | ASW vs. 10mM CsCl | ns | 0,7819 |
| | | | ASW vs. 20mM CsCl | ns | 0,7980 |
| | | 570 min | ASW vs. 1mM CsCl | ns | 0,2895 |
| | | | ASW vs. 10mM CsCl | ns | 0,7100 |
| | | | ASW vs. 20mM CsCl | ns | 0,7830 |
| | | 600 min | ASW vs. 1mM CsCl | ns | 0,2895 |
| | | | ASW vs. 10mM CsCl | ns | 0,5580 |
| | | | ASW vs. 20mM CsCl | ns | 0,5923 |

table S4 Statistics for settlement rates analysis in the presence of KCl

| Figure & Panel | Test | Data point | Comparison | P Value summary | P Value |
|---|---|---|---|---|---|
| Fig. 1F | Two-way RM ANOVA | | | | |
| | | | Time | **** | <0,0001 |
| | | | Fraction Settled | **** | <0,0001 |
| | | | Time x Fraction Settled | **** | <0,0001 |
| | Dunnett's multiple comparisons test | | | | |
| | | 30 min | ASW vs. 0.5mM KCl | ns | 0,9846 |
| | | | ASW vs. 1mM KCl | *** | 0,0007 |
| | | | ASW vs. 10mM KCl | *** | 0,0005 |
| | | 60 min | ASW vs. 0.5mM KCl | ns | 0,8665 |
| | | | ASW vs. 1mM KCl | ** | 0,0022 |
| | | | ASW vs. 10mM KCl | *** | 0,0004 |
| | | 90 min | ASW vs. 0.5mM KCl | ns | 0,9952 |
| | | | ASW vs. 1mM KCl | ** | 0,0039 |
| | | | ASW vs. 10mM KCl | *** | 0,0002 |
| | | 120 min | ASW vs. 0.5mM KCl | ns | 0,5881 |
| | | | ASW vs. 1mM KCl | *** | 0,0003 |
| | | | ASW vs. 10mM KCl | **** | <0,0001 |
| | | 150 min | ASW vs. 0.5mM KCl | ns | 0,1259 |
| | | | ASW vs. 1mM KCl | *** | 0,0005 |
| | | | ASW vs. 10mM KCl | **** | <0,0001 |

| | | | | | |
|---|---|---|---|---|---|
| | | 180 min | ASW vs. 0.5mM KCl | * | 0,0410 |
| | | | ASW vs. 1mM KCl | **** | <0,0001 |
| | | | ASW vs. 10mM KCl | **** | <0,0001 |
| | | 210 min | ASW vs. 0.5mM KCl | * | 0,0102 |
| | | | ASW vs. 1mM KCl | **** | <0,0001 |
| | | | ASW vs. 10mM KCl | **** | <0,0001 |
| | | 240 min | ASW vs. 0.5mM KCl | * | 0,0184 |
| | | | ASW vs. 1mM KCl | *** | 0,0001 |
| | | | ASW vs. 10mM KCl | **** | <0,0001 |
| | | 270 min | ASW vs. 0.5mM KCl | * | 0,0173 |
| | | | ASW vs. 1mM KCl | **** | <0,0001 |
| | | | ASW vs. 10mM KCl | **** | <0,0001 |
| | | 300 min | ASW vs. 0.5mM KCl | ** | 0,0036 |
| | | | ASW vs. 1mM KCl | **** | <0,0001 |
| | | | ASW vs. 10mM KCl | **** | <0,0001 |
| | | 330 min | ASW vs. 0.5mM KCl | **** | <0,0001 |
| | | | ASW vs. 1mM KCl | **** | <0,0001 |
| | | | ASW vs. 10mM KCl | **** | <0,0001 |
| | | 360 min | ASW vs. 0.5mM KCl | ** | 0,0031 |
| | | | ASW vs. 1mM KCl | *** | 0,0003 |
| | | | ASW vs. 10mM KCl | **** | <0,0001 |
| | | 390 min | ASW vs. 0.5mM KCl | * | 0,0495 |
| | | | ASW vs. 1mM KCl | ** | 0,0037 |
| | | | ASW vs. 10mM KCl | *** | 0,0003 |
| | | 420 min | ASW vs. 0.5mM KCl | ns | 0,0966 |
| | | | ASW vs. 1mM KCl | ** | 0,0054 |
| | | | ASW vs. 10mM KCl | *** | 0,0005 |
| | | 450 min | ASW vs. 0.5mM KCl | ns | 0,1143 |
| | | | ASW vs. 1mM KCl | *** | 0,0005 |
| | | | ASW vs. 10mM KCl | **** | <0,0001 |
| | | 480 min | ASW vs. 0.5mM KCl | ns | 0,0655 |
| | | | ASW vs. 1mM KCl | *** | 0,0007 |

| | | | | | |
|---|---|---|---|---|---|
| | | | ASW vs. 10mM KCl | **** | <0,0001 |
| | | 510 min | ASW vs. 0.5mM KCl | ns | 0,1566 |
| | | | ASW vs. 1mM KCl | ** | 0,0031 |
| | | | ASW vs. 10mM KCl | ** | 0,0080 |
| | | 540 min | ASW vs. 0.5mM KCl | ns | 0,5120 |
| | | | ASW vs. 1mM KCl | ns | 0,2011 |
| | | | ASW vs. 10mM KCl | ns | 0,1699 |
| | | 570 min | ASW vs. 0.5mM KCl | ns | 0,9154 |
| | | | ASW vs. 1mM KCl | ns | 0,3661 |
| | | | ASW vs. 10mM KCl | ns | 0,3661 |
| | | 600 min | ASW vs. 0.5mM KCl | ns | 0,9154 |
| | | | ASW vs. 1mM KCl | ns | 0,3661 |
| | | | ASW vs. 10mM KCl | ns | 0,3661 |

table S5 Statistics for settlement rates analysis in the presence of sugars

| Figure & Panel | Test | Data point | Comparison | P Value summary | P Value |
|---|---|---|---|---|---|
| Fig. 1G | Two-way RM ANOVA | | | | |
| | | | Time | **** | <0,0001 |
| | | | Fraction Settled | **** | <0,0001 |
| | | | Time x Fraction Settled | **** | <0,0001 |
| | Dunnett's multiple comparisons test | | | | |
| | | 30 min | ASW vs. sucrose | *** | 0,0001 |
| | | | ASW vs. glucose | ** | 0,0043 |
| | | 60 min | ASW vs. sucrose | **** | <0,0001 |
| | | | ASW vs. glucose | **** | <0,0001 |
| | | 90 min | ASW vs. sucrose | **** | <0,0001 |
| | | | ASW vs. glucose | **** | <0,0001 |
| | | 120 min | ASW vs. sucrose | **** | <0,0001 |
| | | | ASW vs. glucose | **** | <0,0001 |
| | | 150 min | ASW vs. sucrose | **** | <0,0001 |
| | | | ASW vs. glucose | **** | <0,0001 |
| | | 180 min | ASW vs. sucrose | **** | <0,0001 |

| | | | | | |
|---|---|---|---|---|---|
| | | | ASW vs. glucose | *** | 0,0003 |
| | | 210 min | ASW vs. sucrose | **** | <0,0001 |
| | | | ASW vs. glucose | *** | 0,0008 |
| | | 240 min | ASW vs. sucrose | **** | <0,0001 |
| | | | ASW vs. glucose | ** | 0,0031 |
| | | 270 min | ASW vs. sucrose | **** | <0,0001 |
| | | | ASW vs. glucose | ns | 0,1221 |
| | | 300 min | ASW vs. sucrose | ** | 0,0018 |
| | | | ASW vs. glucose | ns | 0,1979 |
| | | 330 min | ASW vs. sucrose | ** | 0,0033 |
| | | | ASW vs. glucose | ns | 0,0730 |
| | | 360 min | ASW vs. sucrose | ** | 0,0061 |
| | | | ASW vs. glucose | ns | 0,1013 |
| | | 390 min | ASW vs. sucrose | * | 0,0111 |
| | | | ASW vs. glucose | ns | 0,0612 |
| | | 420 min | ASW vs. sucrose | ns | 0,1952 |
| | | | ASW vs. glucose | ns | 0,8286 |
| | | 450 min | ASW vs. sucrose | ns | 0,4644 |
| | | | ASW vs. glucose | ns | 0,8809 |
| | | 480 min | ASW vs. sucrose | ns | 0,3596 |
| | | | ASW vs. glucose | ns | 0,5807 |
| | | 510 min | ASW vs. sucrose | ns | 0,3466 |
| | | | ASW vs. glucose | ns | 0,2464 |
| | | 540 min | ASW vs. sucrose | ns | 0,1634 |
| | | | ASW vs. glucose | ns | 0,1634 |
| | | 570 min | ASW vs. sucrose | ns | 0,1887 |
| | | | ASW vs. glucose | ns | 0,1887 |
| | | 600 min | ASW vs. sucrose | ns | 0,1887 |
| | | | ASW vs. glucose | ns | 0,1887 |

table S6 Statistics for settlement rates analysis in the presence of short alcohols

| Figure & Panel | Test | Data point | Comparison | P Value summary | P Value |
|---|---|---|---|---|---|
| Fig. 1H | Two-way RM ANOVA | | | | |
| | | | Time | **** | <0,0001 |
| | | | Fraction Settled | *** | 0,0002 |
| | | | Time x Fraction Settled | **** | <0,0001 |
| | Dunnett's multiple comparisons test | | | | |
| | | 30 min | ASW vs. butanol | ** | 0,0091 |
| | | | ASW vs. isoamyl acetate | * | 0,0490 |
| | | 60 min | ASW vs. butanol | *** | 0,0004 |
| | | | ASW vs. isoamyl acetate | **** | <0,0001 |
| | | 90 min | ASW vs. butanol | ** | 0,0011 |
| | | | ASW vs. isoamyl acetate | *** | 0,0001 |
| | | 120 min | ASW vs. butanol | *** | 0,0007 |
| | | | ASW vs. isoamyl acetate | **** | <0,0001 |
| | | 150 min | ASW vs. butanol | ** | 0,0011 |
| | | | ASW vs. isoamyl acetate | ** | 0,0016 |
| | | 180 min | ASW vs. butanol | ** | 0,0017 |
| | | | ASW vs. isoamyl acetate | ** | 0,0026 |
| | | 210 min | ASW vs. butanol | *** | 0,0009 |
| | | | ASW vs. isoamyl acetate | ** | 0,0029 |
| | | 240 min | ASW vs. butanol | ** | 0,0040 |
| | | | ASW vs. isoamyl acetate | * | 0,0158 |
| | | 270 min | ASW vs. butanol | * | 0,0174 |
| | | | ASW vs. isoamyl acetate | * | 0,0228 |
| | | 300 min | ASW vs. butanol | ns | 0,0568 |
| | | | ASW vs. isoamyl acetate | ns | 0,0743 |
| | | 330 min | ASW vs. butanol | ns | 0,0550 |
| | | | ASW vs. isoamyl acetate | * | 0,0275 |
| | | 360 min | ASW vs. butanol | ns | 0,1496 |
| | | | ASW vs. isoamyl acetate | ns | 0,3566 |

| | | | | | |
|---|---|---|---|---|---|
| | | 390 min | ASW vs. butanol | ns | 0,0748 |
| | | | ASW vs. isoamyl acetate | ns | 0,2789 |
| | | 420 min | ASW vs. butanol | ns | 0,1356 |
| | | | ASW vs. isoamyl acetate | ns | 0,0626 |
| | | 450 min | ASW vs. butanol | ns | 0,2685 |
| | | | ASW vs. isoamyl acetate | ns | 0,0915 |
| | | 480 min | ASW vs. butanol | ns | 0,4721 |
| | | | ASW vs. isoamyl acetate | ns | 0,4112 |
| | | 510 min | ASW vs. butanol | ns | 0,3203 |
| | | | ASW vs. isoamyl acetate | ns | 0,4314 |
| | | 540 min | ASW vs. butanol | ns | 0,4644 |
| | | | ASW vs. isoamyl acetate | ns | 0,5981 |
| | | 570 min | ASW vs. butanol | ns | 0,8049 |
| | | | ASW vs. isoamyl acetate | ns | 0,8049 |
| | | 600 min | ASW vs. butanol | ns | 0,8049 |
| | | | ASW vs. isoamyl acetate | ns | >0,9999 |

table S7 Statistics for settlement rates analysis in the presence of amino acids

| Figure & Panel | Test | Data point | Comparison | P Value summary | P Value |
|---|---|---|---|---|---|
| Fig. 1I | Two-way RM ANOVA | | | | |
| | | | Time | **** | <0,0001 |
| | | | Fraction Settled | **** | <0,0001 |
| | | | Time x Fraction Settled | **** | <0,0001 |
| | Dunnett's multiple comparisons test | | | | |
| | | 30 min | ASW vs. Alanine | * | 0,0433 |
| | | | ASW vs. Glutamate | **** | <0,0001 |
| | | | ASW vs. Methionine | ns | 0,1148 |
| | | | ASW vs. Leucine | ns | 0,6471 |
| | | | ASW vs. Asparagine | ns | 0,9541 |
| | | 60 min | ASW vs. Alanine | * | 0,0116 |
| | | | ASW vs. Glutamate | **** | <0,0001 |

| | | | | | |
|---|---|---|---|---|---|
| | | | ASW vs. Methionine | ns | 0,1232 |
| | | | ASW vs. Leucine | ns | 0,8253 |
| | | | ASW vs. Asparagine | ns | 0,4736 |
| | | 90 min | ASW vs. Alanine | *** | 0,0006 |
| | | | ASW vs. Glutamate | **** | <0,0001 |
| | | | ASW vs. Methionine | * | 0,0207 |
| | | | ASW vs. Leucine | ns | 0,9973 |
| | | | ASW vs. Asparagine | ns | 0,4712 |
| | | 120 min | ASW vs. Alanine | *** | 0,0006 |
| | | | ASW vs. Glutamate | **** | <0,0001 |
| | | | ASW vs. Methionine | *** | 0,0007 |
| | | | ASW vs. Leucine | ns | 0,4978 |
| | | | ASW vs. Asparagine | ns | 0,4712 |
| | | 150 min | ASW vs. Alanine | *** | 0,0005 |
| | | | ASW vs. Glutamate | ** | 0,0020 |
| | | | ASW vs. Methionine | *** | 0,0010 |
| | | | ASW vs. Leucine | ns | 0,4303 |
| | | | ASW vs. Asparagine | ns | 0,4945 |
| | | 180 min | ASW vs. Alanine | *** | 0,0003 |
| | | | ASW vs. Glutamate | ** | 0,0055 |
| | | | ASW vs. Methionine | ** | 0,0021 |
| | | | ASW vs. Leucine | ns | 0,4221 |
| | | | ASW vs. Asparagine | ns | 0,5140 |
| | | 210 min | ASW vs. Alanine | *** | 0,0004 |
| | | | ASW vs. Glutamate | ** | 0,0017 |
| | | | ASW vs. Methionine | *** | 0,0002 |
| | | | ASW vs. Leucine | ns | 0,0623 |
| | | | ASW vs. Asparagine | ns | 0,6135 |
| | | 240 min | ASW vs. Alanine | *** | 0,0007 |
| | | | ASW vs. Glutamate | ** | 0,0042 |
| | | | ASW vs. Methionine | **** | <0,0001 |
| | | | ASW vs. Leucine | ** | 0,0031 |

| | | | | | |
|---|---|---|---|---|---|
| | | | ASW vs. Asparagine | ns | 0,6683 |
| | | 270 min | ASW vs. Alanine | * | 0,0137 |
| | | | ASW vs. Glutamate | * | 0,0439 |
| | | | ASW vs. Methionine | **** | <0,0001 |
| | | | ASW vs. Leucine | *** | 0,0008 |
| | | | ASW vs. Asparagine | ns | 0,8659 |
| | | 300 min | ASW vs. Alanine | * | 0,0261 |
| | | | ASW vs. Glutamate | ns | 0,2465 |
| | | | ASW vs. Methionine | **** | <0,0001 |
| | | | ASW vs. Leucine | **** | <0,0001 |
| | | | ASW vs. Asparagine | ns | 0,9997 |
| | | 330 min | ASW vs. Alanine | ns | 0,0773 |
| | | | ASW vs. Glutamate | ns | 0,7365 |
| | | | ASW vs. Methionine | **** | <0,0001 |
| | | | ASW vs. Leucine | **** | <0,0001 |
| | | | ASW vs. Asparagine | ns | 0,8744 |
| | | 360 min | ASW vs. Alanine | * | 0,0286 |
| | | | ASW vs. Glutamate | ns | 0,7148 |
| | | | ASW vs. Methionine | **** | <0,0001 |
| | | | ASW vs. Leucine | *** | 0,0003 |
| | | | ASW vs. Asparagine | ns | 0,6528 |
| | | 390 min | ASW vs. Alanine | ns | 0,2666 |
| | | | ASW vs. Glutamate | ns | 0,8879 |
| | | | ASW vs. Methionine | **** | <0,0001 |
| | | | ASW vs. Leucine | **** | <0,0001 |
| | | | ASW vs. Asparagine | ns | 0,2940 |
| | | 420 min | ASW vs. Alanine | ns | 0,3234 |
| | | | ASW vs. Glutamate | ns | 0,9997 |
| | | | ASW vs. Methionine | **** | <0,0001 |
| | | | ASW vs. Leucine | *** | 0,0002 |
| | | | ASW vs. Asparagine | ns | 0,5101 |
| | | 450 min | ASW vs. Alanine | ns | 0,3682 |

| | | | | | |
|---|---|---|---|---|---|
| | | | ASW vs. Glutamate | ns | 0,9998 |
| | | | ASW vs. Methionine | **** | <0,0001 |
| | | | ASW vs. Leucine | *** | 0,0001 |
| | | | ASW vs. Asparagine | ns | 0,5803 |
| | | 480 min | ASW vs. Alanine | ns | 0,2378 |
| | | | ASW vs. Glutamate | ns | 0,9997 |
| | | | ASW vs. Methionine | **** | <0,0001 |
| | | | ASW vs. Leucine | *** | 0,0001 |
| | | | ASW vs. Asparagine | ns | 0,4897 |
| | | 510 min | ASW vs. Alanine | ns | 0,1690 |
| | | | ASW vs. Glutamate | ns | 0,9760 |
| | | | ASW vs. Methionine | **** | <0,0001 |
| | | | ASW vs. Leucine | **** | <0,0001 |
| | | | ASW vs. Asparagine | ns | 0,5093 |
| | | 540 min | ASW vs. Alanine | ns | 0,5881 |
| | | | ASW vs. Glutamate | ns | 0,9957 |
| | | | ASW vs. Methionine | **** | <0,0001 |
| | | | ASW vs. Leucine | *** | 0,0001 |
| | | | ASW vs. Asparagine | ns | 0,4358 |
| | | 570 min | ASW vs. Alanine | ns | 0,9950 |
| | | | ASW vs. Glutamate | ns | 0,4014 |
| | | | ASW vs. Methionine | **** | <0,0001 |
| | | | ASW vs. Leucine | *** | 0,0002 |
| | | | ASW vs. Asparagine | ns | 0,2948 |
| | | 600 min | ASW vs. Alanine | ns | 0,9950 |
| | | | ASW vs. Glutamate | ns | 0,5823 |
| | | | ASW vs. Methionine | **** | <0,0001 |
| | | | ASW vs. Leucine | *** | 0,0005 |
| | | | ASW vs. Asparagine | ns | 0,9682 |

table S8 Statistics for settlement rates analysis in the presence of terpenes

| Figure & Panel | Test | Data point | Comparison | P Value summary | P Value |
|---|---|---|---|---|---|
| Fig. 1J | Two-way RM ANOVA | | | | |
| | | | Time | **** | <0,0001 |
| | | | Fraction Settled | **** | <0,0001 |
| | | | Time x Fraction Settled | **** | <0,0001 |
| | Dunnett's multiple comparisons test | | | | |
| | | 30 min | ASW vs. Carvacrol | * | 0,0136 |
| | | | ASW vs. Nootkatone | * | 0,0477 |
| | | | ASW vs. Eucalyptol | * | 0,0357 |
| | | 60 min | ASW vs. Carvacrol | * | 0,0147 |
| | | | ASW vs. Nootkatone | ** | 0,0086 |
| | | | ASW vs. Eucalyptol | ** | 0,0025 |
| | | 90 min | ASW vs. Carvacrol | *** | 0,0005 |
| | | | ASW vs. Nootkatone | **** | <0,0001 |
| | | | ASW vs. Eucalyptol | **** | <0,0001 |
| | | 120 min | ASW vs. Carvacrol | *** | 0,0001 |
| | | | ASW vs. Nootkatone | **** | <0,0001 |
| | | | ASW vs. Eucalyptol | **** | <0,0001 |
| | | 150 min | ASW vs. Carvacrol | **** | <0,0001 |
| | | | ASW vs. Nootkatone | **** | <0,0001 |
| | | | ASW vs. Eucalyptol | **** | <0,0001 |
| | | 180 min | ASW vs. Carvacrol | **** | <0,0001 |
| | | | ASW vs. Nootkatone | **** | <0,0001 |
| | | | ASW vs. Eucalyptol | **** | <0,0001 |
| | | 210 min | ASW vs. Carvacrol | **** | <0,0001 |
| | | | ASW vs. Nootkatone | **** | <0,0001 |
| | | | ASW vs. Eucalyptol | **** | <0,0001 |
| | | 240 min | ASW vs. Carvacrol | **** | <0,0001 |
| | | | ASW vs. Nootkatone | **** | <0,0001 |
| | | | ASW vs. Eucalyptol | **** | <0,0001 |
| | | 270 min | ASW vs. Carvacrol | **** | <0,0001 |

| | | | | | |
|---|---|---|---|---|---|
| | | | ASW vs. Nootkatone | **** | <0,0001 |
| | | | ASW vs. Eucalyptol | **** | <0,0001 |
| | | 300 min | ASW vs. Carvacrol | **** | <0,0001 |
| | | | ASW vs. Nootkatone | **** | <0,0001 |
| | | | ASW vs. Eucalyptol | **** | <0,0001 |
| | | 330 min | ASW vs. Carvacrol | **** | <0,0001 |
| | | | ASW vs. Nootkatone | **** | <0,0001 |
| | | | ASW vs. Eucalyptol | **** | <0,0001 |
| | | 360 min | ASW vs. Carvacrol | **** | <0,0001 |
| | | | ASW vs. Nootkatone | **** | <0,0001 |
| | | | ASW vs. Eucalyptol | **** | <0,0001 |
| | | 390 min | ASW vs. Carvacrol | **** | <0,0001 |
| | | | ASW vs. Nootkatone | **** | <0,0001 |
| | | | ASW vs. Eucalyptol | **** | <0,0001 |
| | | 420 min | ASW vs. Carvacrol | **** | <0,0001 |
| | | | ASW vs. Nootkatone | **** | <0,0001 |
| | | | ASW vs. Eucalyptol | **** | <0,0001 |
| | | 450 min | ASW vs. Carvacrol | **** | <0,0001 |
| | | | ASW vs. Nootkatone | **** | <0,0001 |
| | | | ASW vs. Eucalyptol | **** | <0,0001 |
| | | 480 min | ASW vs. Carvacrol | **** | <0,0001 |
| | | | ASW vs. Nootkatone | **** | <0,0001 |
| | | | ASW vs. Eucalyptol | **** | <0,0001 |
| | | 510 min | ASW vs. Carvacrol | **** | <0,0001 |
| | | | ASW vs. Nootkatone | **** | <0,0001 |
| | | | ASW vs. Eucalyptol | **** | <0,0001 |
| | | 540 min | ASW vs. Carvacrol | **** | <0,0001 |
| | | | ASW vs. Nootkatone | **** | <0,0001 |
| | | | ASW vs. Eucalyptol | **** | <0,0001 |
| | | 570 min | ASW vs. Carvacrol | **** | <0,0001 |
| | | | ASW vs. Nootkatone | **** | <0,0001 |
| | | | ASW vs. Eucalyptol | **** | <0,0001 |

| | | | | | |
|---|---|---|---|---|---|
| | | 600 min | ASW vs. Carvacrol | **** | <0,0001 |
| | | | ASW vs. Nootkatone | **** | <0,0001 |
| | | | ASW vs. Eucalyptol | **** | <0,0001 |

table S9 Statistics for tail regression (metamorphosis) assays with different chemical cues

| Figure & Panel | Test | Comparison | P Value summary | P Value |
|---|---|---|---|---|
| Fig. 1L | Kruskal-Wallis test | | **** | <0,0001 |
| | Dunn's multiple comparisons test | | | |
| | | ASW vs. 0.5mM NH4Cl | ** | 0,0019 |
| | | ASW vs. 1mM NH4Cl | **** | <0,0001 |
| | | ASW vs. 10mM NH4Cl | **** | <0,0001 |
| | | ASW vs. 0.5mM (NH4)2S04 | * | 0,0181 |
| | | ASW vs. 1mM (NH4)2S04 | **** | <0,0001 |
| | | ASW vs. 10mM (NH4)2S04 | **** | <0,0001 |
| | | ASW vs. 1mM CsCl | **** | <0,0001 |
| | | ASW vs. 10mM CsCl | **** | <0,0001 |
| | | ASW vs. 20mM CsCl | **** | <0,0001 |
| | | ASW vs. 0.5mM KCl | ns | 0,1787 |
| | | ASW vs. 1mM KCl | **** | <0,0001 |
| | | ASW vs. 10mM KCl | **** | <0,0001 |
| | | ASW vs. L-Alanine | ** | 0,0014 |
| | | ASW vs. 10mM L-Glutamate | **** | <0,0001 |
| | | ASW vs. Asparagine | ns | 0,2838 |
| | | ASW vs. L-Leucine | ns | >0,9999 |
| | | ASW vs. L-Methaionine | ns | >0,9999 |
| | | ASW vs. Carvacrol | * | 0,0308 |
| | | ASW vs. Eucalyptol | * | 0,0435 |
| | | ASW vs. Nootkatone | ns | >0,9999 |
| | | ASW vs. Butanol | **** | <0,0001 |
| | | ASW vs. isoamyl alcohol | * | 0,0406 |
| | | ASW vs. Sucrose | **** | <0,0001 |
| | | ASW vs. Glucose | **** | <0,0001 |

table S10 Time in minutes to reach 50% of settled larvae for each stimulus.

| Panel | Stimulus | Time at 50% (minutes) |
|---|---|---|
| Fig. 1C | ASW | 239.2 |
| | 0.5mM $NH_4Cl$ | 222.8 |
| | 1mM $NH_4Cl$ | 175.6 |
| | 10mM $NH_4Cl$ | 156.2 |
| Fig. 1D | ASW | 236.0 |
| | 0.5mM $(NH_4)_2SO_4$ | 209.6 |
| | 1mM $(NH_4)_2SO_4$ | 177.8 |
| | 10mM $(NH_4)_2SO_4$ | 151.0 |
| Fig. 1E | ASW | 247.8 |
| | 1mM CsCl | 204.5 |
| | 10mM CsCl | 187.4 |
| | 20mM CsCl | 177.0 |
| Fig. 1F | ASW | 251.2 |
| | 0.5mM KCl | 207.1 |
| | 1mM KCl | 172.8 |
| | 10mM KCl | 154.8 |
| Fig. 1G | ASW | 244.7 |
| | 100µM Sucrose | 167.4 |
| | 100µM Glucose | 267.5 |
| Fig. 1H | ASW | 243.7 |
| | 10µM Butanol | 174.1 |
| | 10µM Isoamyl alcohol | 224.4 |
| Fig. 1I | ASW | 237.4 |
| | 1mM Alanine | 172.4 |
| | 1mM Glutamate | 213.2 |
| | 1mM Methionine | 382.5 |
| | 1mM Leucine | 366.2 |
| | 1mM Asparagine | 235.2 |
| Fig. 1J | ASW | 220.3 |
| | 200µM Carvacrol | Not applicable |
| | 200µM Eucalyptol | Not applicable |
| | 200µM Nootkatone | Not applicable |

table S11 Statistics for settlement rates analysis in the presence of noxious cues

| Figure & Panel | Test | Data point | Comparison | P Value summary | P Value |
|---|---|---|---|---|---|
| S1A | Two-way RM ANOVA | | | | |
| | | | Time | **** | <0,0001 |
| | | | Fraction Settled | **** | <0,0001 |
| | | | Time x Fraction Settled | **** | <0,0001 |
| | Dunnett's multiple comparisons test | | | | |

| | | | | | |
|---|---|---|---|---|---|
| | | 30 min | ASW vs. 0.1 SDS | ** | 0,0014 |
| | | | ASW vs. 100mM $NH_4Cl$ | ** | 0,0090 |
| | | | ASW vs. 10mM Chloroquine | *** | 0,0007 |
| | | | ASW vs. 1mM $CuSO_4$ | *** | 0,0009 |
| | | 60 min | ASW vs. 0.1% SDS | **** | <0,0001 |
| | | | ASW vs. 100mM $NH_4Cl$ | ** | 0,0010 |
| | | | ASW vs. 10mM Chloroquine | **** | <0,0001 |
| | | | ASW vs. 1mM $CuSO_4$ | **** | <0,0001 |
| | | 90 min | ASW vs. 0.1% SDS | **** | <0,0001 |
| | | | ASW vs. 100mM $NH_4Cl$ | *** | 0,0001 |
| | | | ASW vs. 10mM Chloroquine | **** | <0,0001 |
| | | | ASW vs. 1mM $CuSO_4$ | **** | <0,0001 |
| | | 120 min | ASW vs. 0.1% SDS | **** | <0,0001 |
| | | | ASW vs. 100mM $NH_4Cl$ | **** | <0,0001 |
| | | | ASW vs. 10mM Chloroquine | **** | <0,0001 |
| | | | ASW vs. 1mM $CuSO_4$ | **** | <0,0001 |

table S12 Number of cells used to generate average traces in Figure 2 and Figure S2.

| Cell | Stimulus | Number of cells |
|---|---|---|
| PSN | ASW | 9 |
| | 1mM $NH_4Cl$ | 19 |
| | 10mM $NH_4Cl$ | 14 |
| | 100mM $NH_4Cl$ | 35 |
| | 10mM CsCl | 10 |
| | 200μM Eucalyptol | 16 |
| | 200μM Carvacrol | 15 |
| | 10μM Butanol | 19 |
| | 0.1% SDS | 29 |
| | 10mM Chloroquine | 9 |
| | 1mM Glutamate | 10 |
| | 1M Glycerol | 10 |
| | 1mM Asparagine | 12 |
| ACCs | ASW | 6 |
| | 1mM $NH_4Cl$ | 13 |
| | 10mM $NH_4Cl$ | 12 |
| | 100mM $NH_4Cl$ | 15 |
| | 0.1% SDS | 9 |
| | 1M Glycerol | 10 |
| PSN | $1^{st}$ poke | 14 |
| | $2^{nd}$ poke | 16 |
| | $3^{rd}$ poke | 15 |
| | $4^{th}$ poke | 12 |
| | $5^{th}$ poke | 14 |
| | $6^{th}$ poke | 10 |
| | $7^{th}$ poke | 14 |
| | $8^{th}$ poke | 13 |
| | $9^{th}$ poke | 14 |

| | | |
|---|---|---|
| ACC | 1st poke | 15 |
| | 2nd poke | 14 |
| | 3rd poke | 17 |
| | 4th poke | 13 |
| | 5th poke | 12 |
| | 6th poke | 14 |
| | 7th poke | 14 |
| | 8th poke | 14 |
| | 9th poke | 13 |
| PSN | 1st press | 12 |
| | 2nd press | 14 |
| | 3rd press | 14 |
| | 4th press | 14 |
| | 5th press | 13 |
| | 6th press | 11 |
| | 7th press | 11 |
| ACC | 1st press | 14 |
| | 2nd press | 13 |
| | 3rd press | 13 |
| | 4th press | 13 |
| | 5th press | 12 |
| | 6th press | 12 |
| | 7th press | 11 |
| ACC (GFP) | ASW | 5 |
| | 1mM $NH_4Cl$ | 7 |
| PSN (GFP) | ASW | 5 |
| | 1mM $NH_4Cl$ | 5 |

table S13 Statistics for summary quantifications presented in Figure 2P.

| Figure & Panel | Test | Comparison | P Value summary | P Value |
|---|---|---|---|---|
| 2P | Kruskal-Wallis | | | |
| | | Time | **** | <0,0001 |
| | Dunn's test | ASW vs 1mM $NH_4Cl$ | * | 0,0106 |
| | | ASW vs. 10 mM $NH_4Cl$ | *** | 0,0007 |
| | | ASW vs. 100 mM $NH_4Cl$ | **** | <0,0001 |
| | | ASW vs. 10mM CsCl | **** | <0,0001 |
| | | ASW vs 200μM Eucalyptol | ** | 0,0036 |
| | | ASW vs 200μM Carvacrol | *** | 0,0007 |
| | | ASW vs. 10μM Butanol | **** | <0,0001 |
| | | ASW vs. 0.1% SDS | **** | <0,0001 |
| | | ASW vs. 10mM Chloroquine | * | 0,0213 |
| | | ASW vs. 1mM Glutamate | ns | 0,0696 |
| | | ASW vs. 1M Glycerol | ns | 0,5309 |
| | | ASW vs. 1mM Asparagine | ns | 0,5737 |

table S14 Statistics for summary quantifications presented in Figure 2U.

| Figure & Panel | Test | Comparison | P Value summary | P Value |
|---|---|---|---|---|
| 2U | Kruskal-Wallis | | | |
| | | Time | ** | 0,0014 |
| | Dunn's test | ASW vs 1mM $NH_4Cl$ | * | 0,0159 |
| | | ASW vs. 10 mM $NH_4Cl$ | * | 0,0146 |
| | | ASW vs. 100 mM $NH_4Cl$ | ** | 0,0078 |
| | | ASW vs. 0.1% SDS | * | 0,0168 |
| | | ASW vs. 1M Glycerol | ns | 0,9103 |

table S15 Statistics for settlement rates analysis of *pc2>GFP* transgenic animals.

| Figure & Panel | Test | Data point | Comparison | P Value summary | P Value |
|---|---|---|---|---|---|
| Fig. S6A | Two-way RM ANOVA | | | | |
| | | | Time | **** | <0,0001 |
| | | | Fraction Settled | ns | 0,4497 |
| | | | Time x Fraction Settled | ** | 0,0032 |
| | Dunnett's multiple comparisons test | | | | |
| | | 30 min | pc2>GFP (-)CNO vs pc2>GFP (+) CNO | ns | >0,9999 |
| | | 60 min | pc2>GFP (-)CNO vs pc2>GFP (+) CNO | ns | >0,9999 |
| | | 90 min | pc2>GFP (-)CNO vs pc2>GFP (+) CNO | ns | >0,9999 |
| | | 120 min | pc2>GFP (-)CNO vs pc2>GFP (+) CNO | ns | >0,9999 |
| | | 150 min | pc2>GFP (-)CNO vs pc2>GFP (+) CNO | ns | >0,9999 |
| | | 180 min | pc2>GFP (-)CNO vs pc2>GFP (+) CNO | ns | >0,9999 |
| | | 210 min | pc2>GFP (-)CNO vs pc2>GFP (+) CNO | ns | >0,9999 |
| | | 240 min | pc2>GFP (-)CNO vs pc2>GFP (+) CNO | ns | >0,9999 |
| | | 270 min | pc2>GFP (-)CNO vs pc2>GFP (+) CNO | ns | >0,9999 |
| | | 300 min | pc2>GFP (-)CNO vs pc2>GFP (+) CNO | ns | >0,9999 |
| | | 330 min | pc2>GFP (-)CNO vs pc2>GFP (+) CNO | ns | 0,3348 |
| | | 360 min | pc2>GFP (-)CNO vs pc2>GFP (+) CNO | ns | 0,0676 |
| | | 390 min | pc2>GFP (-)CNO vs pc2>GFP (+) CNO | ns | 0,3348 |
| | | 420 min | pc2>GFP (-)CNO vs pc2>GFP (+) CNO | ns | 0,1818 |
| | | 450 min | pc2>GFP (-)CNO vs pc2>GFP (+) CNO | ns | 0,0676 |

| | | | | | |
|---|---|---|---|---|---|
| | | 480 min | pc2>GFP (-)CNO vs pc2>GFP (+) CNO | ns | >0,9999 |
| | | 510 min | pc2>GFP (-)CNO vs pc2>GFP (+) CNO | ns | >0,9999 |
| | | 540 min | pc2>GFP (-)CNO vs pc2>GFP (+) CNO | ns | >0,9999 |
| | | 570 min | pc2>GFP (-)CNO vs pc2>GFP (+) CNO | ns | >0,9999 |
| | | 600 min | pc2>GFP (-)CNO vs pc2>GFP (+) CNO | ns | >0,9999 |

table S16 Statistics for fraction of larvae that have completed tail regression for *pc2>GFP*

| Figure & Panel | Test | Comparison | P Value summary | P Value |
|---|---|---|---|---|
| Fig. S6B | Welch's test | pc2>GFP (-)CNO vs pc2>GFP (+) CNO | ns | 0,9815 |

table S17 Statistics for settlement rates analysis for *pc2>hM4D*, *βγ-crystallin>hM4D* single and *pc2>hM4D*; *βγ-crystallin>hM4D* double transgenics

| Figure & Panel | Test | Data point | Comparison | P Value summary | P Value |
|---|---|---|---|---|---|
| Fig. 5A | Two-way RM ANOVA | | | | |
| | | | Time | **** | <0,0001 |
| | | | Fraction Settled | **** | <0,0001 |
| | | | Time x Fraction Settled | **** | <0,0001 |
| | Tukey's multiple comparisons test | | | | |
| | | 30 min | *pc2>hM4D* CNO (-) vs. *pc2>hM4D* CNO (+) | ** | 0,0065 |
| | | | *pc2>hM4D* CNO (-) vs. *βγ-crystallin>hM4D* CNO (-) | ns | 0,8667 |
| | | | *pc2>hM4D* CNO (-) vs. *βγ-crystallin>hM4D* CNO (+) | ns | 0,2755 |
| | | | *pc2>hM4D* CNO (-) vs. *pc2>hM4D; βγ-crystallin>hM4D* CNO (-) | ns | 0,4578 |
| | | | *pc2>hM4D* CNO (-) vs. *pc2>hM4D; βγ-crystallin>hM4D* CNO (+) | ** | 0,0095 |
| | | | *pc2>hM4D* CNO (+) vs. *βγ-crystallin>hM4D* CNO (-) | ** | 0,0031 |
| | | | *pc2>hM4D* CNO (+) vs. *βγ-crystallin>hM4D* CNO (+) | ns | 0,0990 |
| | | | *pc2>hM4D* CNO (+) vs. *pc2>hM4D; βγ-crystallin>hM4D* CNO (-) | ns | 0,1364 |
| | | | *pc2>hM4D* CNO (+) vs. *pc2>hM4D; βγ-crystallin>hM4D* CNO (+) | ns | >0,9999 |
| | | | *βγ-crystallin>hM4D* CNO (-) vs. *βγ-crystallin>hM4D* CNO (+) | ns | 0,3305 |
| | | | *βγ-crystallin>hM4D* CNO (-) vs. *pc2>hM4D; βγ-crystallin>hM4D* CNO (-) | ns | 0,9205 |
| | | | *βγ-crystallin>hM4D* CNO CNO (-) vs. *pc2>hM4D; βγ-crystallin>hM4D* CNO (+) | ** | 0,0064 |
| | | | *βγ-crystallin>hM4D* CNO (+) vs. *pc2>hM4D; βγ-crystallin>hM4D* CNO (-) | ns | 0,9788 |

| | | | | | |
|---|---|---|---|---|---|
| | | | *βγ-crystallin>hM4D* CNO (+) vs. *pc2>hM4D; βγ-crystallin>hM4D* CNO (+) | * | 0,0313 |
| | | | *pc2>hM4D; βγ-crystallin>hM4D* CNO (-)vs. *pc2>hM4D; βγ-crystallin>hM4D* CNO (+) | ns | 0,1709 |
| | | 60 min | *pc2>hM4D* CNO (-) vs. *pc2>hM4D* CNO (+) | ** | 0,0069 |
| | | | *pc2>hM4D* CNO (-) vs. *βγ-crystallin>hM4D* CNO (-) | ns | >0,9999 |
| | | | *pc2>hM4D* CNO (-) vs. *βγ-crystallin>hM4D* CNO (+) | ns | 0,2720 |
| | | | *pc2>hM4D* CNO (-) vs. *pc2>hM4D; βγ-crystallin>hM4D* CNO (-) | ns | 0,9993 |
| | | | *pc2>hM4D* CNO (-) vs. *pc2>hM4D; βγ-crystallin>hM4D* CNO (+) | ** | 0,0030 |
| | | | *pc2>hM4D* CNO (+) vs. *βγ-crystallin>hM4D* CNO (-) | ** | 0,0018 |
| | | | *pc2>hM4D* CNO (+) vs. *βγ-crystallin>hM4D* CNO (+) | ns | 0,0608 |
| | | | *pc2>hM4D* CNO (+) vs. *pc2>hM4D; βγ-crystallin>hM4D* CNO (-) | ** | 0,0047 |
| | | | *pc2>hM4D* CNO (+) vs. *pc2>hM4D; βγ-crystallin>hM4D* CNO (+) | ns | 0,9320 |
| | | | *βγ-crystallin>hM4D* CNO (-) vs. *βγ-crystallin>hM4D* CNO (+) | ns | 0,4301 |
| | | | *βγ-crystallin>hM4D* CNO (-) vs. *pc2>hM4D; βγ-crystallin>hM4D* CNO (-) | ns | 0,9991 |
| | | | *βγ-crystallin>hM4D* CNO CNO (-) vs. *pc2>hM4D; βγ-crystallin>hM4D* CNO (+) | ** | 0,0030 |
| | | | *βγ-crystallin>hM4D* CNO (+) vs. *pc2>hM4D; βγ-crystallin>hM4D* CNO (-) | ns | 0,5930 |
| | | | *βγ-crystallin>hM4D* CNO (+) vs. *pc2>hM4D; βγ-crystallin>hM4D* CNO (+) | ns | 0,1153 |
| | | | *pc2>hM4D; βγ-crystallin>hM4D* CNO (-)vs. *pc2>hM4D; βγ-crystallin>hM4D* CNO (+) | * | 0,0149 |
| | | 90 min | *pc2>hM4D* CNO (-) vs. *pc2>hM4D* CNO (+) | * | 0,0420 |
| | | | *pc2>hM4D* CNO (-) vs. *βγ-crystallin>hM4D* CNO (-) | ns | 0,9980 |
| | | | *pc2>hM4D* CNO (-) vs. *βγ-crystallin>hM4D* CNO (+) | ns | 0,3146 |
| | | | *pc2>hM4D* CNO (-) vs. *pc2>hM4D; βγ-crystallin>hM4D* CNO (-) | ns | >0,9999 |
| | | | *pc2>hM4D* CNO (-) vs. *pc2>hM4D; βγ-crystallin>hM4D* CNO (+) | ** | 0,0021 |
| | | | *pc2>hM4D* CNO (+) vs. *βγ-crystallin>hM4D* CNO (-) | ns | 0,0858 |
| | | | *pc2>hM4D* CNO (+) vs. *βγ-crystallin>hM4D* CNO (+) | ns | 0,1560 |
| | | | *pc2>hM4D* CNO (+) vs. *pc2>hM4D; βγ-crystallin>hM4D* CNO (-) | * | 0,0459 |
| | | | *pc2>hM4D* CNO (+) vs. *pc2>hM4D; βγ-crystallin>hM4D* CNO (+) | ns | 0,9999 |
| | | | *βγ-crystallin>hM4D* CNO (-) vs. *βγ-crystallin>hM4D* CNO (+) | ns | 0,3701 |
| | | | *βγ-crystallin>hM4D* CNO (-) vs. *pc2>hM4D; βγ-crystallin>hM4D* CNO (-) | ns | >0,9999 |

| | | | | | |
|---|---|---|---|---|---|
| | | | *βγ-crystallin>hM4D* CNO CNO (-) vs. *pc2>hM4D; βγ-crystallin>hM4D* CNO (+) | * | 0,0151 |
| | | | *βγ-crystallin>hM4D* CNO (+) vs. *pc2>hM4D; βγ-crystallin>hM4D* CNO (-) | ns | 0,3216 |
| | | | *βγ-crystallin>hM4D* CNO (+) vs. *pc2>hM4D; βγ-crystallin>hM4D* CNO (+) | ns | 0,5132 |
| | | | *pc2>hM4D; βγ-crystallin>hM4D* CNO (-)vs. *pc2>hM4D; βγ-crystallin>hM4D* CNO (+) | * | 0,0191 |
| | | 120 min | *pc2>hM4D* CNO (-) vs. *pc2>hM4D* CNO (+) | * | 0,0447 |
| | | | *pc2>hM4D* CNO (-) vs. *βγ-crystallin>hM4D* CNO (-) | ns | 0,4688 |
| | | | *pc2>hM4D* CNO (-) vs. *βγ-crystallin>hM4D* CNO (+) | ns | 0,1757 |
| | | | *pc2>hM4D* CNO (-) vs. *pc2>hM4D; βγ-crystallin>hM4D* CNO (-) | ns | 0,9925 |
| | | | *pc2>hM4D* CNO (-) vs. *pc2>hM4D; βγ-crystallin>hM4D* CNO (+) | * | 0,0221 |
| | | | *pc2>hM4D* CNO (+) vs. *βγ-crystallin>hM4D* CNO (-) | * | 0,0122 |
| | | | *pc2>hM4D* CNO (+) vs. *βγ-crystallin>hM4D* CNO (+) | ns | >0,9999 |
| | | | *pc2>hM4D* CNO (+) vs. *pc2>hM4D; βγ-crystallin>hM4D* CNO (-) | ns | 0,0658 |
| | | | *pc2>hM4D* CNO (+) vs. *pc2>hM4D; βγ-crystallin>hM4D* CNO (+) | ns | 0,9601 |
| | | | *βγ-crystallin>hM4D* CNO (-) vs. *βγ-crystallin>hM4D* CNO (+) | * | 0,0246 |
| | | | *βγ-crystallin>hM4D* CNO (-) vs. *pc2>hM4D; βγ-crystallin>hM4D* CNO (-) | ns | 0,9299 |
| | | | *βγ-crystallin>hM4D* CNO CNO (-) vs. *pc2>hM4D; βγ-crystallin>hM4D* CNO (+) | ** | 0,0023 |
| | | | *βγ-crystallin>hM4D* CNO (+) vs. *pc2>hM4D; βγ-crystallin>hM4D* CNO (-) | * | 0,0467 |
| | | | *βγ-crystallin>hM4D* CNO (+) vs. *pc2>hM4D; βγ-crystallin>hM4D* CNO (+) | ns | 0,9826 |
| | | | *pc2>hM4D; βγ-crystallin>hM4D* CNO (-)vs. *pc2>hM4D; βγ-crystallin>hM4D* CNO (+) | * | 0,0196 |
| | | 150 min | *pc2>hM4D* CNO (-) vs. *pc2>hM4D* CNO (+) | * | 0,0471 |
| | | | *pc2>hM4D* CNO (-) vs. *βγ-crystallin>hM4D* CNO (-) | ns | 0,5583 |
| | | | *pc2>hM4D* CNO (-) vs. *βγ-crystallin>hM4D* CNO (+) | ns | 0,1450 |
| | | | *pc2>hM4D* CNO (-) vs. *pc2>hM4D; βγ-crystallin>hM4D* CNO (-) | ns | 0,9993 |
| | | | *pc2>hM4D* CNO (-) vs. *pc2>hM4D; βγ-crystallin>hM4D* CNO (+) | * | 0,0152 |
| | | | *pc2>hM4D* CNO (+) vs. *βγ-crystallin>hM4D* CNO (-) | ** | 0,0019 |
| | | | *pc2>hM4D* CNO (+) vs. *βγ-crystallin>hM4D* CNO (+) | ns | >0,9999 |
| | | | *pc2>hM4D* CNO (+) vs. *pc2>hM4D; βγ-crystallin>hM4D* CNO (-) | ns | 0,0743 |

| | | | | | |
|---|---|---|---|---|---|
| | | | *pc2>hM4D* CNO (+) vs. *pc2>hM4D; βγ-crystallin>hM4D* CNO (+) | ns | 0,9524 |
| | | | *βγ-crystallin>hM4D* CNO (-) vs. *βγ-crystallin>hM4D* CNO (+) | ** | 0,0057 |
| | | | *βγ-crystallin>hM4D* CNO (-) vs. *pc2>hM4D; βγ-crystallin>hM4D* CNO (-) | ns | 0,8222 |
| | | | *βγ-crystallin>hM4D* CNO CNO (-) vs. *pc2>hM4D; βγ-crystallin>hM4D* CNO (+) | *** | 0,0004 |
| | | | *βγ-crystallin>hM4D* CNO (+) vs. *pc2>hM4D; βγ-crystallin>hM4D* CNO (-) | * | 0,0475 |
| | | | *βγ-crystallin>hM4D* CNO (+) vs. *pc2>hM4D; βγ-crystallin>hM4D* CNO (+) | ns | 0,9189 |
| | | | *pc2>hM4D; βγ-crystallin>hM4D* CNO (-)vs. *pc2>hM4D; βγ-crystallin>hM4D* CNO (+) | * | 0,0124 |
| | | 180 min | *pc2>hM4D* CNO (-) vs. *pc2>hM4D* CNO (+) | * | 0,0143 |
| | | | *pc2>hM4D* CNO (-) vs. *βγ-crystallin>hM4D* CNO (-) | ns | 0,6095 |
| | | | *pc2>hM4D* CNO (-) vs. *βγ-crystallin>hM4D* CNO (+) | ns | 0,2004 |
| | | | *pc2>hM4D* CNO (-) vs. *pc2>hM4D; βγ-crystallin>hM4D* CNO (-) | ns | 0,9759 |
| | | | *pc2>hM4D* CNO (-) vs. *pc2>hM4D; βγ-crystallin>hM4D* CNO (+) | * | 0,0161 |
| | | | *pc2>hM4D* CNO (+) vs. *βγ-crystallin>hM4D* CNO (-) | *** | 0,0009 |
| | | | *pc2>hM4D* CNO (+) vs. *βγ-crystallin>hM4D* CNO (+) | ns | 0,9999 |
| | | | *pc2>hM4D* CNO (+) vs. *pc2>hM4D; βγ-crystallin>hM4D* CNO (-) | ** | 0,0082 |
| | | | *pc2>hM4D* CNO (+) vs. *pc2>hM4D; βγ-crystallin>hM4D* CNO (+) | ns | 0,9984 |
| | | | *βγ-crystallin>hM4D* CNO (-) vs. *βγ-crystallin>hM4D* CNO (+) | ** | 0,0092 |
| | | | *βγ-crystallin>hM4D* CNO (-) vs. *pc2>hM4D; βγ-crystallin>hM4D* CNO (-) | ns | 0,9731 |
| | | | *βγ-crystallin>hM4D* CNO CNO (-) vs. *pc2>hM4D; βγ-crystallin>hM4D* CNO (+) | ** | 0,0020 |
| | | | *βγ-crystallin>hM4D* CNO (+) vs. *pc2>hM4D; βγ-crystallin>hM4D* CNO (-) | ** | 0,0058 |
| | | | *βγ-crystallin>hM4D* CNO (+) vs. *pc2>hM4D; βγ-crystallin>hM4D* CNO (+) | ns | 0,9611 |
| | | | *pc2>hM4D; βγ-crystallin>hM4D* CNO (-)vs. *pc2>hM4D; βγ-crystallin>hM4D* CNO (+) | ** | 0,0024 |
| | | 210 min | *pc2>hM4D* CNO (-) vs. *pc2>hM4D* CNO (+) | *** | 0,0007 |
| | | | *pc2>hM4D* CNO (-) vs. *βγ-crystallin>hM4D* CNO (-) | ns | 0,8852 |
| | | | *pc2>hM4D* CNO (-) vs. *βγ-crystallin>hM4D* CNO (+) | * | 0,0363 |
| | | | *pc2>hM4D* CNO (-) vs. *pc2>hM4D; βγ-crystallin>hM4D* CNO (-) | ns | 0,9998 |
| | | | *pc2>hM4D* CNO (-) vs. *pc2>hM4D; βγ-crystallin>hM4D* CNO (+) | ** | 0,0013 |

| | | | | | |
|---|---|---|---|---|---|
| | | | *pc2>hM4D* CNO (+) vs. *βγ-crystallin>hM4D* CNO (-) | *** | 0,0004 |
| | | | *pc2>hM4D* CNO (+) vs. *βγ-crystallin>hM4D* CNO (+) | ns | 0,9555 |
| | | | *pc2>hM4D* CNO (+) vs. *pc2>hM4D; βγ-crystallin>hM4D* CNO (-) | ** | 0,0029 |
| | | | *pc2>hM4D* CNO (+) vs. *pc2>hM4D; βγ-crystallin>hM4D* CNO (+) | ns | 0,9902 |
| | | | *βγ-crystallin>hM4D* CNO (-) vs. *βγ-crystallin>hM4D* CNO (+) | ** | 0,0017 |
| | | | *βγ-crystallin>hM4D* CNO (-) vs. *pc2>hM4D; βγ-crystallin>hM4D* CNO (-) | ns | 0,9599 |
| | | | *βγ-crystallin>hM4D* CNO CNO (-) vs. *pc2>hM4D; βγ-crystallin>hM4D* CNO (+) | *** | 0,0002 |
| | | | *βγ-crystallin>hM4D* CNO (+) vs. *pc2>hM4D; βγ-crystallin>hM4D* CNO (-) | ** | 0,0015 |
| | | | *βγ-crystallin>hM4D* CNO (+) vs. *pc2>hM4D; βγ-crystallin>hM4D* CNO (+) | ns | 0,7523 |
| | | | *pc2>hM4D; βγ-crystallin>hM4D* CNO (-)vs. *pc2>hM4D; βγ-crystallin>hM4D* CNO (+) | ** | 0,0024 |
| | | 240 min | *pc2>hM4D* CNO (-) vs. *pc2>hM4D* CNO (+) | ** | 0,0046 |
| | | | *pc2>hM4D* CNO (-) vs. *βγ-crystallin>hM4D* CNO (-) | ns | 0,9980 |
| | | | *pc2>hM4D* CNO (-) vs. *βγ-crystallin>hM4D* CNO (+) | * | 0,0436 |
| | | | *pc2>hM4D* CNO (-) vs. *pc2>hM4D; βγ-crystallin>hM4D* CNO (-) | ns | 0,8681 |
| | | | *pc2>hM4D* CNO (-) vs. *pc2>hM4D; βγ-crystallin>hM4D* CNO (+) | *** | 0,0009 |
| | | | *pc2>hM4D* CNO (+) vs. *βγ-crystallin>hM4D* CNO (-) | ** | 0,0078 |
| | | | *pc2>hM4D* CNO (+) vs. *βγ-crystallin>hM4D* CNO (+) | ns | 0,9747 |
| | | | *pc2>hM4D* CNO (+) vs. *pc2>hM4D; βγ-crystallin>hM4D* CNO (-) | ** | 0,0051 |
| | | | *pc2>hM4D* CNO (+) vs. *pc2>hM4D; βγ-crystallin>hM4D* CNO (+) | ns | 0,9771 |
| | | | *βγ-crystallin>hM4D* CNO (-) vs. *βγ-crystallin>hM4D* CNO (+) | ** | 0,0035 |
| | | | *βγ-crystallin>hM4D* CNO (-) vs. *pc2>hM4D; βγ-crystallin>hM4D* CNO (-) | ns | 0,5963 |
| | | | *βγ-crystallin>hM4D* CNO CNO (-) vs. *pc2>hM4D; βγ-crystallin>hM4D* CNO (+) | *** | 0,0004 |
| | | | *βγ-crystallin>hM4D* CNO (+) vs. *pc2>hM4D; βγ-crystallin>hM4D* CNO (-) | *** | 0,0002 |
| | | | *βγ-crystallin>hM4D* CNO (+) vs. *pc2>hM4D; βγ-crystallin>hM4D* CNO (+) | ns | 0,6560 |
| | | | *pc2>hM4D; βγ-crystallin>hM4D* CNO (-)vs. *pc2>hM4D; βγ-crystallin>hM4D* CNO (+) | *** | 0,0004 |
| | | 270 min | *pc2>hM4D* CNO (-) vs. *pc2>hM4D* CNO (+) | * | 0,0443 |
| | | | *pc2>hM4D* CNO (-) vs. *βγ-crystallin>hM4D* CNO (-) | ns | 0,9725 |

| | | | | | |
|---|---|---|---|---|---|
| | | | *pc2>hM4D* CNO (-) vs. *βγ-crystallin>hM4D* CNO (+) | * | 0,0414 |
| | | | *pc2>hM4D* CNO (-) vs. *pc2>hM4D; βγ-crystallin>hM4D* CNO (-) | ns | 0,9851 |
| | | | *pc2>hM4D* CNO (-) vs. *pc2>hM4D; βγ-crystallin>hM4D* CNO (+) | ** | 0,0013 |
| | | | *pc2>hM4D* CNO (+) vs. *βγ-crystallin>hM4D* CNO (-) | ns | 0,0513 |
| | | | *pc2>hM4D* CNO (+) vs. *βγ-crystallin>hM4D* CNO (+) | ns | 0,9818 |
| | | | *pc2>hM4D* CNO (+) vs. *pc2>hM4D; βγ-crystallin>hM4D* CNO (-) | * | 0,0338 |
| | | | *pc2>hM4D* CNO (+) vs. *pc2>hM4D; βγ-crystallin>hM4D* CNO (+) | ns | 0,9738 |
| | | | *βγ-crystallin>hM4D* CNO (-) vs. *βγ-crystallin>hM4D* CNO (+) | ns | 0,0569 |
| | | | *βγ-crystallin>hM4D* CNO (-) vs. *pc2>hM4D; βγ-crystallin>hM4D* CNO (-) | ns | 0,4936 |
| | | | *βγ-crystallin>hM4D* CNO CNO (-) vs. *pc2>hM4D; βγ-crystallin>hM4D* CNO (+) | ** | 0,0016 |
| | | | *βγ-crystallin>hM4D* CNO (+) vs. *pc2>hM4D; βγ-crystallin>hM4D* CNO (-) | ** | 0,0040 |
| | | | *βγ-crystallin>hM4D* CNO (+) vs. *pc2>hM4D; βγ-crystallin>hM4D* CNO (+) | ns | 0,7255 |
| | | | *pc2>hM4D; βγ-crystallin>hM4D* CNO (-)vs. *pc2>hM4D; βγ-crystallin>hM4D* CNO (+) | ** | 0,0013 |
| | | 300 min | *pc2>hM4D* CNO (-) vs. *pc2>hM4D* CNO (+) | * | 0,0380 |
| | | | *pc2>hM4D* CNO (-) vs. *βγ-crystallin>hM4D* CNO (-) | ns | 0,5530 |
| | | | *pc2>hM4D* CNO (-) vs. *βγ-crystallin>hM4D* CNO (+) | * | 0,0301 |
| | | | *pc2>hM4D* CNO (-) vs. *pc2>hM4D; βγ-crystallin>hM4D* CNO (-) | ns | 0,9996 |
| | | | *pc2>hM4D* CNO (-) vs. *pc2>hM4D; βγ-crystallin>hM4D* CNO (+) | ** | 0,0020 |
| | | | *pc2>hM4D* CNO (+) vs. *βγ-crystallin>hM4D* CNO (-) | * | 0,0498 |
| | | | *pc2>hM4D* CNO (+) vs. *βγ-crystallin>hM4D* CNO (+) | ns | 0,8225 |
| | | | *pc2>hM4D* CNO (+) vs. *pc2>hM4D; βγ-crystallin>hM4D* CNO (-) | * | 0,0313 |
| | | | *pc2>hM4D* CNO (+) vs. *pc2>hM4D; βγ-crystallin>hM4D* CNO (+) | ns | 0,8398 |
| | | | *βγ-crystallin>hM4D* CNO (-) vs. *βγ-crystallin>hM4D* CNO (+) | ns | 0,1440 |
| | | | *βγ-crystallin>hM4D* CNO (-) vs. *pc2>hM4D; βγ-crystallin>hM4D* CNO (-) | ns | 0,7506 |
| | | | *βγ-crystallin>hM4D* CNO CNO (-) vs. *pc2>hM4D; βγ-crystallin>hM4D* CNO (+) | ** | 0,0023 |
| | | | *βγ-crystallin>hM4D* CNO (+) vs. *pc2>hM4D; βγ-crystallin>hM4D* CNO (-) | * | 0,0205 |
| | | | *βγ-crystallin>hM4D* CNO (+) vs. *pc2>hM4D; βγ-crystallin>hM4D* CNO (+) | ns | 0,2048 |
| | | | *pc2>hM4D; βγ-crystallin>hM4D* CNO (-)vs. *pc2>hM4D; βγ-crystallin>hM4D* CNO (+) | ** | 0,0030 |

| | | | | | |
|---|---|---|---|---|---|
| | | 330 min | *pc2>hM4D* CNO (-) vs. *pc2>hM4D* CNO (+) | ns | 0,0664 |
| | | | *pc2>hM4D* CNO (-) vs. *βγ-crystallin>hM4D* CNO (-) | ns | 0,9403 |
| | | | *pc2>hM4D* CNO (-) vs. *βγ-crystallin>hM4D* CNO (+) | * | 0,0216 |
| | | | *pc2>hM4D* CNO (-) vs. *pc2>hM4D; βγ-crystallin>hM4D* CNO (-) | ns | 0,9933 |
| | | | *pc2>hM4D* CNO (-) vs. *pc2>hM4D; βγ-crystallin>hM4D* CNO (+) | ** | 0,0018 |
| | | | *pc2>hM4D* CNO (+) vs. *βγ-crystallin>hM4D* CNO (-) | * | 0,0270 |
| | | | *pc2>hM4D* CNO (+) vs. *βγ-crystallin>hM4D* CNO (+) | ns | 0,7372 |
| | | | *pc2>hM4D* CNO (+) vs. *pc2>hM4D; βγ-crystallin>hM4D* CNO (-) | * | 0,0335 |
| | | | *pc2>hM4D* CNO (+) vs. *pc2>hM4D; βγ-crystallin>hM4D* CNO (+) | ns | 0,6751 |
| | | | *βγ-crystallin>hM4D* CNO (-) vs. *βγ-crystallin>hM4D* CNO (+) | * | 0,0350 |
| | | | *βγ-crystallin>hM4D* CNO (-) vs. *pc2>hM4D; βγ-crystallin>hM4D* CNO (-) | ns | >0,9999 |
| | | | *βγ-crystallin>hM4D* CNO CNO (-) vs. *pc2>hM4D; βγ-crystallin>hM4D* CNO (+) | ** | 0,0016 |
| | | | *βγ-crystallin>hM4D* CNO (+) vs. *pc2>hM4D; βγ-crystallin>hM4D* CNO (-) | ** | 0,0090 |
| | | | *βγ-crystallin>hM4D* CNO (+) vs. *pc2>hM4D; βγ-crystallin>hM4D* CNO (+) | ns | 0,0875 |
| | | | *pc2>hM4D; βγ-crystallin>hM4D* CNO (-)vs. *pc2>hM4D; βγ-crystallin>hM4D* CNO (+) | ** | 0,0050 |
| | | 360 min | *pc2>hM4D* CNO (-) vs. *pc2>hM4D* CNO (+) | ns | 0,0549 |
| | | | *pc2>hM4D* CNO (-) vs. *βγ-crystallin>hM4D* CNO (-) | ns | 0,9605 |
| | | | *pc2>hM4D* CNO (-) vs. *βγ-crystallin>hM4D* CNO (+) | ns | 0,0796 |
| | | | *pc2>hM4D* CNO (-) vs. *pc2>hM4D; βγ-crystallin>hM4D* CNO (-) | ns | 0,9797 |
| | | | *pc2>hM4D* CNO (-) vs. *pc2>hM4D; βγ-crystallin>hM4D* CNO (+) | ** | 0,0045 |
| | | | *pc2>hM4D* CNO (+) vs. *βγ-crystallin>hM4D* CNO (-) | ** | 0,0057 |
| | | | *pc2>hM4D* CNO (+) vs. *βγ-crystallin>hM4D* CNO (+) | ns | 0,4735 |
| | | | *pc2>hM4D* CNO (+) vs. *pc2>hM4D; βγ-crystallin>hM4D* CNO (-) | * | 0,0275 |
| | | | *pc2>hM4D* CNO (+) vs. *pc2>hM4D; βγ-crystallin>hM4D* CNO (+) | ns | 0,8158 |
| | | | *βγ-crystallin>hM4D* CNO (-) vs. *βγ-crystallin>hM4D* CNO (+) | ns | 0,0990 |
| | | | *βγ-crystallin>hM4D* CNO (-) vs. *pc2>hM4D; βγ-crystallin>hM4D* CNO (-) | ns | >0,9999 |
| | | | *βγ-crystallin>hM4D* CNO CNO (-) vs. *pc2>hM4D; βγ-crystallin>hM4D* CNO (+) | ** | 0,0046 |
| | | | *βγ-crystallin>hM4D* CNO (+) vs. *pc2>hM4D; βγ-crystallin>hM4D* CNO (-) | * | 0,0184 |

| | | | | | |
|---|---|---|---|---|---|
| | | | *βγ-crystallin>hM4D* CNO (+) vs. *pc2>hM4D; βγ-crystallin>hM4D* CNO (+) | ns | 0,1032 |
| | | | *pc2>hM4D; βγ-crystallin>hM4D* CNO (-)vs. *pc2>hM4D; βγ-crystallin>hM4D* CNO (+) | * | 0,0157 |
| | | 390 min | *pc2>hM4D* CNO (-) vs. *pc2>hM4D* CNO (+) | * | 0,0193 |
| | | | *pc2>hM4D* CNO (-) vs. *βγ-crystallin>hM4D* CNO (-) | ns | 0,9601 |
| | | | *pc2>hM4D* CNO (-) vs. *βγ-crystallin>hM4D* CNO (+) | * | 0,0147 |
| | | | *pc2>hM4D* CNO (-) vs. *pc2>hM4D; βγ-crystallin>hM4D* CNO (-) | ns | 0,8472 |
| | | | *pc2>hM4D* CNO (-) vs. *pc2>hM4D; βγ-crystallin>hM4D* CNO (+) | *** | 0,0004 |
| | | | *pc2>hM4D* CNO (+) vs. *βγ-crystallin>hM4D* CNO (-) | ** | 0,0015 |
| | | | *pc2>hM4D* CNO (+) vs. *βγ-crystallin>hM4D* CNO (+) | ns | 0,5005 |
| | | | *pc2>hM4D* CNO (+) vs. *pc2>hM4D; βγ-crystallin>hM4D* CNO (-) | ** | 0,0093 |
| | | | *pc2>hM4D* CNO (+) vs. *pc2>hM4D; βγ-crystallin>hM4D* CNO (+) | ns | 0,6794 |
| | | | *βγ-crystallin>hM4D* CNO (-) vs. *βγ-crystallin>hM4D* CNO (+) | ** | 0,0079 |
| | | | *βγ-crystallin>hM4D* CNO (-) vs. *pc2>hM4D; βγ-crystallin>hM4D* CNO (-) | ns | 0,9340 |
| | | | *βγ-crystallin>hM4D* CNO CNO (-) vs. *pc2>hM4D; βγ-crystallin>hM4D* CNO (+) | *** | 0,0008 |
| | | | *βγ-crystallin>hM4D* CNO (+) vs. *pc2>hM4D; βγ-crystallin>hM4D* CNO (-) | ** | 0,0041 |
| | | | *βγ-crystallin>hM4D* CNO (+) vs. *pc2>hM4D; βγ-crystallin>hM4D* CNO (+) | * | 0,0347 |
| | | | *pc2>hM4D; βγ-crystallin>hM4D* CNO (-)vs. *pc2>hM4D; βγ-crystallin>hM4D* CNO (+) | ** | 0,0030 |
| | | 420 min | *pc2>hM4D* CNO (-) vs. *pc2>hM4D* CNO (+) | ** | 0,0084 |
| | | | *pc2>hM4D* CNO (-) vs. *βγ-crystallin>hM4D* CNO (-) | ns | 0,9705 |
| | | | *pc2>hM4D* CNO (-) vs. *βγ-crystallin>hM4D* CNO (+) | ** | 0,0033 |
| | | | *pc2>hM4D* CNO (-) vs. *pc2>hM4D; βγ-crystallin>hM4D* CNO (-) | ns | 0,6319 |
| | | | *pc2>hM4D* CNO (-) vs. *pc2>hM4D; βγ-crystallin>hM4D* CNO (+) | *** | 0,0004 |
| | | | *pc2>hM4D* CNO (+) vs. *βγ-crystallin>hM4D* CNO (-) | *** | 0,0007 |
| | | | *pc2>hM4D* CNO (+) vs. *βγ-crystallin>hM4D* CNO (+) | ns | 0,6157 |
| | | | *pc2>hM4D* CNO (+) vs. *pc2>hM4D; βγ-crystallin>hM4D* CNO (-) | ** | 0,0026 |
| | | | *pc2>hM4D* CNO (+) vs. *pc2>hM4D; βγ-crystallin>hM4D* CNO (+) | ns | 0,6073 |
| | | | *βγ-crystallin>hM4D* CNO (-) vs. *βγ-crystallin>hM4D* CNO (+) | ** | 0,0013 |
| | | | *βγ-crystallin>hM4D* CNO (-) vs. *pc2>hM4D; βγ-crystallin>hM4D* CNO (-) | ns | 0,7568 |

| | | | | | |
|---|---|---|---|---|---|
| | | | *βγ-crystallin>hM4D* CNO CNO (-) vs. *pc2>hM4D; βγ-crystallin>hM4D* CNO (+) | *** | 0,0007 |
| | | | *βγ-crystallin>hM4D* CNO (+) vs. *pc2>hM4D; βγ-crystallin>hM4D* CNO (-) | *** | 0,0002 |
| | | | *βγ-crystallin>hM4D* CNO (+) vs. *pc2>hM4D; βγ-crystallin>hM4D* CNO (+) | * | 0,0307 |
| | | | *pc2>hM4D; βγ-crystallin>hM4D* CNO (-)vs. *pc2>hM4D; βγ-crystallin>hM4D* CNO (+) | ** | 0,0013 |
| | | 450 min | *pc2>hM4D* CNO (-) vs. *pc2>hM4D* CNO (+) | ** | 0,0052 |
| | | | *pc2>hM4D* CNO (-) vs. *βγ-crystallin>hM4D* CNO (-) | ns | 0,9987 |
| | | | *pc2>hM4D* CNO (-) vs. *βγ-crystallin>hM4D* CNO (+) | ** | 0,0018 |
| | | | *pc2>hM4D* CNO (-) vs. *pc2>hM4D; βγ-crystallin>hM4D* CNO (-) | ns | 0,6013 |
| | | | *pc2>hM4D* CNO (-) vs. *pc2>hM4D; βγ-crystallin>hM4D* CNO (+) | *** | 0,0005 |
| | | | *pc2>hM4D* CNO (+) vs. *βγ-crystallin>hM4D* CNO (-) | ** | 0,0010 |
| | | | *pc2>hM4D* CNO (+) vs. *βγ-crystallin>hM4D* CNO (+) | ns | 0,8413 |
| | | | *pc2>hM4D* CNO (+) vs. *pc2>hM4D; βγ-crystallin>hM4D* CNO (-) | ** | 0,0026 |
| | | | *pc2>hM4D* CNO (+) vs. *pc2>hM4D; βγ-crystallin>hM4D* CNO (+) | ns | 0,5189 |
| | | | *βγ-crystallin>hM4D* CNO (-) vs. *βγ-crystallin>hM4D* CNO (+) | *** | 0,0009 |
| | | | *βγ-crystallin>hM4D* CNO (-) vs. *pc2>hM4D; βγ-crystallin>hM4D* CNO (-) | ns | 0,5446 |
| | | | *βγ-crystallin>hM4D* CNO CNO (-) vs. *pc2>hM4D; βγ-crystallin>hM4D* CNO (+) | *** | 0,0004 |
| | | | *βγ-crystallin>hM4D* CNO (+) vs. *pc2>hM4D; βγ-crystallin>hM4D* CNO (-) | *** | 0,0005 |
| | | | *βγ-crystallin>hM4D* CNO (+) vs. *pc2>hM4D; βγ-crystallin>hM4D* CNO (+) | * | 0,0418 |
| | | | *pc2>hM4D; βγ-crystallin>hM4D* CNO (-)vs. *pc2>hM4D; βγ-crystallin>hM4D* CNO (+) | *** | 0,0006 |
| | | 480 min | *pc2>hM4D* CNO (-) vs. *pc2>hM4D* CNO (+) | ** | 0,0022 |
| | | | *pc2>hM4D* CNO (-) vs. *βγ-crystallin>hM4D* CNO (-) | ns | 0,9999 |
| | | | *pc2>hM4D* CNO (-) vs. *βγ-crystallin>hM4D* CNO (+) | *** | 0,0008 |
| | | | *pc2>hM4D* CNO (-) vs. *pc2>hM4D; βγ-crystallin>hM4D* CNO (-) | ns | 0,6765 |
| | | | *pc2>hM4D* CNO (-) vs. *pc2>hM4D; βγ-crystallin>hM4D* CNO (+) | *** | 0,0004 |
| | | | *pc2>hM4D* CNO (+) vs. *βγ-crystallin>hM4D* CNO (-) | *** | 0,0007 |
| | | | *pc2>hM4D* CNO (+) vs. *βγ-crystallin>hM4D* CNO (+) | ns | 0,8932 |
| | | | *pc2>hM4D* CNO (+) vs. *pc2>hM4D; βγ-crystallin>hM4D* CNO (-) | ** | 0,0015 |

| | | | | | |
|---|---|---|---|---|---|
| | | | *pc2>hM4D* CNO (+) vs. *pc2>hM4D; βγ-crystallin>hM4D* CNO (+) | ns | 0,5938 |
| | | | *βγ-crystallin>hM4D* CNO (-) vs. *βγ-crystallin>hM4D* CNO (+) | *** | 0,0008 |
| | | | *βγ-crystallin>hM4D* CNO (-) vs. *pc2>hM4D; βγ-crystallin>hM4D* CNO (-) | ns | 0,5186 |
| | | | *βγ-crystallin>hM4D* CNO CNO (-) vs. *pc2>hM4D; βγ-crystallin>hM4D* CNO (+) | *** | 0,0004 |
| | | | *βγ-crystallin>hM4D* CNO (+) vs. *pc2>hM4D; βγ-crystallin>hM4D* CNO (-) | *** | 0,0003 |
| | | | *βγ-crystallin>hM4D* CNO (+) vs. *pc2>hM4D; βγ-crystallin>hM4D* CNO (+) | ns | 0,0671 |
| | | | *pc2>hM4D; βγ-crystallin>hM4D* CNO (-)vs. *pc2>hM4D; βγ-crystallin>hM4D* CNO (+) | *** | 0,0005 |
| | | 510 min | *pc2>hM4D* CNO (-) vs. *pc2>hM4D* CNO (+) | ** | 0,0015 |
| | | | *pc2>hM4D* CNO (-) vs. *βγ-crystallin>hM4D* CNO (-) | ns | >0,9999 |
| | | | *pc2>hM4D* CNO (-) vs. *βγ-crystallin>hM4D* CNO (+) | *** | 0,0004 |
| | | | *pc2>hM4D* CNO (-) vs. *pc2>hM4D; βγ-crystallin>hM4D* CNO (-) | ns | 0,5627 |
| | | | *pc2>hM4D* CNO (-) vs. *pc2>hM4D; βγ-crystallin>hM4D* CNO (+) | *** | 0,0004 |
| | | | *pc2>hM4D* CNO (+) vs. *βγ-crystallin>hM4D* CNO (-) | *** | 0,0008 |
| | | | *pc2>hM4D* CNO (+) vs. *βγ-crystallin>hM4D* CNO (+) | ns | 0,9137 |
| | | | *pc2>hM4D* CNO (+) vs. *pc2>hM4D; βγ-crystallin>hM4D* CNO (-) | ** | 0,0010 |
| | | | *pc2>hM4D* CNO (+) vs. *pc2>hM4D; βγ-crystallin>hM4D* CNO (+) | ns | 0,5938 |
| | | | *βγ-crystallin>hM4D* CNO (-) vs. *βγ-crystallin>hM4D* CNO (+) | *** | 0,0010 |
| | | | *βγ-crystallin>hM4D* CNO (-) vs. *pc2>hM4D; βγ-crystallin>hM4D* CNO (-) | ns | 0,4443 |
| | | | *βγ-crystallin>hM4D* CNO CNO (-) vs. *pc2>hM4D; βγ-crystallin>hM4D* CNO (+) | *** | 0,0005 |
| | | | *βγ-crystallin>hM4D* CNO (+) vs. *pc2>hM4D; βγ-crystallin>hM4D* CNO (-) | *** | 0,0005 |
| | | | *βγ-crystallin>hM4D* CNO (+) vs. *pc2>hM4D; βγ-crystallin>hM4D* CNO (+) | ns | 0,0804 |
| | | | *pc2>hM4D; βγ-crystallin>hM4D* CNO (-)vs. *pc2>hM4D; βγ-crystallin>hM4D* CNO (+) | *** | 0,0005 |
| | | 540 min | *pc2>hM4D* CNO (-) vs. *pc2>hM4D* CNO (+) | ** | 0,0016 |
| | | | *pc2>hM4D* CNO (-) vs. *βγ-crystallin>hM4D* CNO (-) | ns | >0,9999 |
| | | | *pc2>hM4D* CNO (-) vs. *βγ-crystallin>hM4D* CNO (+) | *** | 0,0006 |
| | | | *pc2>hM4D* CNO (-) vs. *pc2>hM4D; βγ-crystallin>hM4D* CNO (-) | ns | 0,7205 |
| | | | *pc2>hM4D* CNO (-) vs. *pc2>hM4D; βγ-crystallin>hM4D* CNO (+) | *** | 0,0006 |

| | | | | | |
|---|---|---|---|---|---|
| | | | *pc2>hM4D* CNO (+) vs. *βγ-crystallin>hM4D* CNO (-) | ** | 0,0016 |
| | | | *pc2>hM4D* CNO (+) vs. *βγ-crystallin>hM4D* CNO (+) | ns | 0,7227 |
| | | | *pc2>hM4D* CNO (+) vs. *pc2>hM4D; βγ-crystallin>hM4D* CNO (-) | *** | 0,0006 |
| | | | *pc2>hM4D* CNO (+) vs. *pc2>hM4D; βγ-crystallin>hM4D* CNO (+) | ns | 0,6444 |
| | | | *βγ-crystallin>hM4D* CNO (-) vs. *βγ-crystallin>hM4D* CNO (+) | *** | 0,0008 |
| | | | *βγ-crystallin>hM4D* CNO (-) vs. *pc2>hM4D; βγ-crystallin>hM4D* CNO (-) | ns | 0,3991 |
| | | | *βγ-crystallin>hM4D* CNO CNO (-) vs. *pc2>hM4D; βγ-crystallin>hM4D* CNO (+) | *** | 0,0003 |
| | | | *βγ-crystallin>hM4D* CNO (+) vs. *pc2>hM4D; βγ-crystallin>hM4D* CNO (-) | *** | 0,0001 |
| | | | *βγ-crystallin>hM4D* CNO (+) vs. *pc2>hM4D; βγ-crystallin>hM4D* CNO (+) | ns | 0,0842 |
| | | | *pc2>hM4D; βγ-crystallin>hM4D* CNO (-)vs. *pc2>hM4D; βγ-crystallin>hM4D* CNO (+) | *** | 0,0005 |
| | | 570 min | *pc2>hM4D* CNO (-) vs. *pc2>hM4D* CNO (+) | ** | 0,0014 |
| | | | *pc2>hM4D* CNO (-) vs. *βγ-crystallin>hM4D* CNO (-) | ns | 0,9990 |
| | | | *pc2>hM4D* CNO (-) vs. *βγ-crystallin>hM4D* CNO (+) | *** | 0,0005 |
| | | | *pc2>hM4D* CNO (-) vs. *pc2>hM4D; βγ-crystallin>hM4D* CNO (-) | ns | 0,9941 |
| | | | *pc2>hM4D* CNO (-) vs. *pc2>hM4D; βγ-crystallin>hM4D* CNO (+) | *** | 0,0005 |
| | | | *pc2>hM4D* CNO (+) vs. *βγ-crystallin>hM4D* CNO (-) | ** | 0,0012 |
| | | | *pc2>hM4D* CNO (+) vs. *βγ-crystallin>hM4D* CNO (+) | ns | 0,6963 |
| | | | *pc2>hM4D* CNO (+) vs. *pc2>hM4D; βγ-crystallin>hM4D* CNO (-) | *** | 0,0006 |
| | | | *pc2>hM4D* CNO (+) vs. *pc2>hM4D; βγ-crystallin>hM4D* CNO (+) | ns | 0,6095 |
| | | | *βγ-crystallin>hM4D* CNO (-) vs. *βγ-crystallin>hM4D* CNO (+) | *** | 0,0002 |
| | | | *βγ-crystallin>hM4D* CNO (-) vs. *pc2>hM4D; βγ-crystallin>hM4D* CNO (-) | ns | 0,5552 |
| | | | *βγ-crystallin>hM4D* CNO CNO (-) vs. *pc2>hM4D; βγ-crystallin>hM4D* CNO (+) | *** | 0,0002 |
| | | | *βγ-crystallin>hM4D* CNO (+) vs. *pc2>hM4D; βγ-crystallin>hM4D* CNO (-) | **** | <0,0001 |
| | | | *βγ-crystallin>hM4D* CNO (+) vs. *pc2>hM4D; βγ-crystallin>hM4D* CNO (+) | ns | 0,1056 |
| | | | *pc2>hM4D; βγ-crystallin>hM4D* CNO (-)vs. *pc2>hM4D; βγ-crystallin>hM4D* CNO (+) | *** | 0,0003 |
| | | 600 min | *pc2>hM4D* CNO (-) vs. *pc2>hM4D* CNO (+) | ** | 0,0014 |
| | | | *pc2>hM4D* CNO (-) vs. *βγ-crystallin>hM4D* CNO (-) | ns | >0,9999 |

| | | | | | |
|---|---|---|---|---|---|
| | | | *pc2>hM4D* CNO (-) vs. *βγ-crystallin>hM4D* CNO (+) | ** | 0,0016 |
| | | | *pc2>hM4D* CNO (-) vs. *pc2>hM4D; βγ-crystallin>hM4D* CNO (-) | ns | 0,9989 |
| | | | *pc2>hM4D* CNO (-) vs. *pc2>hM4D; βγ-crystallin>hM4D* CNO (+) | *** | 0,0005 |
| | | | *pc2>hM4D* CNO (+) vs. *βγ-crystallin>hM4D* CNO (-) | ** | 0,0016 |
| | | | *pc2>hM4D* CNO (+) vs. *βγ-crystallin>hM4D* CNO (+) | ns | 0,5060 |
| | | | *pc2>hM4D* CNO (+) vs. *pc2>hM4D; βγ-crystallin>hM4D* CNO (-) | ** | 0,0013 |
| | | | *pc2>hM4D* CNO (+) vs. *pc2>hM4D; βγ-crystallin>hM4D* CNO (+) | ns | 0,6964 |
| | | | *βγ-crystallin>hM4D* CNO (-) vs. *βγ-crystallin>hM4D* CNO (+) | ** | 0,0022 |
| | | | *βγ-crystallin>hM4D* CNO (-) vs. *pc2>hM4D; βγ-crystallin>hM4D* CNO (-) | ns | 0,9068 |
| | | | *βγ-crystallin>hM4D* CNO CNO (-) vs. *pc2>hM4D; βγ-crystallin>hM4D* CNO (+) | *** | 0,0002 |
| | | | *βγ-crystallin>hM4D* CNO (+) vs. *pc2>hM4D; βγ-crystallin>hM4D* CNO (-) | ** | 0,0017 |
| | | | *βγ-crystallin>hM4D* CNO (+) vs. *pc2>hM4D; βγ-crystallin>hM4D* CNO (+) | ns | 0,1573 |
| | | | *pc2>hM4D; βγ-crystallin>hM4D* CNO (-)vs. *pc2>hM4D; βγ-crystallin>hM4D* CNO (+) | *** | 0,0002 |

table S18 Statistics for settlement rates analysis for *pc2>hM4D*, *βγ-crystallin>hM4D* single and *pc2>hM4D*; *βγ-crystallin>hM4D* double transgenics in the presence of 10mM $NH_4Cl$

| Figure & Panel | Test | Data point | Comparison | P Value summary | P Value |
|---|---|---|---|---|---|
| Fig. 5B | Two-way RM ANOVA | | | | |
| | | | Time | **** | <0,0001 |
| | | | Fraction Settled | **** | <0,0001 |
| | | | Time x Fraction Settled | **** | <0,0001 |
| | Tukey's multiple comparisons test | | | | |
| | | 30 min | *pc2>hM4D* CNO (-) vs. *pc2>hM4D* CNO (+) | *** | 0,0003 |
| | | | *pc2>hM4D* CNO (-) vs. *βγ-crystallin>hM4D* CNO (-) | ns | 0,2751 |
| | | | *pc2>hM4D* CNO (-) vs. *βγ-crystallin>hM4D* CNO (+) | ** | 0,0012 |
| | | | *pc2>hM4D* CNO (-) vs. *pc2>hM4D; βγ-crystallin>hM4D* CNO (-) | ns | 0,9093 |
| | | | *pc2>hM4D* CNO (-) vs. *pc2>hM4D; βγ-crystallin>hM4D* CNO (+) | **** | <0,0001 |
| | | | *pc2>hM4D* CNO (+) vs. *βγ-crystallin>hM4D* CNO (-) | ** | 0,0012 |
| | | | *pc2>hM4D* CNO (+) vs. *βγ-crystallin>hM4D* CNO (+) | ns | 0,5755 |
| | | | *pc2>hM4D* CNO (+) vs. *pc2>hM4D; βγ-crystallin>hM4D* CNO (-) | ** | 0,0012 |

| | | | | | |
|---|---|---|---|---|---|
| | | | *pc2>hM4D* CNO (+) vs. *pc2>hM4D; βγ-crystallin>hM4D* CNO (+) | ns | 0,1151 |
| | | | *βγ-crystallin>hM4D* CNO (-) vs. *βγ-crystallin>hM4D* CNO (+) | ** | 0,0058 |
| | | | *βγ-crystallin>hM4D* CNO (-) vs. *pc2>hM4D; βγ-crystallin>hM4D* CNO (-) | ns | >0,9999 |
| | | | *βγ-crystallin>hM4D* CNO CNO (-) vs. *pc2>hM4D; βγ-crystallin>hM4D* CNO (+) | *** | 0,0002 |
| | | | *βγ-crystallin>hM4D* CNO (+) vs. *pc2>hM4D; βγ-crystallin>hM4D* CNO (-) | * | 0,0237 |
| | | | *βγ-crystallin>hM4D* CNO (+) vs. *pc2>hM4D; βγ-crystallin>hM4D* CNO (+) | ** | 0,0026 |
| | | | *pc2>hM4D; βγ-crystallin>hM4D* CNO (-)vs. *pc2>hM4D; βγ-crystallin>hM4D* CNO (+) | *** | 0,0006 |
| | | 60 min | *pc2>hM4D* CNO (-) vs. *pc2>hM4D* CNO (+) | **** | <0,0001 |
| | | | *pc2>hM4D* CNO (-) vs. *βγ-crystallin>hM4D* CNO (-) | ns | >0,9999 |
| | | | *pc2>hM4D* CNO (-) vs. *βγ-crystallin>hM4D* CNO (+) | *** | 0,0005 |
| | | | *pc2>hM4D* CNO (-) vs. *pc2>hM4D; βγ-crystallin>hM4D* CNO (-) | ns | 0,9360 |
| | | | *pc2>hM4D* CNO (-) vs. *pc2>hM4D; βγ-crystallin>hM4D* CNO (+) | **** | <0,0001 |
| | | | *pc2>hM4D* CNO (+) vs. *βγ-crystallin>hM4D* CNO (-) | **** | <0,0001 |
| | | | *pc2>hM4D* CNO (+) vs. *βγ-crystallin>hM4D* CNO (+) | ns | 0,0617 |
| | | | *pc2>hM4D* CNO (+) vs. *pc2>hM4D; βγ-crystallin>hM4D* CNO (-) | ** | 0,0030 |
| | | | *pc2>hM4D* CNO (+) vs. *pc2>hM4D; βγ-crystallin>hM4D* CNO (+) | ** | 0,0083 |
| | | | *βγ-crystallin>hM4D* CNO (-) vs. *βγ-crystallin>hM4D* CNO (+) | *** | 0,0009 |
| | | | *βγ-crystallin>hM4D* CNO (-) vs. *pc2>hM4D; βγ-crystallin>hM4D* CNO (-) | ns | 0,9140 |
| | | | *βγ-crystallin>hM4D* CNO CNO (-) vs. *pc2>hM4D; βγ-crystallin>hM4D* CNO (+) | **** | <0,0001 |
| | | | *βγ-crystallin>hM4D* CNO (+) vs. *pc2>hM4D; βγ-crystallin>hM4D* CNO (-) | ns | 0,0671 |
| | | | *βγ-crystallin>hM4D* CNO (+) vs. *pc2>hM4D; βγ-crystallin>hM4D* CNO (+) | **** | <0,0001 |
| | | | *pc2>hM4D; βγ-crystallin>hM4D* CNO (-)vs. *pc2>hM4D; βγ-crystallin>hM4D* CNO (+) | *** | 0,0002 |
| | | 90 min | *pc2>hM4D* CNO (-) vs. *pc2>hM4D* CNO (+) | *** | 0,0001 |
| | | | *pc2>hM4D* CNO (-) vs. *βγ-crystallin>hM4D* CNO (-) | ns | 0,9589 |
| | | | *pc2>hM4D* CNO (-) vs. *βγ-crystallin>hM4D* CNO (+) | ** | 0,0025 |
| | | | *pc2>hM4D* CNO (-) vs. *pc2>hM4D; βγ-crystallin>hM4D* CNO (-) | ns | 0,9450 |
| | | | *pc2>hM4D* CNO (-) vs. *pc2>hM4D; βγ-crystallin>hM4D* CNO (+) | **** | <0,0001 |

| | | | | | |
|---|---|---|---|---|---|
| | | | *pc2>hM4D* CNO (+) vs. *βγ-crystallin>hM4D* CNO (-) | **** | <0,0001 |
| | | | *pc2>hM4D* CNO (+) vs. *βγ-crystallin>hM4D* CNO (+) | * | 0,0175 |
| | | | *pc2>hM4D* CNO (+) vs. *pc2>hM4D; βγ-crystallin>hM4D* CNO (-) | *** | 0,0001 |
| | | | *pc2>hM4D* CNO (+) vs. *pc2>hM4D; βγ-crystallin>hM4D* CNO (+) | ** | 0,0038 |
| | | | *βγ-crystallin>hM4D* CNO (-) vs. *βγ-crystallin>hM4D* CNO (+) | *** | 0,0002 |
| | | | *βγ-crystallin>hM4D* CNO (-) vs. *pc2>hM4D; βγ-crystallin>hM4D* CNO (-) | ns | 0,7705 |
| | | | *βγ-crystallin>hM4D* CNO CNO (-) vs. *pc2>hM4D; βγ-crystallin>hM4D* CNO (+) | **** | <0,0001 |
| | | | *βγ-crystallin>hM4D* CNO (+) vs. *pc2>hM4D; βγ-crystallin>hM4D* CNO (-) | * | 0,0113 |
| | | | *βγ-crystallin>hM4D* CNO (+) vs. *pc2>hM4D; βγ-crystallin>hM4D* CNO (+) | **** | <0,0001 |
| | | | *pc2>hM4D; βγ-crystallin>hM4D* CNO (-)vs. *pc2>hM4D; βγ-crystallin>hM4D* CNO (+) | **** | <0,0001 |
| | | 120 min | *pc2>hM4D* CNO (-) vs. *pc2>hM4D* CNO (+) | **** | <0,0001 |
| | | | *pc2>hM4D* CNO (-) vs. *βγ-crystallin>hM4D* CNO (-) | ns | 0,4536 |
| | | | *pc2>hM4D* CNO (-) vs. *βγ-crystallin>hM4D* CNO (+) | ** | 0,0032 |
| | | | *pc2>hM4D* CNO (-) vs. *pc2>hM4D; βγ-crystallin>hM4D* CNO (-) | ns | 0,9110 |
| | | | *pc2>hM4D* CNO (-) vs. *pc2>hM4D; βγ-crystallin>hM4D* CNO (+) | **** | <0,0001 |
| | | | *pc2>hM4D* CNO (+) vs. *βγ-crystallin>hM4D* CNO (-) | **** | <0,0001 |
| | | | *pc2>hM4D* CNO (+) vs. *βγ-crystallin>hM4D* CNO (+) | ** | 0,0021 |
| | | | *pc2>hM4D* CNO (+) vs. *pc2>hM4D; βγ-crystallin>hM4D* CNO (-) | **** | <0,0001 |
| | | | *pc2>hM4D* CNO (+) vs. *pc2>hM4D; βγ-crystallin>hM4D* CNO (+) | ns | 0,1891 |
| | | | *βγ-crystallin>hM4D* CNO (-) vs. *βγ-crystallin>hM4D* CNO (+) | ** | 0,0023 |
| | | | *βγ-crystallin>hM4D* CNO (-) vs. *pc2>hM4D; βγ-crystallin>hM4D* CNO (-) | ns | 0,3991 |
| | | | *βγ-crystallin>hM4D* CNO CNO (-) vs. *pc2>hM4D; βγ-crystallin>hM4D* CNO (+) | **** | <0,0001 |
| | | | *βγ-crystallin>hM4D* CNO (+) vs. *pc2>hM4D; βγ-crystallin>hM4D* CNO (-) | ** | 0,0024 |
| | | | *βγ-crystallin>hM4D* CNO (+) vs. *pc2>hM4D; βγ-crystallin>hM4D* CNO (+) | **** | <0,0001 |
| | | | *pc2>hM4D; βγ-crystallin>hM4D* CNO (-)vs. *pc2>hM4D; βγ-crystallin>hM4D* CNO (+) | **** | <0,0001 |
| | | 150 min | *pc2>hM4D* CNO (-) vs. *pc2>hM4D* CNO (+) | *** | 0,0003 |
| | | | *pc2>hM4D* CNO (-) vs. *βγ-crystallin>hM4D* CNO (-) | ns | 0,9970 |

| | | | | | |
|---|---|---|---|---|---|
| | | | *pc2>hM4D* CNO (-) vs. *βγ-crystallin>hM4D* CNO (+) | * | 0,0207 |
| | | | *pc2>hM4D* CNO (-) vs. *pc2>hM4D; βγ-crystallin>hM4D* CNO (-) | ns | 0,9007 |
| | | | *pc2>hM4D* CNO (-) vs. *pc2>hM4D; βγ-crystallin>hM4D* CNO (+) | *** | 0,0002 |
| | | | *pc2>hM4D* CNO (+) vs. *βγ-crystallin>hM4D* CNO (-) | *** | 0,0005 |
| | | | *pc2>hM4D* CNO (+) vs. *βγ-crystallin>hM4D* CNO (+) | * | 0,0372 |
| | | | *pc2>hM4D* CNO (+) vs. *pc2>hM4D; βγ-crystallin>hM4D* CNO (-) | **** | <0,0001 |
| | | | *pc2>hM4D* CNO (+) vs. *pc2>hM4D; βγ-crystallin>hM4D* CNO (+) | ns | 0,1402 |
| | | | *βγ-crystallin>hM4D* CNO (-) vs. *βγ-crystallin>hM4D* CNO (+) | ** | 0,0010 |
| | | | *βγ-crystallin>hM4D* CNO (-) vs. *pc2>hM4D; βγ-crystallin>hM4D* CNO (-) | ns | 0,9863 |
| | | | *βγ-crystallin>hM4D* CNO CNO (-) vs. *pc2>hM4D; βγ-crystallin>hM4D* CNO (+) | **** | <0,0001 |
| | | | *βγ-crystallin>hM4D* CNO (+) vs. *pc2>hM4D; βγ-crystallin>hM4D* CNO (-) | *** | 0,0006 |
| | | | *βγ-crystallin>hM4D* CNO (+) vs. *pc2>hM4D; βγ-crystallin>hM4D* CNO (+) | ** | 0,0017 |
| | | | *pc2>hM4D; βγ-crystallin>hM4D* CNO (-)vs. *pc2>hM4D; βγ-crystallin>hM4D* CNO (+) | **** | <0,0001 |
| | | 180 min | *pc2>hM4D* CNO (-) vs. *pc2>hM4D* CNO (+) | *** | 0,0006 |
| | | | *pc2>hM4D* CNO (-) vs. *βγ-crystallin>hM4D* CNO (-) | ns | 0,9911 |
| | | | *pc2>hM4D* CNO (-) vs. *βγ-crystallin>hM4D* CNO (+) | * | 0,0142 |
| | | | *pc2>hM4D* CNO (-) vs. *pc2>hM4D; βγ-crystallin>hM4D* CNO (-) | ns | 0,9693 |
| | | | *pc2>hM4D* CNO (-) vs. *pc2>hM4D; βγ-crystallin>hM4D* CNO (+) | *** | 0,0006 |
| | | | *pc2>hM4D* CNO (+) vs. *βγ-crystallin>hM4D* CNO (-) | ** | 0,0012 |
| | | | *pc2>hM4D* CNO (+) vs. *βγ-crystallin>hM4D* CNO (+) | ns | 0,1746 |
| | | | *pc2>hM4D* CNO (+) vs. *pc2>hM4D; βγ-crystallin>hM4D* CNO (-) | **** | <0,0001 |
| | | | *pc2>hM4D* CNO (+) vs. *pc2>hM4D; βγ-crystallin>hM4D* CNO (+) | ns | 0,9953 |
| | | | *βγ-crystallin>hM4D* CNO (-) vs. *βγ-crystallin>hM4D* CNO (+) | * | 0,0114 |
| | | | *βγ-crystallin>hM4D* CNO (-) vs. *pc2>hM4D; βγ-crystallin>hM4D* CNO (-) | ns | 0,9982 |
| | | | *βγ-crystallin>hM4D* CNO CNO (-) vs. *pc2>hM4D; βγ-crystallin>hM4D* CNO (+) | *** | 0,0002 |
| | | | *βγ-crystallin>hM4D* CNO (+) vs. *pc2>hM4D; βγ-crystallin>hM4D* CNO (-) | **** | <0,0001 |
| | | | *βγ-crystallin>hM4D* CNO (+) vs. *pc2>hM4D; βγ-crystallin>hM4D* CNO (+) | * | 0,0400 |
| | | | *pc2>hM4D; βγ-crystallin>hM4D* CNO (-)vs. *pc2>hM4D; βγ-crystallin>hM4D* CNO (+) | **** | <0,0001 |

| | | | | | |
|---|---|---|---|---|---|
| | | 210 min | *pc2>hM4D* CNO (-) vs. *pc2>hM4D* CNO (+) | ** | 0,0011 |
| | | | *pc2>hM4D* CNO (-) vs. *βγ-crystallin>hM4D* CNO (-) | ns | 0,9496 |
| | | | *pc2>hM4D* CNO (-) vs. *βγ-crystallin>hM4D* CNO (+) | * | 0,0234 |
| | | | *pc2>hM4D* CNO (-) vs. *pc2>hM4D; βγ-crystallin>hM4D* CNO (-) | ns | 0,9669 |
| | | | *pc2>hM4D* CNO (-) vs. *pc2>hM4D; βγ-crystallin>hM4D* CNO (+) | ** | 0,0014 |
| | | | *pc2>hM4D* CNO (+) vs. *βγ-crystallin>hM4D* CNO (-) | *** | 0,0007 |
| | | | *pc2>hM4D* CNO (+) vs. *βγ-crystallin>hM4D* CNO (+) | ns | 0,2853 |
| | | | *pc2>hM4D* CNO (+) vs. *pc2>hM4D; βγ-crystallin>hM4D* CNO (-) | **** | <0,0001 |
| | | | *pc2>hM4D* CNO (+) vs. *pc2>hM4D; βγ-crystallin>hM4D* CNO (+) | ns | 0,8921 |
| | | | *βγ-crystallin>hM4D* CNO (-) vs. *βγ-crystallin>hM4D* CNO (+) | * | 0,0169 |
| | | | *βγ-crystallin>hM4D* CNO (-) vs. *pc2>hM4D; βγ-crystallin>hM4D* CNO (-) | ns | >0,9999 |
| | | | *βγ-crystallin>hM4D* CNO CNO (-) vs. *pc2>hM4D; βγ-crystallin>hM4D* CNO (+) | *** | 0,0003 |
| | | | *βγ-crystallin>hM4D* CNO (+) vs. *pc2>hM4D; βγ-crystallin>hM4D* CNO (-) | **** | <0,0001 |
| | | | *βγ-crystallin>hM4D* CNO (+) vs. *pc2>hM4D; βγ-crystallin>hM4D* CNO (+) | ns | 0,0590 |
| | | | *pc2>hM4D; βγ-crystallin>hM4D* CNO (-)vs. *pc2>hM4D; βγ-crystallin>hM4D* CNO (+) | **** | <0,0001 |
| | | 240 min | *pc2>hM4D* CNO (-) vs. *pc2>hM4D* CNO (+) | ** | 0,0031 |
| | | | *pc2>hM4D* CNO (-) vs. *βγ-crystallin>hM4D* CNO (-) | ns | 0,9813 |
| | | | *pc2>hM4D* CNO (-) vs. *βγ-crystallin>hM4D* CNO (+) | * | 0,0236 |
| | | | *pc2>hM4D* CNO (-) vs. *pc2>hM4D; βγ-crystallin>hM4D* CNO (-) | ns | >0,9999 |
| | | | *pc2>hM4D* CNO (-) vs. *pc2>hM4D; βγ-crystallin>hM4D* CNO (+) | ** | 0,0058 |
| | | | *pc2>hM4D* CNO (+) vs. *βγ-crystallin>hM4D* CNO (-) | ** | 0,0033 |
| | | | *pc2>hM4D* CNO (+) vs. *βγ-crystallin>hM4D* CNO (+) | ns | 0,2345 |
| | | | *pc2>hM4D* CNO (+) vs. *pc2>hM4D; βγ-crystallin>hM4D* CNO (-) | **** | <0,0001 |
| | | | *pc2>hM4D* CNO (+) vs. *pc2>hM4D; βγ-crystallin>hM4D* CNO (+) | ns | 0,9700 |
| | | | *βγ-crystallin>hM4D* CNO (-) vs. *βγ-crystallin>hM4D* CNO (+) | * | 0,0156 |
| | | | *βγ-crystallin>hM4D* CNO (-) vs. *pc2>hM4D; βγ-crystallin>hM4D* CNO (-) | ns | 0,9935 |
| | | | *βγ-crystallin>hM4D* CNO CNO (-) vs. *pc2>hM4D; βγ-crystallin>hM4D* CNO (+) | ** | 0,0035 |
| | | | *βγ-crystallin>hM4D* CNO (+) vs. *pc2>hM4D; βγ-crystallin>hM4D* CNO (-) | **** | <0,0001 |

| | | | | | |
|---|---|---|---|---|---|
| | | | *βγ-crystallin>hM4D* CNO (+) vs. *pc2>hM4D; βγ-crystallin>hM4D* CNO (+) | ns | 0,1224 |
| | | | *pc2>hM4D; βγ-crystallin>hM4D* CNO (-)vs. *pc2>hM4D; βγ-crystallin>hM4D* CNO (+) | **** | <0,0001 |
| | | 270 min | *pc2>hM4D* CNO (-) vs. *pc2>hM4D* CNO (+) | ** | 0,0038 |
| | | | *pc2>hM4D* CNO (-) vs. *βγ-crystallin>hM4D* CNO (-) | ns | 0,9888 |
| | | | *pc2>hM4D* CNO (-) vs. *βγ-crystallin>hM4D* CNO (+) | * | 0,0206 |
| | | | *pc2>hM4D* CNO (-) vs. *pc2>hM4D; βγ-crystallin>hM4D* CNO (-) | ns | >0,9999 |
| | | | *pc2>hM4D* CNO (-) vs. *pc2>hM4D; βγ-crystallin>hM4D* CNO (+) | ** | 0,0059 |
| | | | *pc2>hM4D* CNO (+) vs. *βγ-crystallin>hM4D* CNO (-) | ** | 0,0022 |
| | | | *pc2>hM4D* CNO (+) vs. *βγ-crystallin>hM4D* CNO (+) | ns | 0,4783 |
| | | | *pc2>hM4D* CNO (+) vs. *pc2>hM4D; βγ-crystallin>hM4D* CNO (-) | **** | <0,0001 |
| | | | *pc2>hM4D* CNO (+) vs. *pc2>hM4D; βγ-crystallin>hM4D* CNO (+) | ns | 0,8426 |
| | | | *βγ-crystallin>hM4D* CNO (-) vs. *βγ-crystallin>hM4D* CNO (+) | ** | 0,0064 |
| | | | *βγ-crystallin>hM4D* CNO (-) vs. *pc2>hM4D; βγ-crystallin>hM4D* CNO (-) | ns | 0,9391 |
| | | | *βγ-crystallin>hM4D* CNO CNO (-) vs. *pc2>hM4D; βγ-crystallin>hM4D* CNO (+) | ** | 0,0014 |
| | | | *βγ-crystallin>hM4D* CNO (+) vs. *pc2>hM4D; βγ-crystallin>hM4D* CNO (-) | **** | <0,0001 |
| | | | *βγ-crystallin>hM4D* CNO (+) vs. *pc2>hM4D; βγ-crystallin>hM4D* CNO (+) | ns | 0,3677 |
| | | | *pc2>hM4D; βγ-crystallin>hM4D* CNO (-)vs. *pc2>hM4D; βγ-crystallin>hM4D* CNO (+) | **** | <0,0001 |
| | | 300 min | *pc2>hM4D* CNO (-) vs. *pc2>hM4D* CNO (+) | ** | 0,0030 |
| | | | *pc2>hM4D* CNO (-) vs. *βγ-crystallin>hM4D* CNO (-) | ns | 0,9943 |
| | | | *pc2>hM4D* CNO (-) vs. *βγ-crystallin>hM4D* CNO (+) | * | 0,0126 |
| | | | *pc2>hM4D* CNO (-) vs. *pc2>hM4D; βγ-crystallin>hM4D* CNO (-) | ns | >0,9999 |
| | | | *pc2>hM4D* CNO (-) vs. *pc2>hM4D; βγ-crystallin>hM4D* CNO (+) | ** | 0,0025 |
| | | | *pc2>hM4D* CNO (+) vs. *βγ-crystallin>hM4D* CNO (-) | ** | 0,0012 |
| | | | *pc2>hM4D* CNO (+) vs. *βγ-crystallin>hM4D* CNO (+) | ns | 0,7699 |
| | | | *pc2>hM4D* CNO (+) vs. *pc2>hM4D; βγ-crystallin>hM4D* CNO (-) | **** | <0,0001 |
| | | | *pc2>hM4D* CNO (+) vs. *pc2>hM4D; βγ-crystallin>hM4D* CNO (+) | ns | >0,9999 |
| | | | *βγ-crystallin>hM4D* CNO (-) vs. *βγ-crystallin>hM4D* CNO (+) | ** | 0,0054 |
| | | | *βγ-crystallin>hM4D* CNO (-) vs. *pc2>hM4D; βγ-crystallin>hM4D* CNO (-) | ns | 0,9812 |

| | | | | | |
|---|---|---|---|---|---|
| | | | *βγ-crystallin>hM4D* CNO CNO (-) vs. *pc2>hM4D; βγ-crystallin>hM4D* CNO (+) | *** | 0,0004 |
| | | | *βγ-crystallin>hM4D* CNO (+) vs. *pc2>hM4D; βγ-crystallin>hM4D* CNO (-) | **** | <0,0001 |
| | | | *βγ-crystallin>hM4D* CNO (+) vs. *pc2>hM4D; βγ-crystallin>hM4D* CNO (+) | ns | 0,6042 |
| | | | *pc2>hM4D; βγ-crystallin>hM4D* CNO (-)vs. *pc2>hM4D; βγ-crystallin>hM4D* CNO (+) | **** | <0,0001 |
| | | 330 min | *pc2>hM4D* CNO (-) vs. *pc2>hM4D* CNO (+) | ** | 0,0029 |
| | | | *pc2>hM4D* CNO (-) vs. *βγ-crystallin>hM4D* CNO (-) | ns | 0,9939 |
| | | | *pc2>hM4D* CNO (-) vs. *βγ-crystallin>hM4D* CNO (+) | ** | 0,0064 |
| | | | *pc2>hM4D* CNO (-) vs. *pc2>hM4D; βγ-crystallin>hM4D* CNO (-) | ns | 0,9996 |
| | | | *pc2>hM4D* CNO (-) vs. *pc2>hM4D; βγ-crystallin>hM4D* CNO (+) | ** | 0,0035 |
| | | | *pc2>hM4D* CNO (+) vs. *βγ-crystallin>hM4D* CNO (-) | ** | 0,0011 |
| | | | *pc2>hM4D* CNO (+) vs. *βγ-crystallin>hM4D* CNO (+) | ns | 0,9741 |
| | | | *pc2>hM4D* CNO (+) vs. *pc2>hM4D; βγ-crystallin>hM4D* CNO (-) | **** | <0,0001 |
| | | | *pc2>hM4D* CNO (+) vs. *pc2>hM4D; βγ-crystallin>hM4D* CNO (+) | ns | >0,9999 |
| | | | *βγ-crystallin>hM4D* CNO (-) vs. *βγ-crystallin>hM4D* CNO (+) | ** | 0,0039 |
| | | | *βγ-crystallin>hM4D* CNO (-) vs. *pc2>hM4D; βγ-crystallin>hM4D* CNO (-) | ns | 0,9712 |
| | | | *βγ-crystallin>hM4D* CNO CNO (-) vs. *pc2>hM4D; βγ-crystallin>hM4D* CNO (+) | *** | 0,0010 |
| | | | *βγ-crystallin>hM4D* CNO (+) vs. *pc2>hM4D; βγ-crystallin>hM4D* CNO (-) | **** | <0,0001 |
| | | | *βγ-crystallin>hM4D* CNO (+) vs. *pc2>hM4D; βγ-crystallin>hM4D* CNO (+) | ns | 0,9140 |
| | | | *pc2>hM4D; βγ-crystallin>hM4D* CNO (-)vs. *pc2>hM4D; βγ-crystallin>hM4D* CNO (+) | **** | <0,0001 |
| | | 360 min | *pc2>hM4D* CNO (-) vs. *pc2>hM4D* CNO (+) | ** | 0,0034 |
| | | | *pc2>hM4D* CNO (-) vs. *βγ-crystallin>hM4D* CNO (-) | ns | >0,9999 |
| | | | *pc2>hM4D* CNO (-) vs. *βγ-crystallin>hM4D* CNO (+) | ** | 0,0034 |
| | | | *pc2>hM4D* CNO (-) vs. *pc2>hM4D; βγ-crystallin>hM4D* CNO (-) | ns | 0,9973 |
| | | | *pc2>hM4D* CNO (-) vs. *pc2>hM4D; βγ-crystallin>hM4D* CNO (+) | ** | 0,0019 |
| | | | *pc2>hM4D* CNO (+) vs. *βγ-crystallin>hM4D* CNO (-) | *** | 0,0008 |
| | | | *pc2>hM4D* CNO (+) vs. *βγ-crystallin>hM4D* CNO (+) | ns | 0,9980 |
| | | | *pc2>hM4D* CNO (+) vs. *pc2>hM4D; βγ-crystallin>hM4D* CNO (-) | **** | <0,0001 |

| | | | | | |
|---|---|---|---|---|---|
| | | | *pc2>hM4D* CNO (+) vs. *pc2>hM4D; βγ-crystallin>hM4D* CNO (+) | ns | 0,9880 |
| | | | *βγ-crystallin>hM4D* CNO (-) vs. *βγ-crystallin>hM4D* CNO (+) | *** | 0,0009 |
| | | | *βγ-crystallin>hM4D* CNO (-) vs. *pc2>hM4D; βγ-crystallin>hM4D* CNO (-) | ns | 0,9919 |
| | | | *βγ-crystallin>hM4D* CNO CNO (-) vs. *pc2>hM4D; βγ-crystallin>hM4D* CNO (+) | *** | 0,0006 |
| | | | *βγ-crystallin>hM4D* CNO (+) vs. *pc2>hM4D; βγ-crystallin>hM4D* CNO (-) | **** | <0,0001 |
| | | | *βγ-crystallin>hM4D* CNO (+) vs. *pc2>hM4D; βγ-crystallin>hM4D* CNO (+) | ns | 0,9994 |
| | | | *pc2>hM4D; βγ-crystallin>hM4D* CNO (-)vs. *pc2>hM4D; βγ-crystallin>hM4D* CNO (+) | **** | <0,0001 |
| | | 390 min | *pc2>hM4D* CNO (-) vs. *pc2>hM4D* CNO (+) | ** | 0,0035 |
| | | | *pc2>hM4D* CNO (-) vs. *βγ-crystallin>hM4D* CNO (-) | ns | 0,9998 |
| | | | *pc2>hM4D* CNO (-) vs. *βγ-crystallin>hM4D* CNO (+) | ** | 0,0033 |
| | | | *pc2>hM4D* CNO (-) vs. *pc2>hM4D; βγ-crystallin>hM4D* CNO (-) | ns | 0,9958 |
| | | | *pc2>hM4D* CNO (-) vs. *pc2>hM4D; βγ-crystallin>hM4D* CNO (+) | ** | 0,0011 |
| | | | *pc2>hM4D* CNO (+) vs. *βγ-crystallin>hM4D* CNO (-) | *** | 0,0008 |
| | | | *pc2>hM4D* CNO (+) vs. *βγ-crystallin>hM4D* CNO (+) | ns | 0,9970 |
| | | | *pc2>hM4D* CNO (+) vs. *pc2>hM4D; βγ-crystallin>hM4D* CNO (-) | **** | <0,0001 |
| | | | *pc2>hM4D* CNO (+) vs. *pc2>hM4D; βγ-crystallin>hM4D* CNO (+) | ns | 0,8261 |
| | | | *βγ-crystallin>hM4D* CNO (-) vs. *βγ-crystallin>hM4D* CNO (+) | *** | 0,0010 |
| | | | *βγ-crystallin>hM4D* CNO (-) vs. *pc2>hM4D; βγ-crystallin>hM4D* CNO (-) | ns | 0,9980 |
| | | | *βγ-crystallin>hM4D* CNO CNO (-) vs. *pc2>hM4D; βγ-crystallin>hM4D* CNO (+) | *** | 0,0003 |
| | | | *βγ-crystallin>hM4D* CNO (+) vs. *pc2>hM4D; βγ-crystallin>hM4D* CNO (-) | **** | <0,0001 |
| | | | *βγ-crystallin>hM4D* CNO (+) vs. *pc2>hM4D; βγ-crystallin>hM4D* CNO (+) | ns | 0,9555 |
| | | | *pc2>hM4D; βγ-crystallin>hM4D* CNO (-)vs. *pc2>hM4D; βγ-crystallin>hM4D* CNO (+) | **** | <0,0001 |
| | | 420 min | *pc2>hM4D* CNO (-) vs. *pc2>hM4D* CNO (+) | ** | 0,0033 |
| | | | *pc2>hM4D* CNO (-) vs. *βγ-crystallin>hM4D* CNO (-) | ns | 0,9211 |
| | | | *pc2>hM4D* CNO (-) vs. *βγ-crystallin>hM4D* CNO (+) | ** | 0,0023 |
| | | | *pc2>hM4D* CNO (-) vs. *pc2>hM4D; βγ-crystallin>hM4D* CNO (-) | ns | 0,9899 |
| | | | *pc2>hM4D* CNO (-) vs. *pc2>hM4D; βγ-crystallin>hM4D* CNO (+) | *** | 0,0007 |

| | | | | | |
|---|---|---|---|---|---|
| | | | *pc2>hM4D* CNO (+) vs. *βγ-crystallin>hM4D* CNO (-) | *** | 0,0003 |
| | | | *pc2>hM4D* CNO (+) vs. *βγ-crystallin>hM4D* CNO (+) | ns | 0,9618 |
| | | | *pc2>hM4D* CNO (+) vs. *pc2>hM4D; βγ-crystallin>hM4D* CNO (-) | **** | <0,0001 |
| | | | *pc2>hM4D* CNO (+) vs. *pc2>hM4D; βγ-crystallin>hM4D* CNO (+) | ns | 0,4145 |
| | | | *βγ-crystallin>hM4D* CNO (-) vs. *βγ-crystallin>hM4D* CNO (+) | *** | 0,0003 |
| | | | *βγ-crystallin>hM4D* CNO (-) vs. *pc2>hM4D; βγ-crystallin>hM4D* CNO (-) | ns | >0,9999 |
| | | | *βγ-crystallin>hM4D* CNO CNO (-) vs. *pc2>hM4D; βγ-crystallin>hM4D* CNO (+) | **** | <0,0001 |
| | | | *βγ-crystallin>hM4D* CNO (+) vs. *pc2>hM4D; βγ-crystallin>hM4D* CNO (-) | **** | <0,0001 |
| | | | *βγ-crystallin>hM4D* CNO (+) vs. *pc2>hM4D; βγ-crystallin>hM4D* CNO (+) | ns | 0,8178 |
| | | | *pc2>hM4D; βγ-crystallin>hM4D* CNO (-)vs. *pc2>hM4D; βγ-crystallin>hM4D* CNO (+) | **** | <0,0001 |
| | | 450 min | *pc2>hM4D* CNO (-) vs. *pc2>hM4D* CNO (+) | ** | 0,0025 |
| | | | *pc2>hM4D* CNO (-) vs. *βγ-crystallin>hM4D* CNO (-) | ns | 0,4608 |
| | | | *pc2>hM4D* CNO (-) vs. *βγ-crystallin>hM4D* CNO (+) | ** | 0,0010 |
| | | | *pc2>hM4D* CNO (-) vs. *pc2>hM4D; βγ-crystallin>hM4D* CNO (-) | ns | 0,9885 |
| | | | *pc2>hM4D* CNO (-) vs. *pc2>hM4D; βγ-crystallin>hM4D* CNO (+) | *** | 0,0003 |
| | | | *pc2>hM4D* CNO (+) vs. *βγ-crystallin>hM4D* CNO (-) | *** | 0,0003 |
| | | | *pc2>hM4D* CNO (+) vs. *βγ-crystallin>hM4D* CNO (+) | ns | 0,9666 |
| | | | *pc2>hM4D* CNO (+) vs. *pc2>hM4D; βγ-crystallin>hM4D* CNO (-) | **** | <0,0001 |
| | | | *pc2>hM4D* CNO (+) vs. *pc2>hM4D; βγ-crystallin>hM4D* CNO (+) | ns | 0,3438 |
| | | | *βγ-crystallin>hM4D* CNO (-) vs. *βγ-crystallin>hM4D* CNO (+) | *** | 0,0001 |
| | | | *βγ-crystallin>hM4D* CNO (-) vs. *pc2>hM4D; βγ-crystallin>hM4D* CNO (-) | ns | 0,9999 |
| | | | *βγ-crystallin>hM4D* CNO CNO (-) vs. *pc2>hM4D; βγ-crystallin>hM4D* CNO (+) | **** | <0,0001 |
| | | | *βγ-crystallin>hM4D* CNO (+) vs. *pc2>hM4D; βγ-crystallin>hM4D* CNO (-) | **** | <0,0001 |
| | | | *βγ-crystallin>hM4D* CNO (+) vs. *pc2>hM4D; βγ-crystallin>hM4D* CNO (+) | ns | 0,7647 |
| | | | *pc2>hM4D; βγ-crystallin>hM4D* CNO (-)vs. *pc2>hM4D; βγ-crystallin>hM4D* CNO (+) | **** | <0,0001 |
| | | 480 min | *pc2>hM4D* CNO (-) vs. *pc2>hM4D* CNO (+) | ** | 0,0021 |
| | | | *pc2>hM4D* CNO (-) vs. *βγ-crystallin>hM4D* CNO (-) | ns | 0,1768 |

| | | | | | |
|---|---|---|---|---|---|
| | | | *pc2>hM4D* CNO (-) vs. *βγ-crystallin>hM4D* CNO (+) | *** | 0,0006 |
| | | | *pc2>hM4D* CNO (-) vs. *pc2>hM4D; βγ-crystallin>hM4D* CNO (-) | ns | 0,9718 |
| | | | *pc2>hM4D* CNO (-) vs. *pc2>hM4D; βγ-crystallin>hM4D* CNO (+) | *** | 0,0002 |
| | | | *pc2>hM4D* CNO (+) vs. *βγ-crystallin>hM4D* CNO (-) | *** | 0,0003 |
| | | | *pc2>hM4D* CNO (+) vs. *βγ-crystallin>hM4D* CNO (+) | ns | 0,9406 |
| | | | *pc2>hM4D* CNO (+) vs. *pc2>hM4D; βγ-crystallin>hM4D* CNO (-) | **** | <0,0001 |
| | | | *pc2>hM4D* CNO (+) vs. *pc2>hM4D; βγ-crystallin>hM4D* CNO (+) | ns | 0,1161 |
| | | | *βγ-crystallin>hM4D* CNO (-) vs. *βγ-crystallin>hM4D* CNO (+) | **** | <0,0001 |
| | | | *βγ-crystallin>hM4D* CNO (-) vs. *pc2>hM4D; βγ-crystallin>hM4D* CNO (-) | ns | >0,9999 |
| | | | *βγ-crystallin>hM4D* CNO CNO (-) vs. *pc2>hM4D; βγ-crystallin>hM4D* CNO (+) | **** | <0,0001 |
| | | | *βγ-crystallin>hM4D* CNO (+) vs. *pc2>hM4D; βγ-crystallin>hM4D* CNO (-) | **** | <0,0001 |
| | | | *βγ-crystallin>hM4D* CNO (+) vs. *pc2>hM4D; βγ-crystallin>hM4D* CNO (+) | ns | 0,5450 |
| | | | *pc2>hM4D; βγ-crystallin>hM4D* CNO (-)vs. *pc2>hM4D; βγ-crystallin>hM4D* CNO (+) | **** | <0,0001 |
| | | 510 min | *pc2>hM4D* CNO (-) vs. *pc2>hM4D* CNO (+) | ** | 0,0014 |
| | | | *pc2>hM4D* CNO (-) vs. *βγ-crystallin>hM4D* CNO (-) | ns | 0,3592 |
| | | | *pc2>hM4D* CNO (-) vs. *βγ-crystallin>hM4D* CNO (+) | *** | 0,0004 |
| | | | *pc2>hM4D* CNO (-) vs. *pc2>hM4D; βγ-crystallin>hM4D* CNO (-) | ns | 0,9312 |
| | | | *pc2>hM4D* CNO (-) vs. *pc2>hM4D; βγ-crystallin>hM4D* CNO (+) | **** | <0,0001 |
| | | | *pc2>hM4D* CNO (+) vs. *βγ-crystallin>hM4D* CNO (-) | *** | 0,0004 |
| | | | *pc2>hM4D* CNO (+) vs. *βγ-crystallin>hM4D* CNO (+) | ns | 0,9445 |
| | | | *pc2>hM4D* CNO (+) vs. *pc2>hM4D; βγ-crystallin>hM4D* CNO (-) | **** | <0,0001 |
| | | | *pc2>hM4D* CNO (+) vs. *pc2>hM4D; βγ-crystallin>hM4D* CNO (+) | ns | 0,0784 |
| | | | *βγ-crystallin>hM4D* CNO (-) vs. *βγ-crystallin>hM4D* CNO (+) | *** | 0,0001 |
| | | | *βγ-crystallin>hM4D* CNO (-) vs. *pc2>hM4D; βγ-crystallin>hM4D* CNO (-) | ns | 0,9995 |
| | | | *βγ-crystallin>hM4D* CNO CNO (-) vs. *pc2>hM4D; βγ-crystallin>hM4D* CNO (+) | **** | <0,0001 |
| | | | *βγ-crystallin>hM4D* CNO (+) vs. *pc2>hM4D; βγ-crystallin>hM4D* CNO (-) | **** | <0,0001 |
| | | | *βγ-crystallin>hM4D* CNO (+) vs. *pc2>hM4D; βγ-crystallin>hM4D* CNO (+) | ns | 0,5237 |
| | | | *pc2>hM4D; βγ-crystallin>hM4D* CNO (-)vs. *pc2>hM4D; βγ-crystallin>hM4D* CNO (+) | **** | <0,0001 |

| | | | | | |
|---|---|---|---|---|---|
| | | 540 min | *pc2>hM4D* CNO (-) vs. *pc2>hM4D* CNO (+) | ** | 0,0010 |
| | | | *pc2>hM4D* CNO (-) vs. *βγ-crystallin>hM4D* CNO (-) | ns | 0,4378 |
| | | | *pc2>hM4D* CNO (-) vs. *βγ-crystallin>hM4D* CNO (+) | *** | 0,0002 |
| | | | *pc2>hM4D* CNO (-) vs. *pc2>hM4D; βγ-crystallin>hM4D* CNO (-) | ns | 0,9124 |
| | | | *pc2>hM4D* CNO (-) vs. *pc2>hM4D; βγ-crystallin>hM4D* CNO (+) | **** | <0,0001 |
| | | | *pc2>hM4D* CNO (+) vs. *βγ-crystallin>hM4D* CNO (-) | *** | 0,0002 |
| | | | *pc2>hM4D* CNO (+) vs. *βγ-crystallin>hM4D* CNO (+) | ns | 0,9671 |
| | | | *pc2>hM4D* CNO (+) vs. *pc2>hM4D; βγ-crystallin>hM4D* CNO (-) | **** | <0,0001 |
| | | | *pc2>hM4D* CNO (+) vs. *pc2>hM4D; βγ-crystallin>hM4D* CNO (+) | ns | 0,0778 |
| | | | *βγ-crystallin>hM4D* CNO (-) vs. *βγ-crystallin>hM4D* CNO (+) | **** | <0,0001 |
| | | | *βγ-crystallin>hM4D* CNO (-) vs. *pc2>hM4D; βγ-crystallin>hM4D* CNO (-) | ns | 0,9990 |
| | | | *βγ-crystallin>hM4D* CNO CNO (-) vs. *pc2>hM4D; βγ-crystallin>hM4D* CNO (+) | **** | <0,0001 |
| | | | *βγ-crystallin>hM4D* CNO (+) vs. *pc2>hM4D; βγ-crystallin>hM4D* CNO (-) | **** | <0,0001 |
| | | | *βγ-crystallin>hM4D* CNO (+) vs. *pc2>hM4D; βγ-crystallin>hM4D* CNO (+) | ns | 0,3613 |
| | | | *pc2>hM4D; βγ-crystallin>hM4D* CNO (-)vs. *pc2>hM4D; βγ-crystallin>hM4D* CNO (+) | **** | <0,0001 |
| | | 570 min | *pc2>hM4D* CNO (-) vs. *pc2>hM4D* CNO (+) | *** | 0,0003 |
| | | | *pc2>hM4D* CNO (-) vs. *βγ-crystallin>hM4D* CNO (-) | ns | 0,5280 |
| | | | *pc2>hM4D* CNO (-) vs. *βγ-crystallin>hM4D* CNO (+) | **** | <0,0001 |
| | | | *pc2>hM4D* CNO (-) vs. *pc2>hM4D; βγ-crystallin>hM4D* CNO (-) | ns | 0,9494 |
| | | | *pc2>hM4D* CNO (-) vs. *pc2>hM4D; βγ-crystallin>hM4D* CNO (+) | **** | <0,0001 |
| | | | *pc2>hM4D* CNO (+) vs. *βγ-crystallin>hM4D* CNO (-) | *** | 0,0002 |
| | | | *pc2>hM4D* CNO (+) vs. *βγ-crystallin>hM4D* CNO (+) | ns | 0,9761 |
| | | | *pc2>hM4D* CNO (+) vs. *pc2>hM4D; βγ-crystallin>hM4D* CNO (-) | **** | <0,0001 |
| | | | *pc2>hM4D* CNO (+) vs. *pc2>hM4D; βγ-crystallin>hM4D* CNO (+) | ns | 0,0563 |
| | | | *βγ-crystallin>hM4D* CNO (-) vs. *βγ-crystallin>hM4D* CNO (+) | **** | <0,0001 |
| | | | *βγ-crystallin>hM4D* CNO (-) vs. *pc2>hM4D; βγ-crystallin>hM4D* CNO (-) | ns | >0,9999 |
| | | | *βγ-crystallin>hM4D* CNO CNO (-) vs. *pc2>hM4D; βγ-crystallin>hM4D* CNO (+) | **** | <0,0001 |
| | | | *βγ-crystallin>hM4D* CNO (+) vs. *pc2>hM4D; βγ-crystallin>hM4D* CNO (-) | **** | <0,0001 |

| | | | | | |
|---|---|---|---|---|---|
| | | | *βγ-crystallin>hM4D* CNO (+) vs. *pc2>hM4D; βγ-crystallin>hM4D* CNO (+) | ns | 0,2581 |
| | | | *pc2>hM4D; βγ-crystallin>hM4D* CNO (-)vs. *pc2>hM4D; βγ-crystallin>hM4D* CNO (+) | **** | <0,0001 |
| | | 600 min | *pc2>hM4D* CNO (-) vs. *pc2>hM4D* CNO (+) | *** | 0,0002 |
| | | | *pc2>hM4D* CNO (-) vs. *βγ-crystallin>hM4D* CNO (-) | ns | 0,9068 |
| | | | *pc2>hM4D* CNO (-) vs. *βγ-crystallin>hM4D* CNO (+) | **** | <0,0001 |
| | | | *pc2>hM4D* CNO (-) vs. *pc2>hM4D; βγ-crystallin>hM4D* CNO (-) | ns | 0,9991 |
| | | | *pc2>hM4D* CNO (-) vs. *pc2>hM4D; βγ-crystallin>hM4D* CNO (+) | **** | <0,0001 |
| | | | *pc2>hM4D* CNO (+) vs. *βγ-crystallin>hM4D* CNO (-) | *** | 0,0002 |
| | | | *pc2>hM4D* CNO (+) vs. *βγ-crystallin>hM4D* CNO (+) | ns | 0,9879 |
| | | | *pc2>hM4D* CNO (+) vs. *pc2>hM4D; βγ-crystallin>hM4D* CNO (-) | **** | <0,0001 |
| | | | *pc2>hM4D* CNO (+) vs. *pc2>hM4D; βγ-crystallin>hM4D* CNO (+) | * | 0,0190 |
| | | | *βγ-crystallin>hM4D* CNO (-) vs. *βγ-crystallin>hM4D* CNO (+) | **** | <0,0001 |
| | | | *βγ-crystallin>hM4D* CNO (-) vs. *pc2>hM4D; βγ-crystallin>hM4D* CNO (-) | ns | >0,9999 |
| | | | *βγ-crystallin>hM4D* CNO CNO (-) vs. *pc2>hM4D; βγ-crystallin>hM4D* CNO (+) | **** | <0,0001 |
| | | | *βγ-crystallin>hM4D* CNO (+) vs. *pc2>hM4D; βγ-crystallin>hM4D* CNO (-) | **** | <0,0001 |
| | | | *βγ-crystallin>hM4D* CNO (+) vs. *pc2>hM4D; βγ-crystallin>hM4D* CNO (+) | ns | 0,1051 |
| | | | *pc2>hM4D; βγ-crystallin>hM4D* CNO (-)vs. *pc2>hM4D; βγ-crystallin>hM4D* CNO (+) | **** | <0,0001 |

table S19 Statistics for settlement rates analysis for *pc2>hM4D*, *βγ-crystallin>hM4D* single and *pc2>hM4D*; *βγ-crystallin>hM4D* double transgenics in the presence of 200μM Carvacrol

| Figure & Panel | Test | Data point | Comparison | P Value summary | P Value |
|---|---|---|---|---|---|
| Fig. 5C | Two-way RM ANOVA | | | | |
| | | | Time | **** | <0,0001 |
| | | | Fraction Settled | **** | <0,0001 |
| | | | Time x Fraction Settled | **** | <0,0001 |
| | Tukey's multiple comparisons test | | | | |
| | | 30 min | *pc2>hM4D* CNO (-) vs. *pc2>hM4D* CNO (+) | * | 0,0176 |
| | | | *pc2>hM4D* CNO (-) vs. *βγ-crystallin>hM4D* CNO (-) | ns | 0,9998 |

| | | | | | |
|---|---|---|---|---|---|
| | | | *pc2>hM4D* CNO (-) vs. *βγ-crystallin>hM4D* CNO (+) | * | 0,0199 |
| | | | *pc2>hM4D* CNO (-) vs. *pc2>hM4D; βγ-crystallin>hM4D* CNO (-) | ns | 0,9998 |
| | | | *pc2>hM4D* CNO (-) vs. *pc2>hM4D; βγ-crystallin>hM4D* CNO (+) | ns | 0,6729 |
| | | | *pc2>hM4D* CNO (+) vs. *βγ-crystallin>hM4D* CNO (-) | * | 0,0181 |
| | | | *pc2>hM4D* CNO (+) vs. *βγ-crystallin>hM4D* CNO (+) | ns | 0,9916 |
| | | | *pc2>hM4D* CNO (+) vs. *pc2>hM4D; βγ-crystallin>hM4D* CNO (-) | * | 0,0272 |
| | | | *pc2>hM4D* CNO (+) vs. *pc2>hM4D; βγ-crystallin>hM4D* CNO (+) | ns | 0,1419 |
| | | | *βγ-crystallin>hM4D* CNO (-) vs. *βγ-crystallin>hM4D* CNO (+) | ** | 0,0089 |
| | | | *βγ-crystallin>hM4D* CNO (-) vs. *pc2>hM4D; βγ-crystallin>hM4D* CNO (-) | ns | 0,9984 |
| | | | *βγ-crystallin>hM4D* CNO CNO (-) vs. *pc2>hM4D; βγ-crystallin>hM4D* CNO (+) | ns | 0,9401 |
| | | | *βγ-crystallin>hM4D* CNO (+) vs. *pc2>hM4D; βγ-crystallin>hM4D* CNO (-) | * | 0,0285 |
| | | | *βγ-crystallin>hM4D* CNO (+) vs. *pc2>hM4D; βγ-crystallin>hM4D* CNO (+) | ns | 0,1338 |
| | | | *pc2>hM4D; βγ-crystallin>hM4D* CNO (-)vs. *pc2>hM4D; βγ-crystallin>hM4D* CNO (+) | ns | 0,1698 |
| | | 60 min | *pc2>hM4D* CNO (-) vs. *pc2>hM4D* CNO (+) | ns | 0,0819 |
| | | | *pc2>hM4D* CNO (-) vs. *βγ-crystallin>hM4D* CNO (-) | ns | 0,9401 |
| | | | *pc2>hM4D* CNO (-) vs. *βγ-crystallin>hM4D* CNO (+) | ns | 0,0574 |
| | | | *pc2>hM4D* CNO (-) vs. *pc2>hM4D; βγ-crystallin>hM4D* CNO (-) | ns | >0,9999 |
| | | | *pc2>hM4D* CNO (-) vs. *pc2>hM4D; βγ-crystallin>hM4D* CNO (+) | ns | 0,8687 |
| | | | *pc2>hM4D* CNO (+) vs. *βγ-crystallin>hM4D* CNO (-) | * | 0,0296 |
| | | | *pc2>hM4D* CNO (+) vs. *βγ-crystallin>hM4D* CNO (+) | ns | >0,9999 |
| | | | *pc2>hM4D* CNO (+) vs. *pc2>hM4D; βγ-crystallin>hM4D* CNO (-) | ns | 0,0837 |
| | | | *pc2>hM4D* CNO (+) vs. *pc2>hM4D; βγ-crystallin>hM4D* CNO (+) | ns | 0,2646 |
| | | | *βγ-crystallin>hM4D* CNO (-) vs. *βγ-crystallin>hM4D* CNO (+) | ** | 0,0094 |
| | | | *βγ-crystallin>hM4D* CNO (-) vs. *pc2>hM4D; βγ-crystallin>hM4D* CNO (-) | ns | 0,8256 |
| | | | *βγ-crystallin>hM4D* CNO CNO (-) vs. *pc2>hM4D; βγ-crystallin>hM4D* CNO (+) | ns | 0,7109 |
| | | | *βγ-crystallin>hM4D* CNO (+) vs. *pc2>hM4D; βγ-crystallin>hM4D* CNO (-) | ns | 0,0510 |
| | | | *βγ-crystallin>hM4D* CNO (+) vs. *pc2>hM4D; βγ-crystallin>hM4D* CNO (+) | ns | 0,1827 |
| | | | *pc2>hM4D; βγ-crystallin>hM4D* CNO (-)vs. *pc2>hM4D; βγ-crystallin>hM4D* CNO (+) | ns | 0,9931 |

| | | | | | |
|---|---|---|---|---|---|
| | | 90 min | *pc2>hM4D* CNO (-) vs. *pc2>hM4D* CNO (+) | * | 0,0395 |
| | | | *pc2>hM4D* CNO (-) vs. *βγ-crystallin>hM4D* CNO (-) | ns | 0,6532 |
| | | | *pc2>hM4D* CNO (-) vs. *βγ-crystallin>hM4D* CNO (+) | ns | 0,0608 |
| | | | *pc2>hM4D* CNO (-) vs. *pc2>hM4D; βγ-crystallin>hM4D* CNO (-) | ns | 0,9998 |
| | | | *pc2>hM4D* CNO (-) vs. *pc2>hM4D; βγ-crystallin>hM4D* CNO (+) | * | 0,0268 |
| | | | *pc2>hM4D* CNO (+) vs. *βγ-crystallin>hM4D* CNO (-) | ** | 0,0041 |
| | | | *pc2>hM4D* CNO (+) vs. *βγ-crystallin>hM4D* CNO (+) | ns | >0,9999 |
| | | | *pc2>hM4D* CNO (+) vs. *pc2>hM4D; βγ-crystallin>hM4D* CNO (-) | * | 0,0330 |
| | | | *pc2>hM4D* CNO (+) vs. *pc2>hM4D; βγ-crystallin>hM4D* CNO (+) | ns | 0,7241 |
| | | | *βγ-crystallin>hM4D* CNO (-) vs. *βγ-crystallin>hM4D* CNO (+) | ** | 0,0012 |
| | | | *βγ-crystallin>hM4D* CNO (-) vs. *pc2>hM4D; βγ-crystallin>hM4D* CNO (-) | ns | 0,2501 |
| | | | *βγ-crystallin>hM4D* CNO CNO (-) vs. *pc2>hM4D; βγ-crystallin>hM4D* CNO (+) | * | 0,0201 |
| | | | *βγ-crystallin>hM4D* CNO (+) vs. *pc2>hM4D; βγ-crystallin>hM4D* CNO (-) | * | 0,0300 |
| | | | *βγ-crystallin>hM4D* CNO (+) vs. *pc2>hM4D; βγ-crystallin>hM4D* CNO (+) | ns | 0,6729 |
| | | | *pc2>hM4D; βγ-crystallin>hM4D* CNO (-)vs. *pc2>hM4D; βγ-crystallin>hM4D* CNO (+) | ns | 0,2811 |
| | | 120 min | *pc2>hM4D* CNO (-) vs. *pc2>hM4D* CNO (+) | * | 0,0111 |
| | | | *pc2>hM4D* CNO (-) vs. *βγ-crystallin>hM4D* CNO (-) | ns | 0,9991 |
| | | | *pc2>hM4D* CNO (-) vs. *βγ-crystallin>hM4D* CNO (+) | ns | 0,1169 |
| | | | *pc2>hM4D* CNO (-) vs. *pc2>hM4D; βγ-crystallin>hM4D* CNO (-) | ns | 0,8600 |
| | | | *pc2>hM4D* CNO (-) vs. *pc2>hM4D; βγ-crystallin>hM4D* CNO (+) | **** | <0,0001 |
| | | | *pc2>hM4D* CNO (+) vs. *βγ-crystallin>hM4D* CNO (-) | ** | 0,0026 |
| | | | *pc2>hM4D* CNO (+) vs. *βγ-crystallin>hM4D* CNO (+) | ns | 0,9864 |
| | | | *pc2>hM4D* CNO (+) vs. *pc2>hM4D; βγ-crystallin>hM4D* CNO (-) | *** | 0,0008 |
| | | | *pc2>hM4D* CNO (+) vs. *pc2>hM4D; βγ-crystallin>hM4D* CNO (+) | ns | >0,9999 |
| | | | *βγ-crystallin>hM4D* CNO (-) vs. *βγ-crystallin>hM4D* CNO (+) | * | 0,0335 |
| | | | *βγ-crystallin>hM4D* CNO (-) vs. *pc2>hM4D; βγ-crystallin>hM4D* CNO (-) | ns | 0,9486 |
| | | | *βγ-crystallin>hM4D* CNO CNO (-) vs. *pc2>hM4D; βγ-crystallin>hM4D* CNO (+) | *** | 0,0003 |
| | | | *βγ-crystallin>hM4D* CNO (+) vs. *pc2>hM4D; βγ-crystallin>hM4D* CNO (-) | ns | 0,1238 |

| | | | | | |
|---|---|---|---|---|---|
| | | | *βγ-crystallin>hM4D* CNO (+) vs. *pc2>hM4D; βγ-crystallin>hM4D* CNO (+) | ns | 0,9951 |
| | | | *pc2>hM4D; βγ-crystallin>hM4D* CNO (-)vs. *pc2>hM4D; βγ-crystallin>hM4D* CNO (+) | ** | 0,0055 |
| | | 150 min | *pc2>hM4D* CNO (-) vs. *pc2>hM4D* CNO (+) | ** | 0,0100 |
| | | | *pc2>hM4D* CNO (-) vs. *βγ-crystallin>hM4D* CNO (-) | ns | 0,9704 |
| | | | *pc2>hM4D* CNO (-) vs. *βγ-crystallin>hM4D* CNO (+) | ns | 0,0667 |
| | | | *pc2>hM4D* CNO (-) vs. *pc2>hM4D; βγ-crystallin>hM4D* CNO (-) | ns | 0,6515 |
| | | | *pc2>hM4D* CNO (-) vs. *pc2>hM4D; βγ-crystallin>hM4D* CNO (+) | *** | 0,0001 |
| | | | *pc2>hM4D* CNO (+) vs. *βγ-crystallin>hM4D* CNO (-) | ** | 0,0017 |
| | | | *pc2>hM4D* CNO (+) vs. *βγ-crystallin>hM4D* CNO (+) | ns | >0,9999 |
| | | | *pc2>hM4D* CNO (+) vs. *pc2>hM4D; βγ-crystallin>hM4D* CNO (-) | ** | 0,0042 |
| | | | *pc2>hM4D* CNO (+) vs. *pc2>hM4D; βγ-crystallin>hM4D* CNO (+) | ns | 0,8957 |
| | | | *βγ-crystallin>hM4D* CNO (-) vs. *βγ-crystallin>hM4D* CNO (+) | * | 0,0346 |
| | | | *βγ-crystallin>hM4D* CNO (-) vs. *pc2>hM4D; βγ-crystallin>hM4D* CNO (-) | ns | 0,9311 |
| | | | *βγ-crystallin>hM4D* CNO CNO (-) vs. *pc2>hM4D; βγ-crystallin>hM4D* CNO (+) | ** | 0,0011 |
| | | | *βγ-crystallin>hM4D* CNO (+) vs. *pc2>hM4D; βγ-crystallin>hM4D* CNO (-) | ns | 0,0782 |
| | | | *βγ-crystallin>hM4D* CNO (+) vs. *pc2>hM4D; βγ-crystallin>hM4D* CNO (+) | ns | 0,9203 |
| | | | *pc2>hM4D; βγ-crystallin>hM4D* CNO (-)vs. *pc2>hM4D; βγ-crystallin>hM4D* CNO (+) | *** | 0,0001 |
| | | 180 min | *pc2>hM4D* CNO (-) vs. *pc2>hM4D* CNO (+) | * | 0,0388 |
| | | | *pc2>hM4D* CNO (-) vs. *βγ-crystallin>hM4D* CNO (-) | ns | 0,7576 |
| | | | *pc2>hM4D* CNO (-) vs. *βγ-crystallin>hM4D* CNO (+) | ns | 0,0520 |
| | | | *pc2>hM4D* CNO (-) vs. *pc2>hM4D; βγ-crystallin>hM4D* CNO (-) | ns | 0,9320 |
| | | | *pc2>hM4D* CNO (-) vs. *pc2>hM4D; βγ-crystallin>hM4D* CNO (+) | *** | 0,0002 |
| | | | *pc2>hM4D* CNO (+) vs. *βγ-crystallin>hM4D* CNO (-) | * | 0,0157 |
| | | | *pc2>hM4D* CNO (+) vs. *βγ-crystallin>hM4D* CNO (+) | ns | >0,9999 |
| | | | *pc2>hM4D* CNO (+) vs. *pc2>hM4D; βγ-crystallin>hM4D* CNO (-) | * | 0,0194 |
| | | | *pc2>hM4D* CNO (+) vs. *pc2>hM4D; βγ-crystallin>hM4D* CNO (+) | ns | 0,4401 |
| | | | *βγ-crystallin>hM4D* CNO (-) vs. *βγ-crystallin>hM4D* CNO (+) | * | 0,0443 |
| | | | *βγ-crystallin>hM4D* CNO (-) vs. *pc2>hM4D; βγ-crystallin>hM4D* CNO (-) | ns | 0,9873 |

| | | | | | |
|---|---|---|---|---|---|
| | | | *βγ-crystallin>hM4D* CNO CNO (-) vs. *pc2>hM4D; βγ-crystallin>hM4D* CNO (+) | *** | 0,0003 |
| | | | *βγ-crystallin>hM4D* CNO (+) vs. *pc2>hM4D; βγ-crystallin>hM4D* CNO (-) | * | 0,0411 |
| | | | *βγ-crystallin>hM4D* CNO (+) vs. *pc2>hM4D; βγ-crystallin>hM4D* CNO (+) | ns | 0,6303 |
| | | | *pc2>hM4D; βγ-crystallin>hM4D* CNO (-)vs. *pc2>hM4D; βγ-crystallin>hM4D* CNO (+) | **** | <0,0001 |
| | | 210 min | *pc2>hM4D* CNO (-) vs. *pc2>hM4D* CNO (+) | * | 0,0383 |
| | | | *pc2>hM4D* CNO (-) vs. *βγ-crystallin>hM4D* CNO (-) | ns | 0,3204 |
| | | | *pc2>hM4D* CNO (-) vs. *βγ-crystallin>hM4D* CNO (+) | * | 0,0412 |
| | | | *pc2>hM4D* CNO (-) vs. *pc2>hM4D; βγ-crystallin>hM4D* CNO (-) | ns | 0,6003 |
| | | | *pc2>hM4D* CNO (-) vs. *pc2>hM4D; βγ-crystallin>hM4D* CNO (+) | **** | <0,0001 |
| | | | *pc2>hM4D* CNO (+) vs. *βγ-crystallin>hM4D* CNO (-) | * | 0,0224 |
| | | | *pc2>hM4D* CNO (+) vs. *βγ-crystallin>hM4D* CNO (+) | ns | 0,9976 |
| | | | *pc2>hM4D* CNO (+) vs. *pc2>hM4D; βγ-crystallin>hM4D* CNO (-) | * | 0,0302 |
| | | | *pc2>hM4D* CNO (+) vs. *pc2>hM4D; βγ-crystallin>hM4D* CNO (+) | ns | 0,5482 |
| | | | *βγ-crystallin>hM4D* CNO (-) vs. *βγ-crystallin>hM4D* CNO (+) | * | 0,0328 |
| | | | *βγ-crystallin>hM4D* CNO (-) vs. *pc2>hM4D; βγ-crystallin>hM4D* CNO (-) | ns | 0,8283 |
| | | | *βγ-crystallin>hM4D* CNO CNO (-) vs. *pc2>hM4D; βγ-crystallin>hM4D* CNO (+) | *** | 0,0002 |
| | | | *βγ-crystallin>hM4D* CNO (+) vs. *pc2>hM4D; βγ-crystallin>hM4D* CNO (-) | * | 0,0252 |
| | | | *βγ-crystallin>hM4D* CNO (+) vs. *pc2>hM4D; βγ-crystallin>hM4D* CNO (+) | ns | 0,4734 |
| | | | *pc2>hM4D; βγ-crystallin>hM4D* CNO (-)vs. *pc2>hM4D; βγ-crystallin>hM4D* CNO (+) | **** | <0,0001 |
| | | 240 min | *pc2>hM4D* CNO (-) vs. *pc2>hM4D* CNO (+) | * | 0,0324 |
| | | | *pc2>hM4D* CNO (-) vs. *βγ-crystallin>hM4D* CNO (-) | ns | 0,2781 |
| | | | *pc2>hM4D* CNO (-) vs. *βγ-crystallin>hM4D* CNO (+) | * | 0,0231 |
| | | | *pc2>hM4D* CNO (-) vs. *pc2>hM4D; βγ-crystallin>hM4D* CNO (-) | ns | 0,8020 |
| | | | *pc2>hM4D* CNO (-) vs. *pc2>hM4D; βγ-crystallin>hM4D* CNO (+) | *** | 0,0001 |
| | | | *pc2>hM4D* CNO (+) vs. *βγ-crystallin>hM4D* CNO (-) | * | 0,0264 |
| | | | *pc2>hM4D* CNO (+) vs. *βγ-crystallin>hM4D* CNO (+) | ns | 0,7466 |
| | | | *pc2>hM4D* CNO (+) vs. *pc2>hM4D; βγ-crystallin>hM4D* CNO (-) | * | 0,0247 |

| | | | | | |
|---|---|---|---|---|---|
| | | | *pc2>hM4D* CNO (+) vs. *pc2>hM4D; βγ-crystallin>hM4D* CNO (+) | ns | 0,0944 |
| | | | *βγ-crystallin>hM4D* CNO (-) vs. *βγ-crystallin>hM4D* CNO (+) | * | 0,0178 |
| | | | *βγ-crystallin>hM4D* CNO (-) vs. *pc2>hM4D; βγ-crystallin>hM4D* CNO (-) | ns | 0,4960 |
| | | | *βγ-crystallin>hM4D* CNO CNO (-) vs. *pc2>hM4D; βγ-crystallin>hM4D* CNO (+) | *** | 0,0002 |
| | | | *βγ-crystallin>hM4D* CNO (+) vs. *pc2>hM4D; βγ-crystallin>hM4D* CNO (-) | * | 0,0136 |
| | | | *βγ-crystallin>hM4D* CNO (+) vs. *pc2>hM4D; βγ-crystallin>hM4D* CNO (+) | * | 0,0388 |
| | | | *pc2>hM4D; βγ-crystallin>hM4D* CNO (-)vs. *pc2>hM4D; βγ-crystallin>hM4D* CNO (+) | **** | <0,0001 |
| | | 270 min | *pc2>hM4D* CNO (-) vs. *pc2>hM4D* CNO (+) | * | 0,0312 |
| | | | *pc2>hM4D* CNO (-) vs. *βγ-crystallin>hM4D* CNO (-) | ns | 0,3944 |
| | | | *pc2>hM4D* CNO (-) vs. *βγ-crystallin>hM4D* CNO (+) | * | 0,0130 |
| | | | *pc2>hM4D* CNO (-) vs. *pc2>hM4D; βγ-crystallin>hM4D* CNO (-) | ns | 0,9987 |
| | | | *pc2>hM4D* CNO (-) vs. *pc2>hM4D; βγ-crystallin>hM4D* CNO (+) | **** | <0,0001 |
| | | | *pc2>hM4D* CNO (+) vs. *βγ-crystallin>hM4D* CNO (-) | * | 0,0195 |
| | | | *pc2>hM4D* CNO (+) vs. *βγ-crystallin>hM4D* CNO (+) | ns | 0,6303 |
| | | | *pc2>hM4D* CNO (+) vs. *pc2>hM4D; βγ-crystallin>hM4D* CNO (-) | * | 0,0201 |
| | | | *pc2>hM4D* CNO (+) vs. *pc2>hM4D; βγ-crystallin>hM4D* CNO (+) | ns | 0,1481 |
| | | | *βγ-crystallin>hM4D* CNO (-) vs. *βγ-crystallin>hM4D* CNO (+) | ** | 0,0047 |
| | | | *βγ-crystallin>hM4D* CNO (-) vs. *pc2>hM4D; βγ-crystallin>hM4D* CNO (-) | ns | 0,3221 |
| | | | *βγ-crystallin>hM4D* CNO CNO (-) vs. *pc2>hM4D; βγ-crystallin>hM4D* CNO (+) | **** | <0,0001 |
| | | | *βγ-crystallin>hM4D* CNO (+) vs. *pc2>hM4D; βγ-crystallin>hM4D* CNO (-) | ** | 0,0050 |
| | | | *βγ-crystallin>hM4D* CNO (+) vs. *pc2>hM4D; βγ-crystallin>hM4D* CNO (+) | ** | 0,0050 |
| | | | *pc2>hM4D; βγ-crystallin>hM4D* CNO (-)vs. *pc2>hM4D; βγ-crystallin>hM4D* CNO (+) | **** | <0,0001 |
| | | 300 min | *pc2>hM4D* CNO (-) vs. *pc2>hM4D* CNO (+) | ** | 0,0067 |
| | | | *pc2>hM4D* CNO (-) vs. *βγ-crystallin>hM4D* CNO (-) | ns | 0,1372 |
| | | | *pc2>hM4D* CNO (-) vs. *βγ-crystallin>hM4D* CNO (+) | ** | 0,0024 |
| | | | *pc2>hM4D* CNO (-) vs. *pc2>hM4D; βγ-crystallin>hM4D* CNO (-) | ns | 0,7196 |
| | | | *pc2>hM4D* CNO (-) vs. *pc2>hM4D; βγ-crystallin>hM4D* CNO (+) | **** | <0,0001 |

| | | | | | |
|---|---|---|---|---|---|
| | | | *pc2>hM4D* CNO (+) vs. *βγ-crystallin>hM4D* CNO (-) | ** | 0,0053 |
| | | | *pc2>hM4D* CNO (+) vs. *βγ-crystallin>hM4D* CNO (+) | ns | 0,4537 |
| | | | *pc2>hM4D* CNO (+) vs. *pc2>hM4D; βγ-crystallin>hM4D* CNO (-) | ** | 0,0090 |
| | | | *pc2>hM4D* CNO (+) vs. *pc2>hM4D; βγ-crystallin>hM4D* CNO (+) | ns | 0,2869 |
| | | | *βγ-crystallin>hM4D* CNO (-) vs. *βγ-crystallin>hM4D* CNO (+) | *** | 0,0009 |
| | | | *βγ-crystallin>hM4D* CNO (-) vs. *pc2>hM4D; βγ-crystallin>hM4D* CNO (-) | ns | 0,3348 |
| | | | *βγ-crystallin>hM4D* CNO CNO (-) vs. *pc2>hM4D; βγ-crystallin>hM4D* CNO (+) | **** | <0,0001 |
| | | | *βγ-crystallin>hM4D* CNO (+) vs. *pc2>hM4D; βγ-crystallin>hM4D* CNO (-) | ** | 0,0011 |
| | | | *βγ-crystallin>hM4D* CNO (+) vs. *pc2>hM4D; βγ-crystallin>hM4D* CNO (+) | ** | 0,0028 |
| | | | *pc2>hM4D; βγ-crystallin>hM4D* CNO (-)vs. *pc2>hM4D; βγ-crystallin>hM4D* CNO (+) | **** | <0,0001 |
| | | 330 min | *pc2>hM4D* CNO (-) vs. *pc2>hM4D* CNO (+) | ** | 0,0021 |
| | | | *pc2>hM4D* CNO (-) vs. *βγ-crystallin>hM4D* CNO (-) | ns | 0,1232 |
| | | | *pc2>hM4D* CNO (-) vs. *βγ-crystallin>hM4D* CNO (+) | ** | 0,0021 |
| | | | *pc2>hM4D* CNO (-) vs. *pc2>hM4D; βγ-crystallin>hM4D* CNO (-) | ns | 0,8240 |
| | | | *pc2>hM4D* CNO (-) vs. *pc2>hM4D; βγ-crystallin>hM4D* CNO (+) | **** | <0,0001 |
| | | | *pc2>hM4D* CNO (+) vs. *βγ-crystallin>hM4D* CNO (-) | *** | 0,0003 |
| | | | *pc2>hM4D* CNO (+) vs. *βγ-crystallin>hM4D* CNO (+) | ns | 0,1030 |
| | | | *pc2>hM4D* CNO (+) vs. *pc2>hM4D; βγ-crystallin>hM4D* CNO (-) | *** | 0,0009 |
| | | | *pc2>hM4D* CNO (+) vs. *pc2>hM4D; βγ-crystallin>hM4D* CNO (+) | ns | 0,3507 |
| | | | *βγ-crystallin>hM4D* CNO (-) vs. *βγ-crystallin>hM4D* CNO (+) | *** | 0,0005 |
| | | | *βγ-crystallin>hM4D* CNO (-) vs. *pc2>hM4D; βγ-crystallin>hM4D* CNO (-) | ns | 0,2366 |
| | | | *βγ-crystallin>hM4D* CNO CNO (-) vs. *pc2>hM4D; βγ-crystallin>hM4D* CNO (+) | **** | <0,0001 |
| | | | *βγ-crystallin>hM4D* CNO (+) vs. *pc2>hM4D; βγ-crystallin>hM4D* CNO (-) | *** | 0,0006 |
| | | | *βγ-crystallin>hM4D* CNO (+) vs. *pc2>hM4D; βγ-crystallin>hM4D* CNO (+) | ** | 0,0045 |
| | | | *pc2>hM4D; βγ-crystallin>hM4D* CNO (-)vs. *pc2>hM4D; βγ-crystallin>hM4D* CNO (+) | **** | <0,0001 |
| | | 360 min | *pc2>hM4D* CNO (-) vs. *pc2>hM4D* CNO (+) | *** | 0,0006 |
| | | | *pc2>hM4D* CNO (-) vs. *βγ-crystallin>hM4D* CNO (-) | ns | 0,1435 |

| | | | | | |
|---|---|---|---|---|---|
| | | | *pc2>hM4D* CNO (-) vs. *βγ-crystallin>hM4D* CNO (+) | ** | 0,0015 |
| | | | *pc2>hM4D* CNO (-) vs. *pc2>hM4D; βγ-crystallin>hM4D* CNO (-) | ns | 0,6971 |
| | | | *pc2>hM4D* CNO (-) vs. *pc2>hM4D; βγ-crystallin>hM4D* CNO (+) | **** | <0,0001 |
| | | | *pc2>hM4D* CNO (+) vs. *βγ-crystallin>hM4D* CNO (-) | **** | <0,0001 |
| | | | *pc2>hM4D* CNO (+) vs. *βγ-crystallin>hM4D* CNO (+) | * | 0,0367 |
| | | | *pc2>hM4D* CNO (+) vs. *pc2>hM4D; βγ-crystallin>hM4D* CNO (-) | *** | 0,0002 |
| | | | *pc2>hM4D* CNO (+) vs. *pc2>hM4D; βγ-crystallin>hM4D* CNO (+) | ns | 0,8738 |
| | | | *βγ-crystallin>hM4D* CNO (-) vs. *βγ-crystallin>hM4D* CNO (+) | *** | 0,0006 |
| | | | *βγ-crystallin>hM4D* CNO (-) vs. *pc2>hM4D; βγ-crystallin>hM4D* CNO (-) | ns | 0,2967 |
| | | | *βγ-crystallin>hM4D* CNO CNO (-) vs. *pc2>hM4D; βγ-crystallin>hM4D* CNO (+) | **** | <0,0001 |
| | | | *βγ-crystallin>hM4D* CNO (+) vs. *pc2>hM4D; βγ-crystallin>hM4D* CNO (-) | *** | 0,0003 |
| | | | *βγ-crystallin>hM4D* CNO (+) vs. *pc2>hM4D; βγ-crystallin>hM4D* CNO (+) | * | 0,0211 |
| | | | *pc2>hM4D; βγ-crystallin>hM4D* CNO (-)vs. *pc2>hM4D; βγ-crystallin>hM4D* CNO (+) | **** | <0,0001 |
| | | 390 min | *pc2>hM4D* CNO (-) vs. *pc2>hM4D* CNO (+) | *** | 0,0005 |
| | | | *pc2>hM4D* CNO (-) vs. *βγ-crystallin>hM4D* CNO (-) | ns | 0,3326 |
| | | | *pc2>hM4D* CNO (-) vs. *βγ-crystallin>hM4D* CNO (+) | *** | 0,0008 |
| | | | *pc2>hM4D* CNO (-) vs. *pc2>hM4D; βγ-crystallin>hM4D* CNO (-) | ns | 0,9534 |
| | | | *pc2>hM4D* CNO (-) vs. *pc2>hM4D; βγ-crystallin>hM4D* CNO (+) | **** | <0,0001 |
| | | | *pc2>hM4D* CNO (+) vs. *βγ-crystallin>hM4D* CNO (-) | **** | <0,0001 |
| | | | *pc2>hM4D* CNO (+) vs. *βγ-crystallin>hM4D* CNO (+) | ns | 0,0545 |
| | | | *pc2>hM4D* CNO (+) vs. *pc2>hM4D; βγ-crystallin>hM4D* CNO (-) | **** | <0,0001 |
| | | | *pc2>hM4D* CNO (+) vs. *pc2>hM4D; βγ-crystallin>hM4D* CNO (+) | ns | 0,6559 |
| | | | *βγ-crystallin>hM4D* CNO (-) vs. *βγ-crystallin>hM4D* CNO (+) | **** | <0,0001 |
| | | | *βγ-crystallin>hM4D* CNO (-) vs. *pc2>hM4D; βγ-crystallin>hM4D* CNO (-) | ns | 0,2461 |
| | | | *βγ-crystallin>hM4D* CNO CNO (-) vs. *pc2>hM4D; βγ-crystallin>hM4D* CNO (+) | **** | <0,0001 |
| | | | *βγ-crystallin>hM4D* CNO (+) vs. *pc2>hM4D; βγ-crystallin>hM4D* CNO (-) | **** | <0,0001 |
| | | | *βγ-crystallin>hM4D* CNO (+) vs. *pc2>hM4D; βγ-crystallin>hM4D* CNO (+) | * | 0,0178 |
| | | | *pc2>hM4D; βγ-crystallin>hM4D* CNO (-)vs. *pc2>hM4D; βγ-crystallin>hM4D* CNO (+) | **** | <0,0001 |

| | | | | | |
|---|---|---|---|---|---|
| | | 420 min | *pc2>hM4D* CNO (-) vs. *pc2>hM4D* CNO (+) | *** | 0,0002 |
| | | | *pc2>hM4D* CNO (-) vs. *βγ-crystallin>hM4D* CNO (-) | ns | 0,4376 |
| | | | *pc2>hM4D* CNO (-) vs. *βγ-crystallin>hM4D* CNO (+) | *** | 0,0002 |
| | | | *pc2>hM4D* CNO (-) vs. *pc2>hM4D; βγ-crystallin>hM4D* CNO (-) | ns | >0,9999 |
| | | | *pc2>hM4D* CNO (-) vs. *pc2>hM4D; βγ-crystallin>hM4D* CNO (+) | **** | <0,0001 |
| | | | *pc2>hM4D* CNO (+) vs. *βγ-crystallin>hM4D* CNO (-) | **** | <0,0001 |
| | | | *pc2>hM4D* CNO (+) vs. *βγ-crystallin>hM4D* CNO (+) | ns | 0,0906 |
| | | | *pc2>hM4D* CNO (+) vs. *pc2>hM4D; βγ-crystallin>hM4D* CNO (-) | **** | <0,0001 |
| | | | *pc2>hM4D* CNO (+) vs. *pc2>hM4D; βγ-crystallin>hM4D* CNO (+) | ns | 0,4523 |
| | | | *βγ-crystallin>hM4D* CNO (-) vs. *βγ-crystallin>hM4D* CNO (+) | **** | <0,0001 |
| | | | *βγ-crystallin>hM4D* CNO (-) vs. *pc2>hM4D; βγ-crystallin>hM4D* CNO (-) | ns | 0,1309 |
| | | | *βγ-crystallin>hM4D* CNO CNO (-) vs. *pc2>hM4D; βγ-crystallin>hM4D* CNO (+) | **** | <0,0001 |
| | | | *βγ-crystallin>hM4D* CNO (+) vs. *pc2>hM4D; βγ-crystallin>hM4D* CNO (-) | **** | <0,0001 |
| | | | *βγ-crystallin>hM4D* CNO (+) vs. *pc2>hM4D; βγ-crystallin>hM4D* CNO (+) | * | 0,0112 |
| | | | *pc2>hM4D; βγ-crystallin>hM4D* CNO (-)vs. *pc2>hM4D; βγ-crystallin>hM4D* CNO (+) | **** | <0,0001 |
| | | 450 min | *pc2>hM4D* CNO (-) vs. *pc2>hM4D* CNO (+) | **** | <0,0001 |
| | | | *pc2>hM4D* CNO (-) vs. *βγ-crystallin>hM4D* CNO (-) | ns | 0,5546 |
| | | | *pc2>hM4D* CNO (-) vs. *βγ-crystallin>hM4D* CNO (+) | **** | <0,0001 |
| | | | *pc2>hM4D* CNO (-) vs. *pc2>hM4D; βγ-crystallin>hM4D* CNO (-) | ns | 0,9672 |
| | | | *pc2>hM4D* CNO (-) vs. *pc2>hM4D; βγ-crystallin>hM4D* CNO (+) | **** | <0,0001 |
| | | | *pc2>hM4D* CNO (+) vs. *βγ-crystallin>hM4D* CNO (-) | **** | <0,0001 |
| | | | *pc2>hM4D* CNO (+) vs. *βγ-crystallin>hM4D* CNO (+) | ns | 0,0595 |
| | | | *pc2>hM4D* CNO (+) vs. *pc2>hM4D; βγ-crystallin>hM4D* CNO (-) | **** | <0,0001 |
| | | | *pc2>hM4D* CNO (+) vs. *pc2>hM4D; βγ-crystallin>hM4D* CNO (+) | ns | 0,2626 |
| | | | *βγ-crystallin>hM4D* CNO (-) vs. *βγ-crystallin>hM4D* CNO (+) | **** | <0,0001 |
| | | | *βγ-crystallin>hM4D* CNO (-) vs. *pc2>hM4D; βγ-crystallin>hM4D* CNO (-) | ns | 0,1825 |
| | | | *βγ-crystallin>hM4D* CNO CNO (-) vs. *pc2>hM4D; βγ-crystallin>hM4D* CNO (+) | **** | <0,0001 |
| | | | *βγ-crystallin>hM4D* CNO (+) vs. *pc2>hM4D; βγ-crystallin>hM4D* CNO (-) | **** | <0,0001 |

| | | | | | |
|---|---|---|---|---|---|
| | | | *βγ-crystallin>hM4D* CNO (+) vs. *pc2>hM4D; βγ-crystallin>hM4D* CNO (+) | * | 0,0264 |
| | | | *pc2>hM4D; βγ-crystallin>hM4D* CNO (-)vs. *pc2>hM4D; βγ-crystallin>hM4D* CNO (+) | **** | <0,0001 |
| | | 480 min | *pc2>hM4D* CNO (-) vs. *pc2>hM4D* CNO (+) | **** | <0,0001 |
| | | | *pc2>hM4D* CNO (-) vs. *βγ-crystallin>hM4D* CNO (-) | ns | 0,6113 |
| | | | *pc2>hM4D* CNO (-) vs. *βγ-crystallin>hM4D* CNO (+) | **** | <0,0001 |
| | | | *pc2>hM4D* CNO (-) vs. *pc2>hM4D; βγ-crystallin>hM4D* CNO (-) | ns | 0,5886 |
| | | | *pc2>hM4D* CNO (-) vs. *pc2>hM4D; βγ-crystallin>hM4D* CNO (+) | **** | <0,0001 |
| | | | *pc2>hM4D* CNO (+) vs. *βγ-crystallin>hM4D* CNO (-) | **** | <0,0001 |
| | | | *pc2>hM4D* CNO (+) vs. *βγ-crystallin>hM4D* CNO (+) | ns | 0,2371 |
| | | | *pc2>hM4D* CNO (+) vs. *pc2>hM4D; βγ-crystallin>hM4D* CNO (-) | **** | <0,0001 |
| | | | *pc2>hM4D* CNO (+) vs. *pc2>hM4D; βγ-crystallin>hM4D* CNO (+) | ns | 0,1151 |
| | | | *βγ-crystallin>hM4D* CNO (-) vs. *βγ-crystallin>hM4D* CNO (+) | **** | <0,0001 |
| | | | *βγ-crystallin>hM4D* CNO (-) vs. *pc2>hM4D; βγ-crystallin>hM4D* CNO (-) | ns | 0,0971 |
| | | | *βγ-crystallin>hM4D* CNO CNO (-) vs. *pc2>hM4D; βγ-crystallin>hM4D* CNO (+) | **** | <0,0001 |
| | | | *βγ-crystallin>hM4D* CNO (+) vs. *pc2>hM4D; βγ-crystallin>hM4D* CNO (-) | **** | <0,0001 |
| | | | *βγ-crystallin>hM4D* CNO (+) vs. *pc2>hM4D; βγ-crystallin>hM4D* CNO (+) | * | 0,0392 |
| | | | *pc2>hM4D; βγ-crystallin>hM4D* CNO (-)vs. *pc2>hM4D; βγ-crystallin>hM4D* CNO (+) | **** | <0,0001 |
| | | 510 min | *pc2>hM4D* CNO (-) vs. *pc2>hM4D* CNO (+) | **** | <0,0001 |
| | | | *pc2>hM4D* CNO (-) vs. *βγ-crystallin>hM4D* CNO (-) | ns | 0,8087 |
| | | | *pc2>hM4D* CNO (-) vs. *βγ-crystallin>hM4D* CNO (+) | **** | <0,0001 |
| | | | *pc2>hM4D* CNO (-) vs. *pc2>hM4D; βγ-crystallin>hM4D* CNO (-) | ns | 0,3677 |
| | | | *pc2>hM4D* CNO (-) vs. *pc2>hM4D; βγ-crystallin>hM4D* CNO (+) | **** | <0,0001 |
| | | | *pc2>hM4D* CNO (+) vs. *βγ-crystallin>hM4D* CNO (-) | **** | <0,0001 |
| | | | *pc2>hM4D* CNO (+) vs. *βγ-crystallin>hM4D* CNO (+) | ns | 0,9164 |
| | | | *pc2>hM4D* CNO (+) vs. *pc2>hM4D; βγ-crystallin>hM4D* CNO (-) | **** | <0,0001 |
| | | | *pc2>hM4D* CNO (+) vs. *pc2>hM4D; βγ-crystallin>hM4D* CNO (+) | ns | 0,0800 |
| | | | *βγ-crystallin>hM4D* CNO (-) vs. *βγ-crystallin>hM4D* CNO (+) | **** | <0,0001 |
| | | | *βγ-crystallin>hM4D* CNO (-) vs. *pc2>hM4D; βγ-crystallin>hM4D* CNO (-) | ns | 0,1068 |

| | | | | | |
|---|---|---|---|---|---|
| | | | *βγ-crystallin>hM4D* CNO CNO (-) vs. *pc2>hM4D; βγ-crystallin>hM4D* CNO (+) | **** | <0,0001 |
| | | | *βγ-crystallin>hM4D* CNO (+) vs. *pc2>hM4D; βγ-crystallin>hM4D* CNO (-) | **** | <0,0001 |
| | | | *βγ-crystallin>hM4D* CNO (+) vs. *pc2>hM4D; βγ-crystallin>hM4D* CNO (+) | * | 0,0483 |
| | | | *pc2>hM4D; βγ-crystallin>hM4D* CNO (-)vs. *pc2>hM4D; βγ-crystallin>hM4D* CNO (+) | **** | <0,0001 |
| | | 540 min | *pc2>hM4D* CNO (-) vs. *pc2>hM4D* CNO (+) | **** | <0,0001 |
| | | | *pc2>hM4D* CNO (-) vs. *βγ-crystallin>hM4D* CNO (-) | ns | 0,9286 |
| | | | *pc2>hM4D* CNO (-) vs. *βγ-crystallin>hM4D* CNO (+) | **** | <0,0001 |
| | | | *pc2>hM4D* CNO (-) vs. *pc2>hM4D; βγ-crystallin>hM4D* CNO (-) | ns | 0,2531 |
| | | | *pc2>hM4D* CNO (-) vs. *pc2>hM4D; βγ-crystallin>hM4D* CNO (+) | **** | <0,0001 |
| | | | *pc2>hM4D* CNO (+) vs. *βγ-crystallin>hM4D* CNO (-) | **** | <0,0001 |
| | | | *pc2>hM4D* CNO (+) vs. *βγ-crystallin>hM4D* CNO (+) | ns | 0,9971 |
| | | | *pc2>hM4D* CNO (+) vs. *pc2>hM4D; βγ-crystallin>hM4D* CNO (-) | **** | <0,0001 |
| | | | *pc2>hM4D* CNO (+) vs. *pc2>hM4D; βγ-crystallin>hM4D* CNO (+) | ns | 0,1201 |
| | | | *βγ-crystallin>hM4D* CNO (-) vs. *βγ-crystallin>hM4D* CNO (+) | **** | <0,0001 |
| | | | *βγ-crystallin>hM4D* CNO (-) vs. *pc2>hM4D; βγ-crystallin>hM4D* CNO (-) | ns | 0,3053 |
| | | | *βγ-crystallin>hM4D* CNO CNO (-) vs. *pc2>hM4D; βγ-crystallin>hM4D* CNO (+) | **** | <0,0001 |
| | | | *βγ-crystallin>hM4D* CNO (+) vs. *pc2>hM4D; βγ-crystallin>hM4D* CNO (-) | **** | <0,0001 |
| | | | *βγ-crystallin>hM4D* CNO (+) vs. *pc2>hM4D; βγ-crystallin>hM4D* CNO (+) | ns | 0,1701 |
| | | | *pc2>hM4D; βγ-crystallin>hM4D* CNO (-)vs. *pc2>hM4D; βγ-crystallin>hM4D* CNO (+) | **** | <0,0001 |
| | | 570 min | *pc2>hM4D* CNO (-) vs. *pc2>hM4D* CNO (+) | **** | <0,0001 |
| | | | *pc2>hM4D* CNO (-) vs. *βγ-crystallin>hM4D* CNO (-) | ns | 0,9820 |
| | | | *pc2>hM4D* CNO (-) vs. *βγ-crystallin>hM4D* CNO (+) | **** | <0,0001 |
| | | | *pc2>hM4D* CNO (-) vs. *pc2>hM4D; βγ-crystallin>hM4D* CNO (-) | ns | 0,3592 |
| | | | *pc2>hM4D* CNO (-) vs. *pc2>hM4D; βγ-crystallin>hM4D* CNO (+) | **** | <0,0001 |
| | | | *pc2>hM4D* CNO (+) vs. *βγ-crystallin>hM4D* CNO (-) | **** | <0,0001 |
| | | | *pc2>hM4D* CNO (+) vs. *βγ-crystallin>hM4D* CNO (+) | ns | 0,8632 |
| | | | *pc2>hM4D* CNO (+) vs. *pc2>hM4D; βγ-crystallin>hM4D* CNO (-) | **** | <0,0001 |

| | | | | | |
|---|---|---|---|---|---|
| | | | *pc2>hM4D* CNO (+) vs. *pc2>hM4D; βγ-crystallin>hM4D* CNO (+) | ns | 0,2032 |
| | | | *βγ-crystallin>hM4D* CNO (-) vs. *βγ-crystallin>hM4D* CNO (+) | **** | <0,0001 |
| | | | *βγ-crystallin>hM4D* CNO (-) vs. *pc2>hM4D; βγ-crystallin>hM4D* CNO (-) | ns | 0,5517 |
| | | | *βγ-crystallin>hM4D* CNO CNO (-) vs. *pc2>hM4D; βγ-crystallin>hM4D* CNO (+) | **** | <0,0001 |
| | | | *βγ-crystallin>hM4D* CNO (+) vs. *pc2>hM4D; βγ-crystallin>hM4D* CNO (-) | **** | <0,0001 |
| | | | *βγ-crystallin>hM4D* CNO (+) vs. *pc2>hM4D; βγ-crystallin>hM4D* CNO (+) | ns | 0,3496 |
| | | | *pc2>hM4D; βγ-crystallin>hM4D* CNO (-)vs. *pc2>hM4D; βγ-crystallin>hM4D* CNO (+) | **** | <0,0001 |
| | | 600 min | *pc2>hM4D* CNO (-) vs. *pc2>hM4D* CNO (+) | **** | <0,0001 |
| | | | *pc2>hM4D* CNO (-) vs. *βγ-crystallin>hM4D* CNO (-) | ns | >0,9999 |
| | | | *pc2>hM4D* CNO (-) vs. *βγ-crystallin>hM4D* CNO (+) | **** | <0,0001 |
| | | | *pc2>hM4D* CNO (-) vs. *pc2>hM4D; βγ-crystallin>hM4D* CNO (-) | ns | 0,3241 |
| | | | *pc2>hM4D* CNO (-) vs. *pc2>hM4D; βγ-crystallin>hM4D* CNO (+) | **** | <0,0001 |
| | | | *pc2>hM4D* CNO (+) vs. *βγ-crystallin>hM4D* CNO (-) | **** | <0,0001 |
| | | | *pc2>hM4D* CNO (+) vs. *βγ-crystallin>hM4D* CNO (+) | ns | 0,7865 |
| | | | *pc2>hM4D* CNO (+) vs. *pc2>hM4D; βγ-crystallin>hM4D* CNO (-) | **** | <0,0001 |
| | | | *pc2>hM4D* CNO (+) vs. *pc2>hM4D; βγ-crystallin>hM4D* CNO (+) | ns | 0,3089 |
| | | | *βγ-crystallin>hM4D* CNO (-) vs. *βγ-crystallin>hM4D* CNO (+) | **** | <0,0001 |
| | | | *βγ-crystallin>hM4D* CNO (-) vs. *pc2>hM4D; βγ-crystallin>hM4D* CNO (-) | ns | 0,6672 |
| | | | *βγ-crystallin>hM4D* CNO CNO (-) vs. *pc2>hM4D; βγ-crystallin>hM4D* CNO (+) | **** | <0,0001 |
| | | | *βγ-crystallin>hM4D* CNO (+) vs. *pc2>hM4D; βγ-crystallin>hM4D* CNO (-) | **** | <0,0001 |
| | | | *βγ-crystallin>hM4D* CNO (+) vs. *pc2>hM4D; βγ-crystallin>hM4D* CNO (+) | ns | 0,5261 |
| | | | *pc2>hM4D; βγ-crystallin>hM4D* CNO (-)vs. *pc2>hM4D; βγ-crystallin>hM4D* CNO (+) | **** | <0,0001 |

table S20 Time to reach 50% of settled larvae per treatment.

| Panel | Stimulus | Time at 50% (minutes) |
|---|---|---|
| Fig. 5A | *pc2>hM4D* CNO (-) | 279,1 |
| | *pc2>hM4D* CNO (+) | 284,8 |
| | *βγ-crystallin>hM4D* CNO (-) | 271,3 |
| | *βγ-crystallin>hM4D* CNO (+) | 316,9 |

| | | |
|---|---|---|
| | *pc2>hM4D; βγ-crystallin>hM4D* CNO (-) | 287,3 |
| | *pc2>hM4D; βγ-crystallin>hM4D* CNO (+) | 297,9 |
| Fig. 5B (with 10mM $NH_4Cl$) | *pc2>hM4D* CNO (-) | 202,3 |
| | *pc2>hM4D* CNO (+) | 390,8 |
| | *βγ-crystallin>hM4D* CNO (-) | 226,6 |
| | *βγ-crystallin>hM4D* CNO (+) | 414,8 |
| | *pc2>hM4D; βγ-crystallin>hM4D* CNO (-) | 206,8 |
| | *pc2>hM4D; βγ-crystallin>hM4D* CNO (+) | 257,6 |
| Fig. S6A | *pc2>GFP* CNO (-) | 233,7 |
| | *pc2>GFP* CNO (+) | 250,4 |

table S21 Statistics of tail regression assay for *pc2>hM4D* and *βγ-crystallin>hM4D* in the presence of 10mM $NH_4Cl$

| Figure & Panel | Test | Comparison | P Value summary | P Value |
|---|---|---|---|---|
| Fig. 5D | Kruskal-Wallis Test | | *** | <0,0001 |
| | Dunn's multiple comparisons test | | | |
| | | *pc2>hM4D* CNO (-) vs. *pc2>hM4D* CNO (+) | **** | <0,0001 |
| | | *βγ-crystallin>hM4D* CNO (-) vs *βγ-crystallin>hM4D* CNO (+) | ns | >0,9999 |

table S22 Statistics for Statistics for metamorphosis assay for *pc2>hM4D* and *βγ-crystallin>hM4D* in the presence of 200μM Carvacrol

| Figure & Panel | Test | Comparison | P Value summary | P Value |
|---|---|---|---|---|
| Fig. 5E | Kruskal-Wallis Test | | **** | 0,0002 |
| | Dunn's multiple comparisons test | | | |
| | | *pc2>hM4D* CNO (-) vs. *pc2>hM4D* CNO (+) | *** | 0,0002 |
| | | *βγ-crystallin>hM4D* CNO (-) vs *βγ-crystallin>hM4D* CNO (+) | ns | 0,3104 |

table S23 List of primers

| Primer | Sequence |
|---|---|
| hM3D/hM4D-mCherry -Fw | ggggacaagtttgtacaaaaaagcaggctaaccATGACCTTGCACAATAACAGTAC |
| hM3D/hM4D-mCherry -Rv | ggggaccactttgtacaagaaagctgggtTTACTTGTACAGCTCGTCCATG |
| Promoter Cii-βγ-Crystallin -Fw | ggggacaactttgtatagaaaagttgTACGTCATAATAAACATTTCAATGGG |
| Promoter Cii-βγ-Crystallin -Rv | ggggactgcttttttgtacaaacttgATCAATAATTCAAACGTTAACAAC |
| Promoter Cii-pc2 -Fw | ggggacaactttgtatagaaaagttgCAGCAGTCAAAGGGTTTCTTGAAACAC |
| Promoter Cii-pc2 -Rv | ggggactgcttttttgtacaaacttgGCTGCTTTAAGAATTCTTCGTTTTTTCAC |
| Promoter Cii-Etr1-Fw | ggggacaactttgtatagaaaagttgCGACCACGGAGTTAATTGAAAAC |
| Promoter Cii-Etr1 -Rv | ggggactgcttttttgtacaaacttgTCTGGATAAAGCAATACATACGAG |
| nls::GCaMP6s::nls -Fw | ggggacaagtttgtacaaaaaagcaggc<br>taaccATGGCTAGCCCCAAAAAGAAGAGGAAAGTGGTCGACTCATCACGTCGTAAGTG |
| nls::GCaMP6s::nls -Rv | ggggaccactttgtacaagaaagctggg<br>tTCATACCTTGCGCTTTTTCTTCTTCGCTGTCATCATTTGTACAAACTCTTCGTAG |

| | |
|---|---|
| mKate2 -Fw | ggggacaagtttgtacaaaaaagcaggctaaccATGGTGAGCGAGCTGATTAAGGAGAAC |
| mKate2 -Rv | ggggaccactttgtacaagaaagctgggtTTATCTGTGCCCCAGTTTGCTAG |
| GFP -Fw | ggggacaagtttgtacaaaaaagcaggctaaccATGGCGGATCTGCGAGTACC |
| GFP -Rv | ggggaccactttgtacaagaaagctgggtTTACTTGTACAGCTCGTCCATG |
